# Computationally Driven Discovery and Characterization of SIRT3 Activating Compounds that Fully Recover Catalytic Activity under NAD^+^ Depletion

**DOI:** 10.1101/2023.11.09.566481

**Authors:** Xiangying Guan, Alok Upadhyay, Rama Krishna Dumpati, Sudipto Munshi, Samir Roy, Santu Chall, Ali Rahnamoun, Celina Reverdy, Gauthier Errasti, Thomas Delacroix, Anisha Ghosh, Raj Chakrabarti

## Abstract

Mammalian sirtuins (SIRT1-SIRT7) are a family of nicotinamide adenine dinucleotide (NAD^+^)-dependent protein deacylases that play critical roles in lifespan and age-related diseases. The physiological importance of sirtuins has stimulated intense interest in designing sirtuin activating compounds. However, except for allosteric activators of SIRT1-catalyzed reactions that are limited to specific substrates, methodologies for the rational design of sirtuin activating compounds -- including compounds that activate mitochondrial sirtuins implicated in the age-related decline of cellular metabolism -- have been lacking. Here, we use computational high-throughput screening methodologies and a biophysical model for activation of the major mitochondrial sirtuin SIRT3 to identify novel small molecule activators of the human SIRT3 enzyme from a 1.2 million compound library. Unlike previously reported SIRT3 activators like Honokiol, which only transiently upregulate SIRT3 under non-steady state conditions and reduce the steady state catalytic efficiency of the enzyme, several of the novel compounds identified here are potent SIRT3 activators in both the steady and non-steady states. Two such compounds can almost double the catalytic efficiency of the enzyme with respect to NAD^+^, which would be sufficient to almost entirely compensate for the loss in SIRT3 activity that occurs due to the reduction in mitochondrial coenzyme concentration associated with aging, and display AC50s (concentrations of half-maximal activation) as low as 100 nM. The current work thus reports first-in-class, non-allosteric steady state activators that activate SIRT3 through a novel, mechanism-based mode of activation and that may be developed further for therapeutic applications.

## INTRODUCTION

Sirtuins are NAD^+^-dependent deacylation enzymes that have emerged as critical regulators of many cellular pathways including age-related diseases (*1*). They depend on the cofactor NAD^+^ to cleave acyl groups from lysine side chains of substrate proteins, producing nicotinamide (NAM) as a by-product. Their mechanism of action is depicted in **Fig. S1** (*2–7*).

The search for pharmacological agents that activate sirtuins has become a focus for many anti-aging studies due to the lifespan extending effects of sirtuin upregulation in mammals (*8, 9*). Almost all reported sirtuin activators target SIRT1 through allosteric activation (*9–13*) that reduces the dissociation constant of the substrate (*K*_*d*,*Ac*−*Pr*_, the acylated protein dissociation constant) to activate the enzyme. Allosteric activators of SIRT1 bind outside the active site to an allosteric domain that is not shared by SIRT2-7 (*11*). Moreover, allosteric activators only work with a limited set of SIRT1 substrates (*14–18*).

Small molecules that can activate sirtuins through non-allosteric modes of action are of particular interest, because sirtuin activity often declines during aging for reasons other than substrate depletion. In particular, a hallmark of aging is a significant reduction in cellular NAD^+^ levels, which can fall below 50% of the basal cofactor level in youth (*19*). Unlike allosteric activators such as resveratrol, which are specific to SIRT1 and have not been applied successfully to other sirtuins (*11*), NAD^+^ supplementation (*20*) can activate most mammalian sirtuins in a substrate-independent fashion. On the other hand, NAD^+^ is a central coenzyme in metabolism (*21*) so its supplementation can result in nonspecific side effects, and moreover it must be delivered in the form of cell-permeable precursors whose conversion to the active cofactor depends on cellular biosynthetic pathways that differ between cellular subcompartments (e.g., mitochondria, cytoplasm) (*22*).

A preferred strategy for activation of sirtuins (**Fig. S1**) is to lower the *K_m_* for NAD^+^ (*K*_*m*,*NAD*^+^_). Such a reduction in *K*_*m*,*NAD*^+^_ would activate sirtuins similarly to NAD^+^ supplementation, but would be specific to sirtuins, would not depend on cellular biosynthetic pathways, and may potentially enable isoform-specific sirtuin activation. In contrast to allosteric activation that reduces *K*_*d*,*Ac*−*Pr*_, this approach could be applicable to sirtuins other than SIRT1 as well as more diverse substrates. Moreover, since NAD^+^ is directly involved in the sirtuin ADP ribosylation reaction, regulation of *K*_*m*,*NAD*^+^_ may theoretically be achievable by methods including, but not limited to, varying the binding affinity of NAD^+^.

A few compounds have been reported in the literature as being activators of sirtuins other than SIRT1 (*23–26*). Concurrently, sirtuins whose preferred substrates are long-chain acylated peptides were reported to have their activities on shorter chain acyl groups enhanced in the presence of certain fatty acids, further corroborating the feasibility of substrate-specific sirtuin activation (*27, 28*). However, these studies have not shed light on how the underlying mechanism by which such activation may occur can be used to guide the design of novel sirtuin activating compounds.

In our recent work (*29*), we presented a biophysical theory for activation of sirtuins that is distinct from any of the known modes of enzyme activation (*10*), and that does not rely on binding to an allosteric pocket. We introduced a steady state model of sirtuin-catalyzed deacylation reactions in the presence of NAD^+^ cofactor and endogenous inhibitor NAM, and then quantitatively established how the catalytic efficiency *k*_*cat*_/*K*_*m*,*NAD*^+^_ can be altered by small molecules, determining their required biophysical properties to function as non-allosteric activators. This theoretical framework, termed mechanism-based enzyme activation, has been further generalized to include non-steady state activation, and several novel compounds that activate sirtuins other than SIRT1 under non-steady state conditions were recently identified from DNA-encoded libraries (DEL) and characterized using this theory (*30, 31*).

In this paper, computational high-throughput screening methodologies were utilized to identify novel small molecule activators of the human SIRT3 enzyme. SIRT3 was chosen as the target because it is the major mitochondrial sirtuin (*32*) and because the mitochondrial theory of aging (*33*) indicates that metabolic decline in mitochondria is an important driver of age-related disorders. To develop a rational design methodology for SIRT3 activators, we first applied the theory of mechanism-based enzyme activation to experimentally and computationally characterize the effects of the compound Honokiol (HKL) that has been previously reported as a SIRT3 activator and shown to reverse age-related cardiac hypertrophy in mice (*24*). We were able to demonstrate that, through modulation of local active site conformational degrees of freedom, Honokiol activates SIRT3 under non-steady state conditions, while having an inhibitory effect in the steady state that increases *K*_*m*,*NAD*^+^_. Elucidation of the mechanism whereby Honokiol modulates SIRT3 under the above conditions allowed us to identify potential ligand-receptor interactions that can be used as markers to identify compounds with Honokiol-like activity, which could be developed into steady state SIRT3 activator lead compounds with accordingly improved potency and pharmacodynamics.

To identify novel SIRT3 modulators using our integrated experimental/computational framework, a 1.2 million compound library was screened leveraging our knowledge of the binding mode and activation mechanism of Honokiol. Unlike previously reported activators like Honokiol and those identified from DEL libraries in our prior work (*31*), several of the novel hits identified are SIRT3 activators in both the steady and non-steady states, nearly doubling catalytic efficiency primarily through *K*_*m*,*NAD*^+^_ reduction. Since these compounds activate SIRT3 under a wider range of physiologically relevant conditions than previously reported non-steady state activators like Honokiol and display nanomolar AC50s (concentration of half-maximal activation), they constitute an ideal starting point for further development of SIRT3 activating drug-like compounds. The current work thus provides what are first-in-class steady state activators of SIRT3, which operate through a new non-allosteric, mechanism-based mode of activation.

## RESULTS

### Structural and computational characterization of SIRT3 activation by HKL and virtual screening for novel SIRT3 activators

Our earlier work (*29–31*) established that non-allosteric, mechanism-based activators can modulate the distributions of local degrees of freedom for residues in the flexible loop whose conformations change during the reaction, resulting in tradeoffs of the ΔΔG’s for various reaction steps upon stabilization of one such conformation. This is a critical distinction from allosteric activation, where the conformational changes stabilized by the modulator are not specifically associated with certain steps in the enzymatic reaction.

Activators of sirtuin enzymes that do not possess a known allosteric site (*34, 35*) have been shown through crystallographic studies to induce conformational changes in the so-called flexible cofactor binding loop in these enzymes. The conformation of this loop changes after the first chemical step of the reaction (cleavage of nicotinamide from the NAD^+^ cofactor). SIRT3, the mitochondrial sirtuin, is one of the most important sirtuins implicated in regulating healthspan (*36*); hence we chose it as the subject of study. **Fig. 1** presents a structural alignment that depicts the conformational changes occurring in this loop during the catalytic cycle of SIRT3. Honokiol has been reported as a SIRT3 activator for the MnSOD protein substrate (*24*) and since SIRT3 does not possess a known allosteric site, we studied HKL as a potential mechanism-based activator of SIRT3. Honokiol was co-crystallized with the SIRT3 ternary reactants complex, resulting in loop closure akin to that which occurs following the first step of the chemical reaction (**Fig. 1**). Molecular dynamics simulations were employed to study the flexibility of the SIRT3 cofactor binding loop in different conformations associated with the various stages of the sirtuin catalytic cycle (**Fig. 2**). The high B-factors for residues of cofactor binding loop in experimentally obtained structures of ternary complex are consistent with the potential ability of modulator binding near the loop to alter its conformation.

**Fig. 1.**
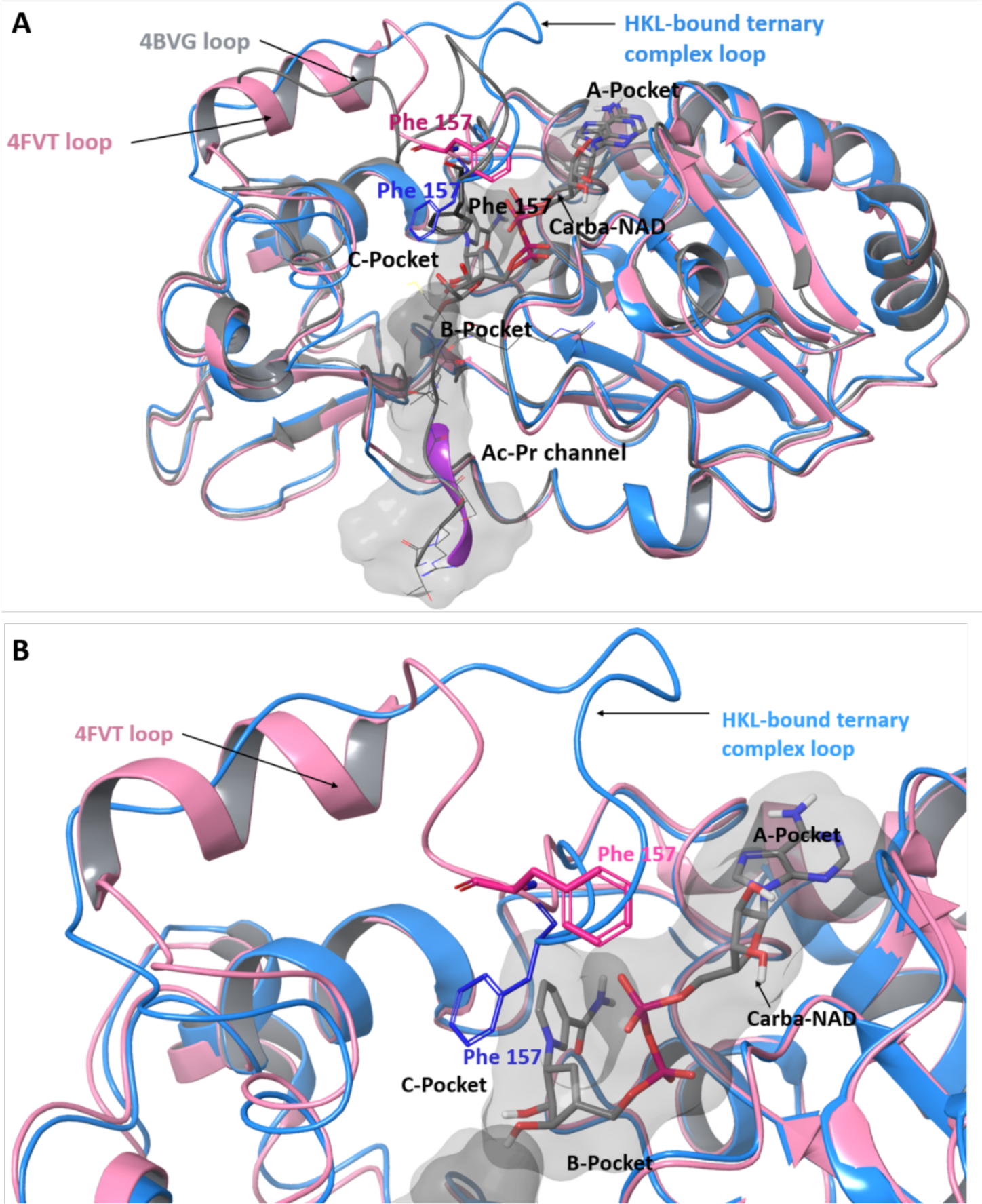
Structural alignment that depicts the conformational changes occurring in the loop during the catalytic cycle of SIRT3. **(A)** Superposition of SIRT3 ternary complex 4FVT (Pink) with SIRT3 ternary complex in the presence of Honokiol (HKL-bound ternary complex; HKL not shown) (Blue) and SIRT3 alkylimidate intermediate 4BVG (grey). The figure highlights the loop regions of each structure and their position relative to internal cavities occupied by an unreactive substrate analogue for NAD-consuming enzymes (Carba-NAD, in stick representation) and peptide (channel highlighted in purple, acetyl-CoA synthetase 2 (ACS2) peptide in grey lines). The loop region, Ile 154-Tyr 175, in HKL-bound ternary complex, 4BVG depicted in gray is largely disordered random-coil, while its equivalent in 4FVT (pink) has a significant amount of α-helical structure. The co-crystallized ligand, Carba-NAD, occupies a significant portion of Pockets A and C in the catalytically active conformation, with its nicotinamide (NAM) moiety in the C pocket rendering loop closure energetically unfavorable. ADP ribosylation of the acetylated peptide substrate and associated NAM dissociation results in loop closure and Phe 157 entering the C pocket (4BVG). In HKL-bound ternary complex structure, crystallized in the presence of Honokiol (with Honokiol unresolved possibly due to loop disorder), the loop adopts a conformation similar to that in 4BVG despite the presence of Carba-NAD, but with Phe 157 slightly less buried in the C pocket due to the presence of nicotinamide there. Loop residues 164 to 171 in HKL-bound ternary complex structure, which were unresolved due to flexibility, were built *ab initio*. **(B)** Close-up showing only the 4FVT and HKL-bound ternary complex structures.

**Fig. 2.**
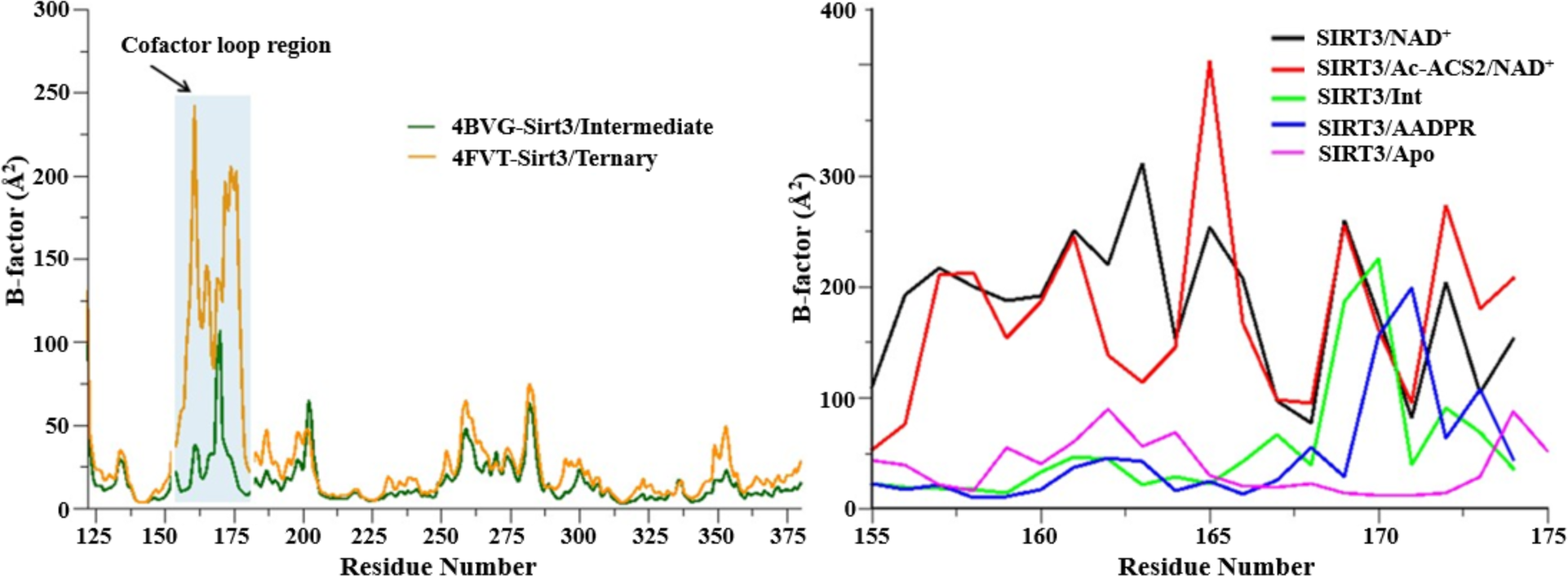
Plot showing simulated B-factor values for C_α_ atoms belonging to (A) residues 125-375; (B) the co-factor binding loop region of various SIRT3 complexes (catalytic cycle). Residues (162–170) are known to adopt α helix conformation when bound to substrate. Ternary = Ac-ACS2 (acetyl-CoA synthetase 2)/NAD^+^ complex, Int = alkylimidate intermediate, AADPR =O-Acetyl-Adenosine-Diphosphate-Ribose.

Co-crystallization of HKL with the human SIRT3 reactants complex produced two crystal structures: a) a 1.5Å structure including acetylated p53-AMC peptide but not Carba-NAD (HKL-SIRT3-1.5Å structure, HKL:SIRT3:AcPr), and b) a 2.4Å structure including acetylated p53-AMC peptide and Carba-NAD (HKL-SIRT3-2.4Å structure, HKL:SIRT3:AcPr:Carba-NAD) (**Fig. 3**). Due to the conformational flexibility of the loop, a few residues were unresolved. In both structures, HKL induces loop closure. In the absence of Carba-NAD (**Fig. 3A**), HKL binds within the A pocket where Carba-NAD would otherwise be located and engages in hydrogen bonding interactions with the backbone atoms of Glu 323, and hydrophobic interactions with FDL. In the presence of Carba-NAD (**Fig. 3C**), HKL binds within a partially exposed pocket above the active site and forms hydrogen bond (H-bond) interactions with Arg 158, and vdW interactions with Phe 237, Leu 322, and Glu 323.

**Fig. 3.**
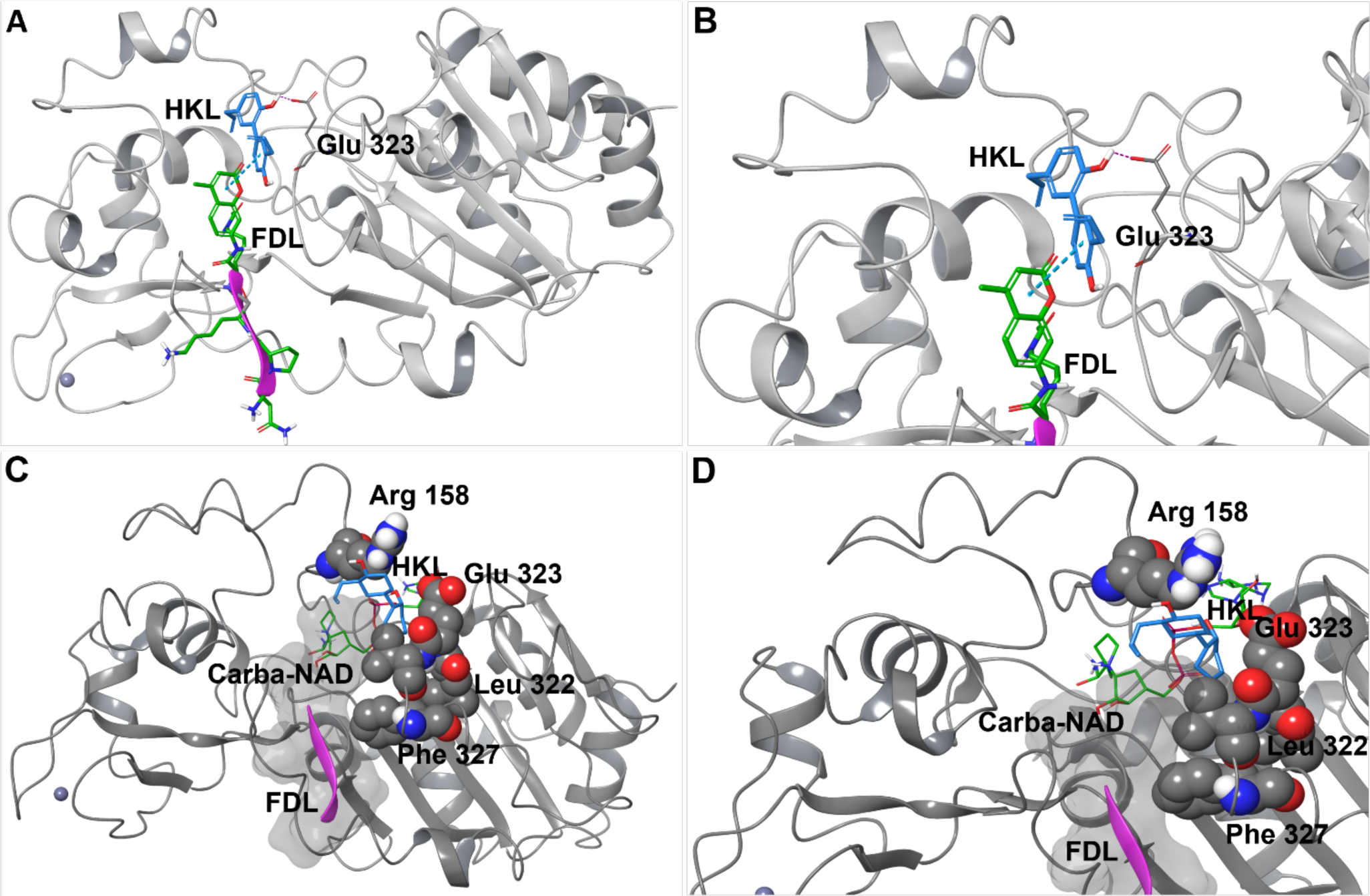
Crystallographic structures of SIRT3 receptor structures co-crystallized with Honokiol without (A, B) and with (C, D) Carba-NAD. Color scheme is as follows: p53-AMC acetylated peptide (FDL) = Pink tube with green sticks, HKL = blue, Carba-NAD = green. **(A)** In the absence of Carba-NAD (HKL-SIRT3-1.5Å structure), HKL binds in an internal pocket overlapping with the binding pocket of NAD^+^. **(B)** Close-up of the binding site of HKL provides another view of the HKL binding interactions, including hydrogen bonding interactions with backbone atoms of Glu 323, and hydrophobic interactions with FDL. **(C)** In the presence of Carba-NAD (HKL-SIRT3-2.4Å structure), HKL binds in an internal pocket overlapping with the binding pocket of NAD^+^. **(D)** Close-up of the binding site provides another view of HKL forming H-bond interactions with Arg 158, vdW interactions with Phe 327, Leu 322, and Glu 323 (shown in space-filling representation) stabilizing a closed loop conformation despite the presence of the nicotinamide moiety of Carba-NAD in the C pocket.

Molecular dynamics simulations for two different conformations of the cofactor binding loop in the ternary reactants complex – the open and closed loop conformations – were also carried out. Comparison of interactions between the loop and NAD^+^ based on these simulations are shown in **Fig. 4**. Whereas cationic residues including Arg 158 engage in favorable electrostatic interactions with NAD^+^ upon loop closure, Phe 157 induces strain in the nicotinamide moiety of NAD^+^, which typically prevents loop closure in the absence of a modulator.

**Fig. 4.**
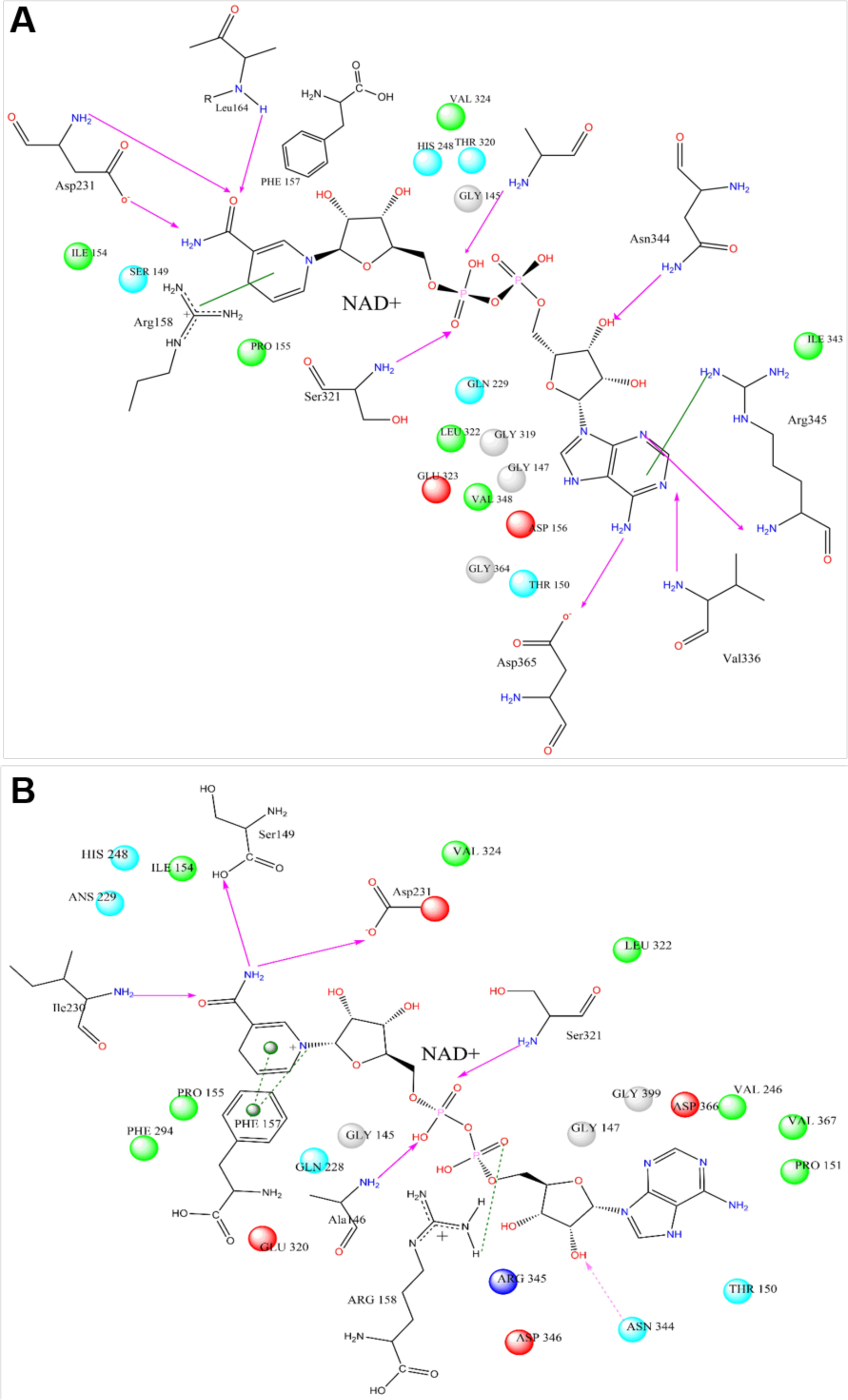
NAD^+^ ligand interaction diagrams for the SIRT ternary reactants complex with (A) open and (B) closed loop conformations depicting the significant changes in the residue interactions with NAD^+^ upon closing of the loop induced by modulators. The geometries depicted are MD averages.

To investigate the effect of HKL-induced changes of the cofactor binding loop conformation on coproduct binding affinity, we calculated AADPR (O-Acetyl-ADP-Ribose, a metabolite produced from NAD^+^ as a product of sirtuin-mediated protein deacetylation) binding affinities from molecular dynamics simulations for open and closed conformations of the loop (**Fig. 5**). The significantly higher binding energies for the closed loop conformation demonstrate that stabilization of the closed loop conformation will increase the co-product binding affinity and reduce the rate of co-product dissociation.

**Fig. 5.**
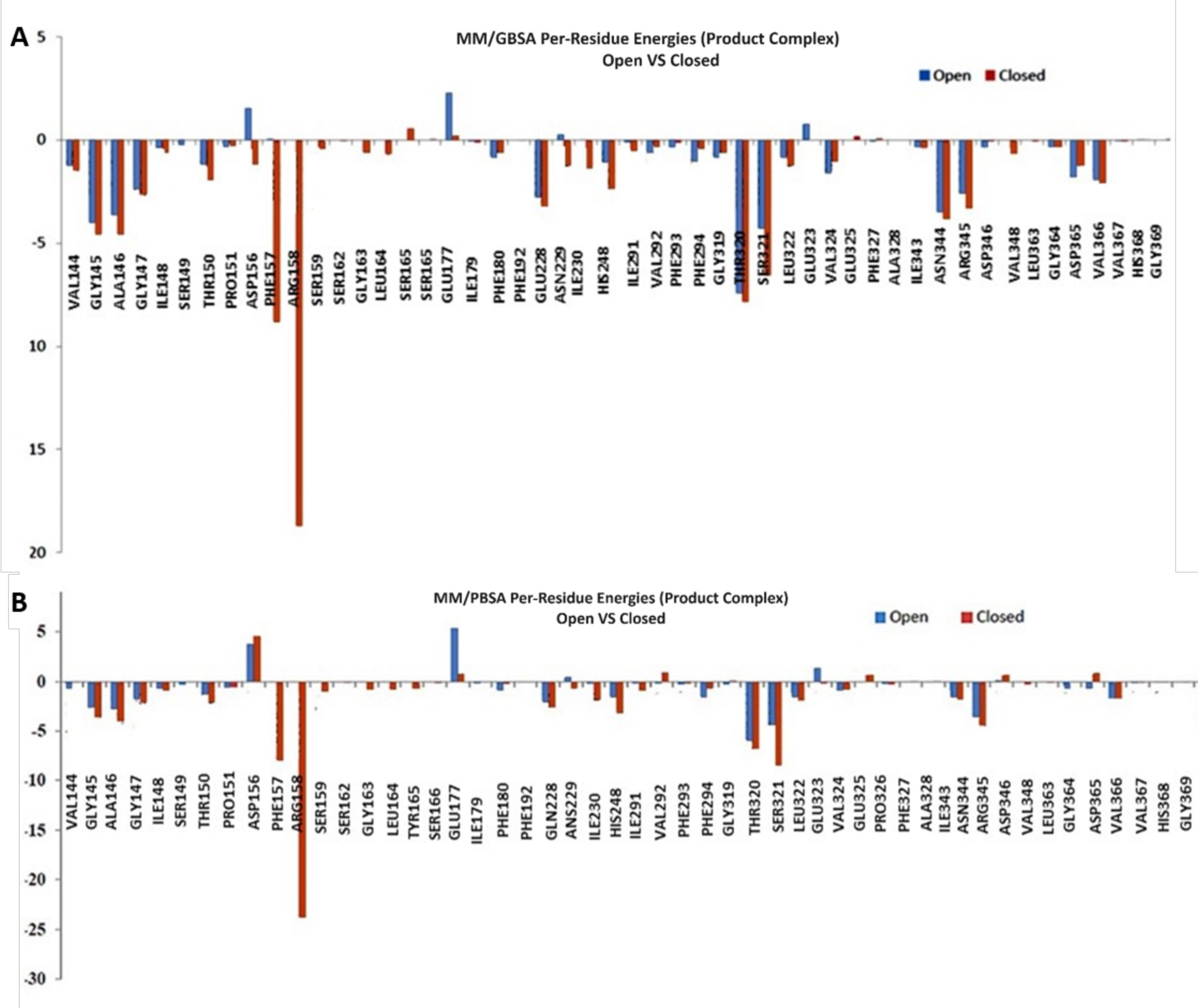
By-residue (A) MM-GBSA and (B) MM-PBSA binding energies of SIRT3 to the AADPR coproduct for open vs closed loop conformations. The red bars show significant stabilizing interactions that increase AADPR binding affinity are generated upon loop closure.

Finally, to assess the model fit to the electron density, we obtained its RSCC and RSR values from the wwPDB (*37*) validation report. These statistics, reported in **Table S1**, are indicative of good fit.

### Virtual screening for non-allosteric activators of SIRT3

Building on the above results, we next applied high-throughput computational/experimental screening methodologies to identify novel SIRT3 activators. Amazon cloud servers were used for virtual screening (VS) of a 1.2 million compound database from ChemBridge using AutoDock Vina (*38, 39*). Two QM-MM geometry optimized SIRT3 complexes (4FVT and 4BVG) were used as receptors in this docking exercise. Hits were identified based on their docking scores (in AutoDock Vina and MOE (*40*)) for these two receptors as well as their H-bond interactions with specific residues lining the binding site, including those in the cofactor binding loop. Honokiol was included as a positive control in the library and subsequently identified as a hit compound according to these criteria. Selected compounds, out of the top one thousand as ranked by AutoDock Vina, were chosen based on their docking scores (between −10.6 to −7.0), best coverage of scaffold diversity, lack of repetition, and maximum number of favorable interactions with residues at the putative binding sites. These compounds were subjected to enzymatic activity assessment (% Control) to confirm binding and modulation (**Tables 1 and S2**) prior to more detailed characterization. Specific residues displaying H-bonds with the top docked poses of these compounds and their degrees of receptor buriedness are reported in **Tables 1** and **S2**. **Scheme 1** illustrates the virtual screening workflow.

**Scheme 1.**
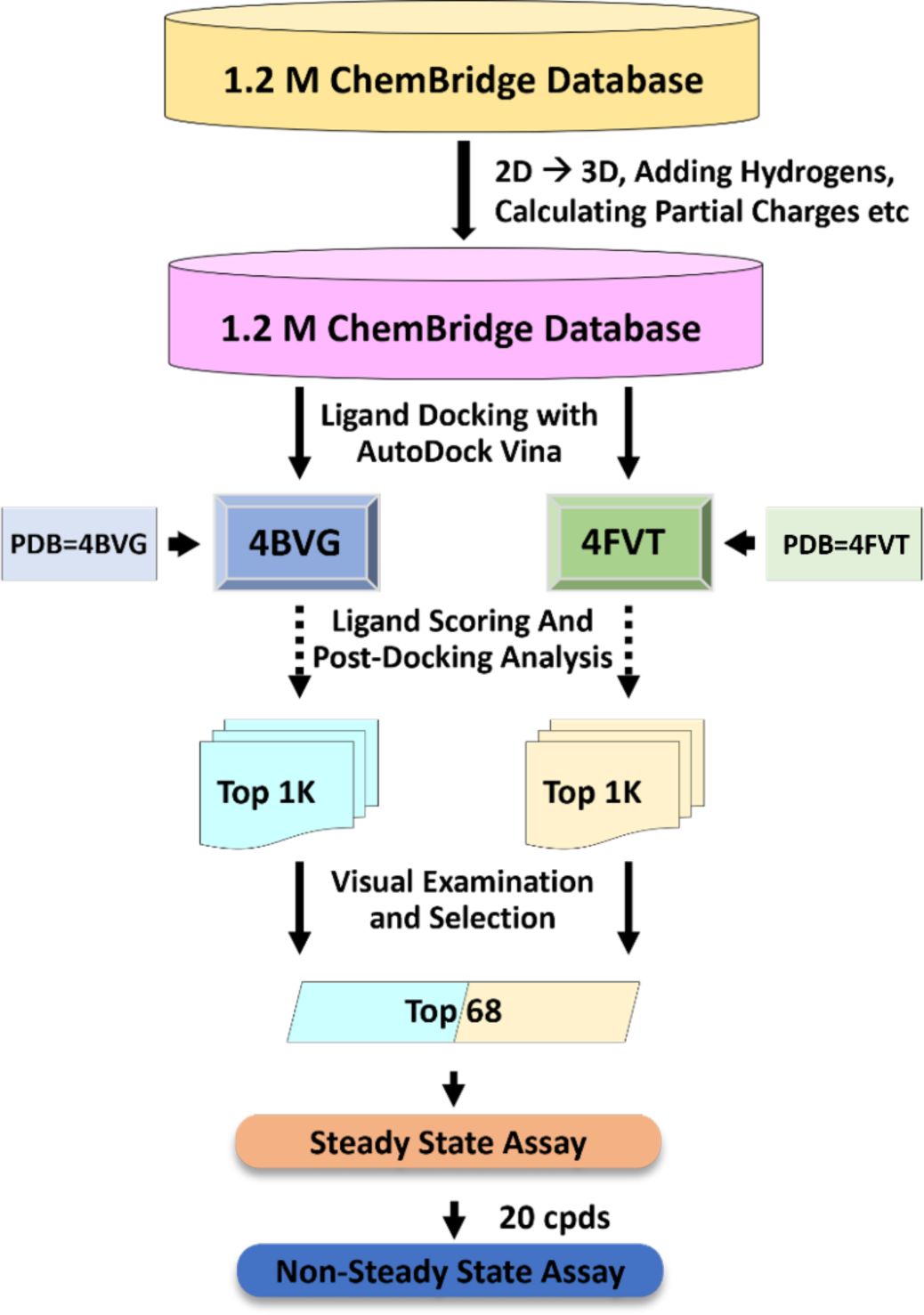
The virtual screening process used in this study. Schematic representation of virtual high-throughput screening (VHTS) for non-allosteric modulators of SIRT3. A database containing 1.2M compounds was subjected to a series of operations which started with conversion from 2D to 3D and ended with a quick energy minimization for each compound. These compounds were then, independently, docked with two SIRT3 receptor structures (4BVG and 4FVT). The best poses for the top 1,000 compounds (by calculated binding energy) in each docking run were chosen for further post-docking analysis - which was performed using MOE as described in post-docking analysis below. The final step involved careful examination and selection of 68 compounds, from both docking runs, which gave us the best coverage of scaffold diversity, lack of repetition and maximum number of favorable interactions with residues at the putative binding site. A total of 68 compounds were tested under steady state conditions and 20 of these that were identified as steady state inhibitors were then tested under non-steady state conditions.

To assess the overall performance of our computational-experimental methodology, **Fig. 6** plots the ROC curve of the FPR (False Positive Rate) against the TPR (True Positive Rate). True positives are defined as compounds with experimentally observed modulation of SIRT3 that also exhibit favorable docking scores, while false positives are defined as compounds with experimentally observed non-modulation of SIRT3 which nonetheless exhibit favorable docking scores. The ROC plot was generated based on the docking scores on 4BVG and 4FVT receptors for about 1.2M docked compounds. The modulation effect on SIRT3 of 68 compounds that comprised a top-ranked subset of 1.2M compounds was determined experimentally with the HPLC assay. These experimental data were used to generate probability distributions of positives for specified ranges of the compound docking scores for the full set of 1.2M compounds (details in Materials and Methods). With an AUC of 0.89, the extrapolated 1.2M ROC curve indicates that our methodology is successful in identifying novel SIRT3 modulators.

**Fig. 6.**
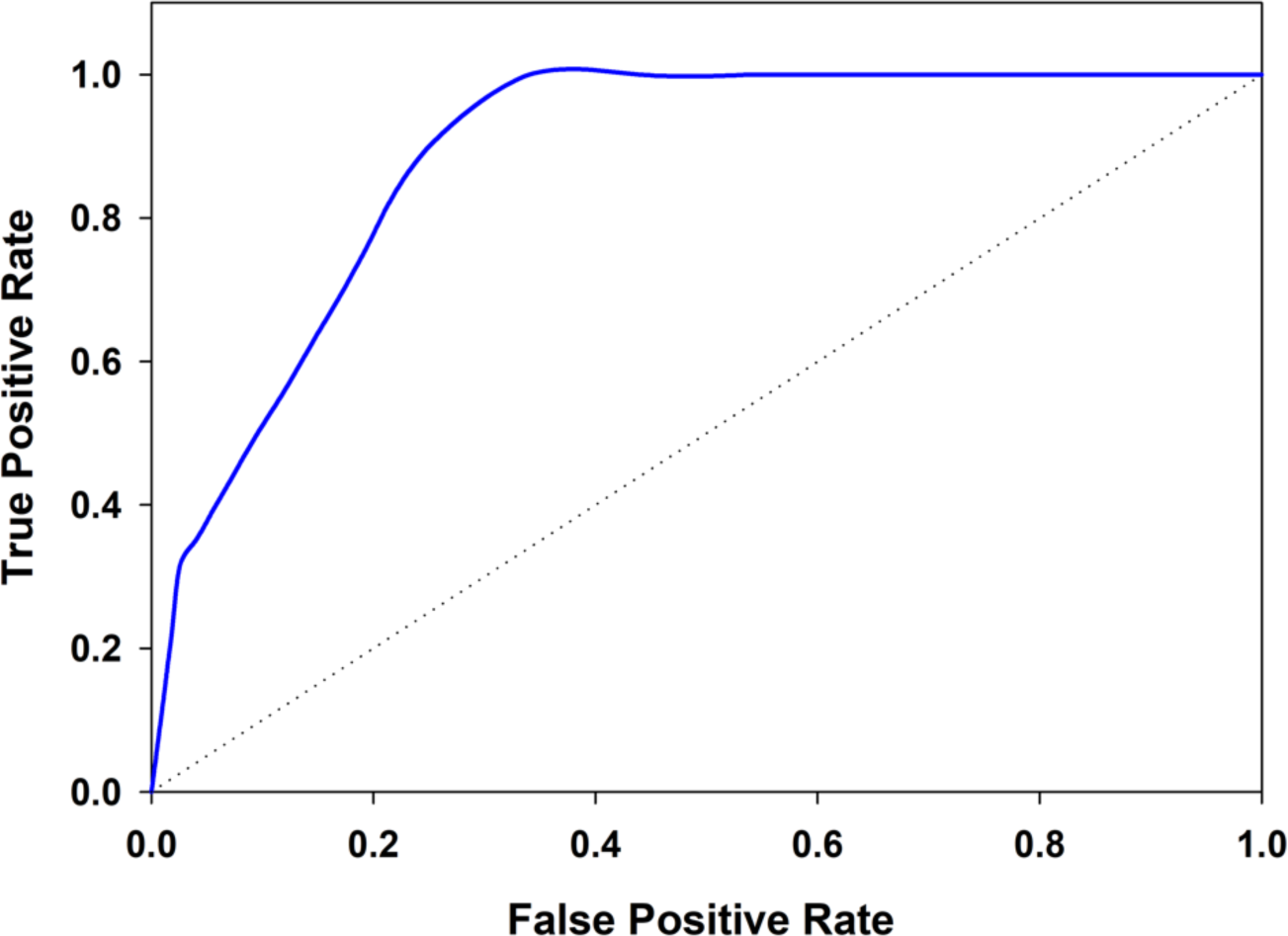
ROC curve for screened compounds generated using the docking scores on 4FVT and 4BVG receptors. The modulation effect values used for calculating the False Positive Rate (FPR) and True Positive Rate (TPR) for the full set of 1.2M compounds were sampled, for specified ranges of the docking score, from probability distributions obtained from the positive rates of the subset of 68 compounds for which modulation was experimentally measured (Materials and Methods). The ROC curve shown with AUC value of 0.89 corresponds to the best docking score across the 4BVG and 4FVT receptors.

### Comparison of binding modes of top VS hit compounds to HKL

The H-bond interaction pattern of the residues in the binding site, especially those in the flexible loop, with the top VS hit compounds were analyzed and are shown in **Table 1**. H-bond interactions with Pro 155, Asp 156, Phe 157, Arg 158, Glu 177, Glu 198, Tyr 204, Asn 229, and Glu 323 are the most common among hits with good experimental activity. A few of these compounds exhibit a similar pattern of H-bond interaction with SIRT3 as HKL. The binding modes of two VS hit compounds (the non-steady state activator compound CAT_S30 and the steady state activator compound CAT_S35) in two receptor models of SIRT3 (SIRT3 ternary complex (4FVT) and HKL-SIRT3 ternary complex (HKL-SIRT3-2.4Å)) are depicted in **Fig. 7**. Steady state activator CAT_S35 was found to interact with 4FVT and HKL-SIRT3 ternary complex structures via 0 and 1 H-bonds respectively, whereas one H-bond interaction with Arg 158 was found between non-steady state activator CAT_S30 and 4FVT but 3 H-bonds were detected with the HKL-SIRT3 ternary complex receptor (**Fig. 7**).

**Fig. 7.**
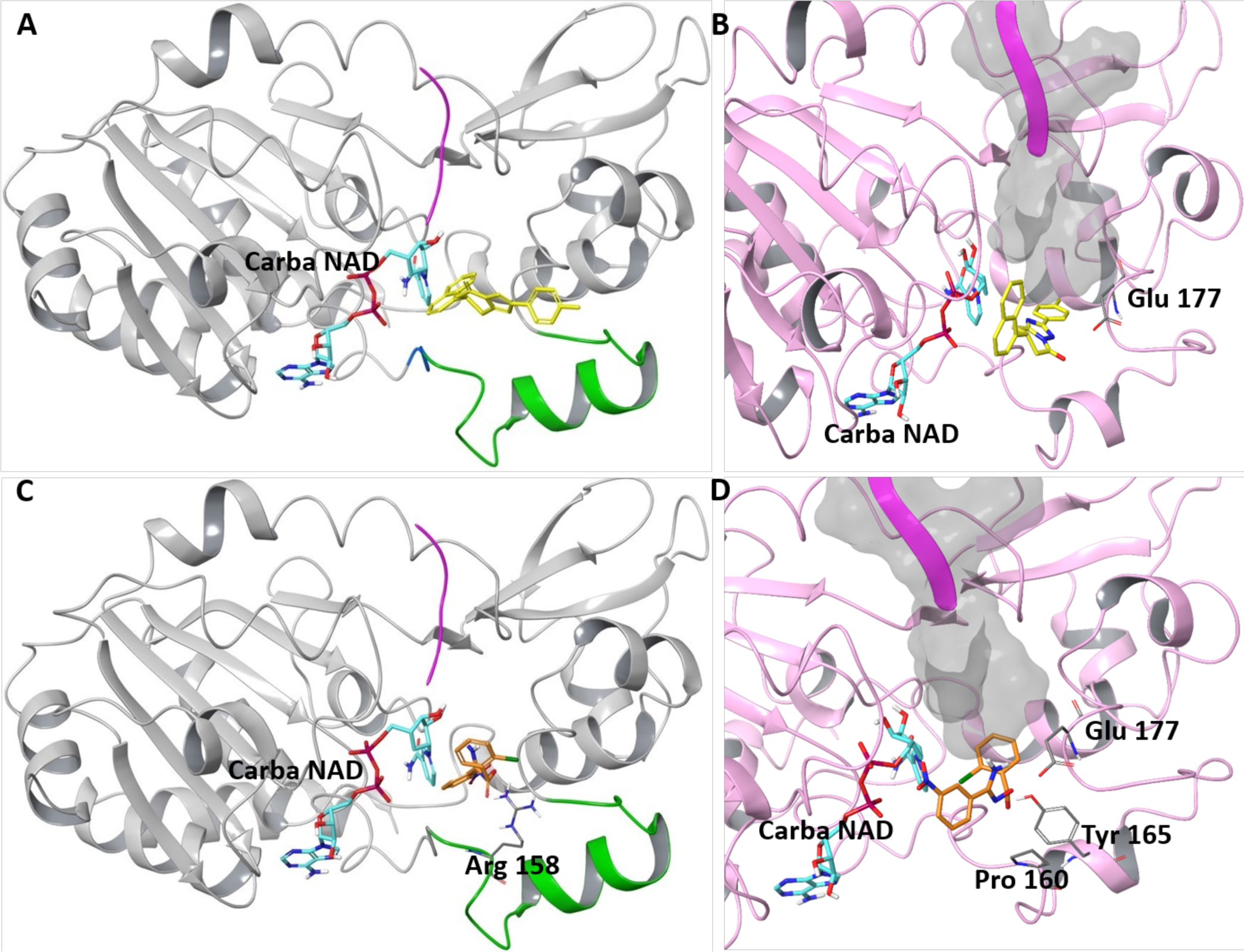
Top docked poses of two selected activator hits in both receptor models of SIRT3. **(A)** In 4FVT, the top non-steady state activator hit compound, Compound ID = CAT_S30, exhibits a score of – 9.8, but forms no H-bonds with residues lining the site. (**B**) In HKL-SIRT3-2.4Å structure, the same compound has Vina score of −6.9 and forms H-bond interaction with Glu 177. Color scheme in A and B is as follows: loop = green, top pose of CAT_S30 = yellow, Carba-NAD = cyan. **(C)** Top pose of best steady state activator hit, Compound ID = CAT_S35, has Vina score of −8.4 in 4FVT and forms H-bonds with Arg 158. **(D)** In HKL-SIRT3-2.4Å structure, it has a score of - 6.7, and forms H-bond with Pro 160, Tyr 165, and Glu 177. Color scheme in **C** and **D** is as same as in **A** and **B**, except for the top pose of CAT_S35 = orange.

**Table 1.** Docking scores of, and in vitro SIRT3 modulation by, the top activator compounds selected through virtual screening. 1.2 million compound library was docked against two SIRT3 complexes (4FVT and 4BVG) using AutoDock Vina. The docking scores, H-bond interaction, and Rec_Buried scores are presented. The modulation effects of top hits were validated and % activities are reported ([cpd]=10 μM). Blue: non-steady state activator; green: steady state activator.

| Compound ID | Docking score | H-bond | Rec_Buried | % Control |  |
| --- | --- | --- | --- | --- | --- |
|  |  |  |  | Steady-state | Non-Steady state |
| CAT_S4 | 4FVT: -9.7 | 4FVT: E177 | 4FVT: 0.82 | 65.8 | 103.0 |
| CAT_S32 | 4FVT: -9.6 | 4FVT: R158 | 4FVT: 0.87 | 52.7 | 103.1 |
|  | 4BVG: -9.8 | 4BVG: None | 4BVG: 0.85 |  |  |
| CAT_S30 | 4FVT: -9.8 | 4FVT: None | 4FVT: 0.84 | 91.3±1.5 | 114.0±3.6 |
|  | 4BVG: -9.6 | 4BVG: P155, Y204 | 4BVG: 0.87 |  |  |
| CAT_S21 | 4BVG: -9.9 | 4BVG: P155, I230, D231 | 4BVG: 0.87 | 96.9±1.1 | 106.1±0.6 |
| CAT_S19 | 4BVG: -9.9 | 4BVG: Y204, I230, D231 | 4BVG: 0.95 | 98.2 | 103.1 |
| CAT_S25 | 4BVG: -9.6 | 4BVG: P155, Y204, I230 | 4BVG: 0.88 | 86.8±1.6 | 104.2±1.7 |
| CAT_S1 | 4FVT: -9.9 | 4FVT: R158, E177 | 4FVT: 0.77 | 92.6±2.9 | 103.3±1.5 |
| CAT_S35 | 4FVT: -7.1 | 4FVT: R158 | 4FVT: 0.71 | 115.2±4.5 | NA |
|  | 4BVG: -8.4 | 4BVG: E198 | 4BVG: 0.36 |  |  |
| CAT_S26 | 4FVT: -8.1 | 4FVT: F157, R158 | 4FVT: NA | 113.6±6.4 | NA |
|  | 4BVG: -8.7 | 4BVG: N167, S162, Y204 | 4BVG: 0.76 |  |  |
| CAT_S14 | 4BVG: -9.8 | 4BVG: P155, N229, Y204 | 4BVG: 0.86 | 103.1±1.4 | 112.6±4.1 |
| CAT_S2 | 4FVT: -10.1 | 4FVT: R158, E323 | 4FVT: 0.90 | 106.7±1.7 | 108.1±1.6 |
| CAT_S53 | 4FVT: -7.1 | 4FVT: P155, S159, D323 | 4FVT: 0.45 | 113.6±3.6 | NA |
|  | 4BVG: -7.4 | 4BVG: E198, Q228 | 4BVG: 0.80 |  |  |
| CAT_S59 | 4FVT: -7.6 | 4FVT: E177 | 4FVT: 0.52 | 102.7±2.9 | NA |
|  | 4BVG: -7.0 | 4BVG: E198 | 4BVG: 0.68 |  |  |
| HKL | 4FVT: -5.8 | 4FVT: F157, E323 | 4FVT: 1.00 | 85.9 | 109.8 |
|  | 4BVG: -6.9 | 4BVG: D156, N229 | 4BVG: 1.00 |  |  |

### Quantification of SIRT3 activation by HKL

Measurement of HKL’s effect on SIRT3 activity was carried out accounting for the fact that non-allosteric activators can have competing effects on the rates of enzymatic reaction steps (*29*), and hence can display qualitatively different activity modulation effects under different reaction conditions. The activation of SIRT3 by HKL was observed under non-steady state conditions, either at early time-points during the reaction (pre-steady state) with a low ratio of enzyme to limiting substrate concentration [E]_0_/[S]_0_ <<1 (**Figs. 8A** and **C**) or a high [E]_0_/[S]_0_ such that steady state was achieved later (**Figs. 8B** and **D**), where [S]_0_ is the initial concentration of limiting substrate NAD^+^ or acetylated peptide (**Fig. 8** for MnSOD-K122, **Fig. S2** for p53-AMC). The results indicated that the measurement time has a significant impact on the effect of HKL on SIRT3 activity. Under [E]_0_/[S]_0_<<1, SIRT3 activation was only detected at early times in the reaction. For example, the red, blue, and green curves in **Fig. 8A** were assayed at significantly longer times compared to the purple curve in **Fig. 8A**. The dose-response curves were also assayed at different concentrations of NAD^+^ and NAM, which affect the duration of the pre-steady state phase of reaction (note that 100 µM NAM is similar to the intracellular NAM concentration (*41*)). The observed activation of SIRT3 by HKL in **Fig. 8** is statistically significant, with p < 0.001. Time series analyses were carried out to further investigate the dynamics of activation of SIRT3 by non-steady state activators like HKL. The initial rate of deacetylation of the MnSOD substrate was observed to increase in the presence of HKL (**Fig. 8C**). When low enzyme concentrations were used, activation was only observed in the pre-steady state phase of the reaction. Under high ratio of enzyme to limiting substrate concentration ([E]_0_/[NAD^+^]_0_=0.3, non-steady state activation of SIRT3-catalyzed deacetylation by HKL was detected within a long-time window (up to 50 min) (**Fig. 8D**).

**Fig. 8.**
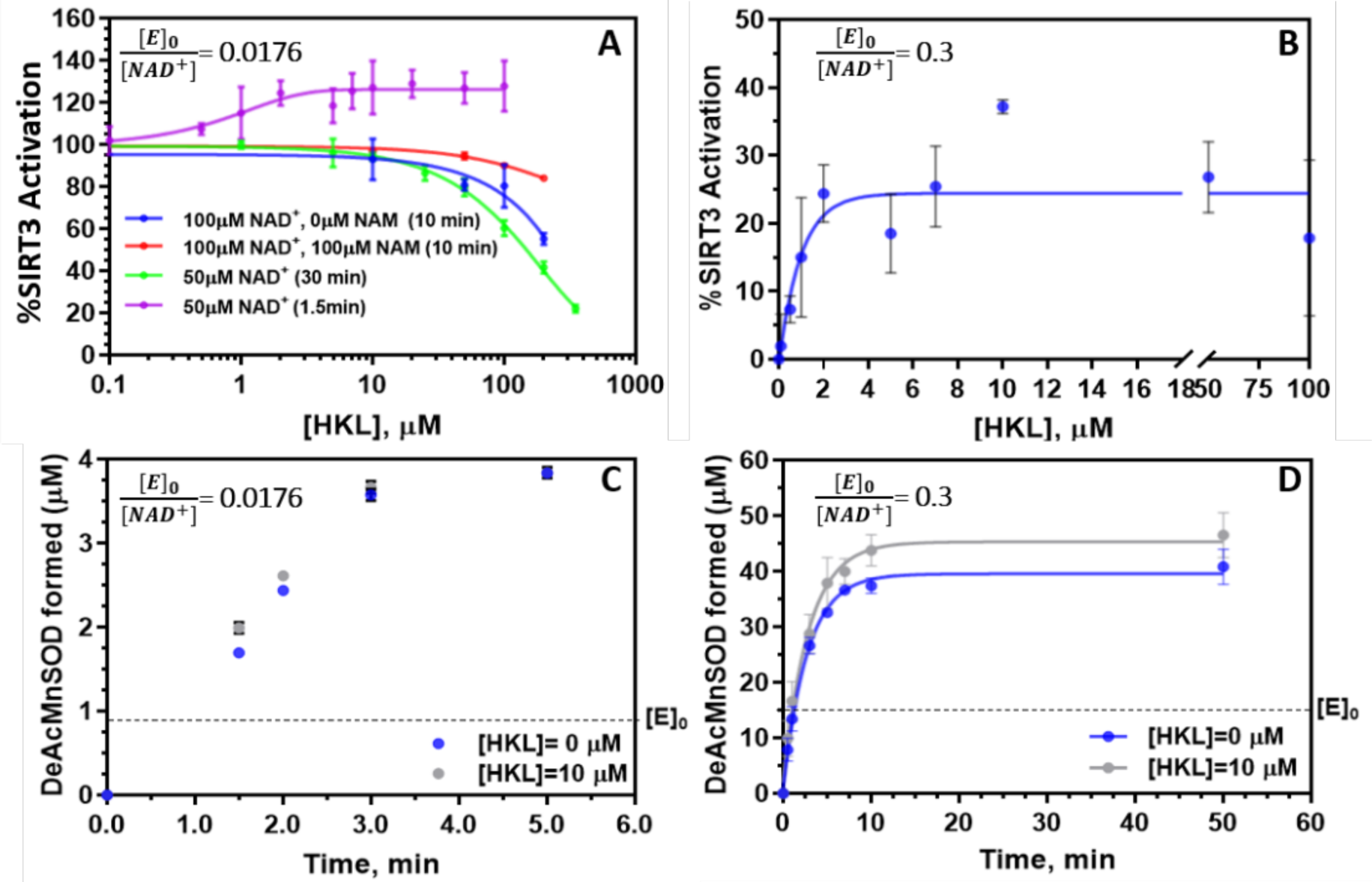
Effect of HKL on SIRT3 deacetylation activity; dose-response curves and time series of enzyme activation. **(A)** Dose-response curves were measured under conditions where [E]_0_/[NAD^+^]_0_= 0.0176, at various times. Non-saturating NAD^+^ and saturating MnSOD K122 peptide dose-response curves are presented in four formats of (blue) 10 mins, 100 µM NAD^+^ (N=2); (red) 10 mins, 100 µM NAD^+^/100 µM NAM (N=2), (green) 30 min, 50 µM NAD^+^ (N=3), and (purple) 1.5 min, 50 µM NAD^+^, N=2. (**B**) Dose-response curve for MnSOD substrate measured under higher enzyme concentration conditions, [E]_0_/[NAD^+^]_0_=0.3, in the presence of 50 µM NAD^+^ and 600 µM MnSOD substrate, t=5 min (N=3). Plots of product formation vs. time in the presence and absence of (**C**) 10 µM HKL for [NAD^+^]_0_ = 50µM, [MnSOD K122] = 600µM, [E]_0_/[NAD^+^]_0_ = 0.0176 (N = 2; error bars are omitted for clarity); (**D**) 10 µM HKL for [NAD^+^]_0_ = 50µM, [MnSOD K122] = 600µM, [E]_0_/[NAD^+^]_0_ = 0.3 (N = 2).

### Numerical simulation of activation by non-steady state SIRT3 activating compounds

As shown above (**Table 1**, **Fig. 8**), HKL and some newly discovered activators increase the non-steady rate of deacetylation but not the steady state rate, whereas other newly discovered activators increase the steady state rate of SIRT3-catalyzed deacetylation. In order to further analyze the result of these effects on the dynamics of SIRT3-catalyzed deacetylation over the course of the reaction, we carried out numerical simulations of the reaction dynamics under multiple initial conditions including low and high values of [E]_0_/[NAD^+^]_0_ at saturating acetylated peptide, over all phases of the reaction, for non-steady state activation by compounds like HKL. In these simulations, we applied the simplifying assumptions that a) all ligands have equal on rates in the absence of HKL and NAD^+^ has an off rate 10 times lower than that of NAM; b) HKL decreases the off rate of NAD^+^ by a factor of five, increases the rate of ADP ribosylation by a factor of five, and reduces the off rate of AADPR by a factor of 10. The chosen parameter values were based on the results of the MM-GB(PB)SA simulations above (**Fig. 5**) that show increase in O-AADPR binding affinity upon loop closure, and the experimental observation (**Fig. 8**) of enzyme activation under non-steady state conditions. The results are shown in **Fig. 9**, in which Top corresponds to a low value of [E]_0_/[NAD^+^]_0_ comparable to that used for the experiments in **Figs. 8A** and **C**, whereas Bottom corresponds to a high value of [E]_0_/[NAD^+^]_0_ comparable to that used for the experiments in **Figs. 8B** and **D**. The numerical simulation of activation under [E]_0_/[NAD^+^]_0_ = 0.01 (**Fig. 9 Top**) shows that the pre-steady state activation is followed by a long steady state phase showing inhibition. The net inhibition is consistent with the experimental inhibition under steady state conditions (**Fig. 8A**). The results from numerical simulation for [E]_0_/[NAD^+^]_0_ = 0.01 and [E]_0_/[NAD^+^]_0_ = 0.25 (**Fig. 9 Bottom**) of the effects on the three phases of the reaction (e.g., in the former case, a short pre-steady state activation by the modulator followed by a long steady state phase inhibition and finally a post-steady state phase) are also fully consistent with the experimental time series on the effects of HKL in **Figs. 8C** and **D**, respectively.

**Fig. 9.**
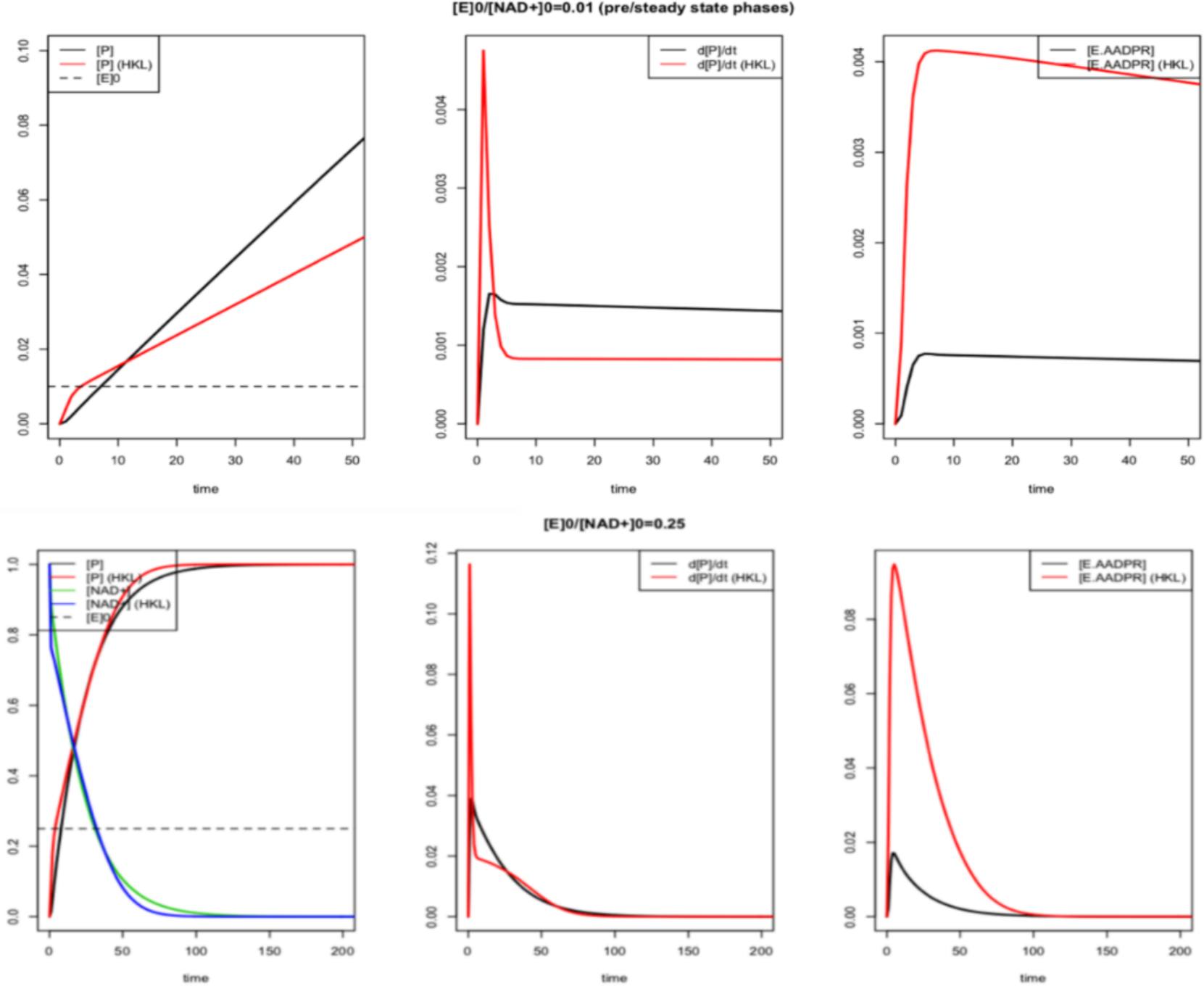
Numerical simulations of the dynamics of SIRT3-catalyzed deacetylation under different [E]_0_/[NAD^+^]_0_ conditions similar to those studied experimentally, in the presence and absence of a non-steady state activator like HKL. **Top**: [E]_0_/[NAD^+^]_0 =_ 0.01**; Bottom:** [E]_0_/[NAD^+^]_0_ = 0.25. P = deacetylated peptide product. Simulation parameters are defined in the text, and do not precisely correspond to the true parameter values for HKL. The horizontal dashed line depicts [E]_0._

### Validation of selected hit compounds for SIRT3 activation and identification of steady state activators

The *in vitro* validation of hit compounds from virtual screening was first performed with a commercial Fluor de Lys enzymatic assay to assess their modulation effects on SIRT3. While compounds from virtual screening were defined as hits as long as they produced statistically significant enzyme activity modulation (**Table S2**), additional characterization was required to assess whether the compounds upregulated SIRT3 activity and under what conditions (steady state and/or non-steady state, **Table 1**), given the identification of HKL as a non-steady state activator and steady state inhibitor. To avoid the controversy of sirtuin activation by small molecules induced by fluorophore-labeled peptides and to assess substrate dependence (*42*), a HPLC assay using a native SIRT3 substrate (acetylated MnSOD peptide) was applied to confirm the observations under both steady state and non-steady state conditions.

Under steady state screening conditions (low enzyme concentration, high reaction time), the modulation effect of the top 20 hit compounds in the presence of fluorophore-labeled peptide (FDL assay) and MnSOD peptide substrates (HPLC assay) is presented in **Fig. 10A**. A good correlation between the results obtained from both assays indicated that the tested compounds were not substrate specific SIRT3 modulators. Another 48 compounds with high docking scores were tested for their modulation effect on SIRT3 (HPLC results throughout the paper unless specified otherwise). Their modulation behavior is presented in **Fig. 10B**. The compounds identified as steady state inhibitors were also tested under non-steady state conditions (high enzyme concentration, low reaction time; **Fig. 11**). **Table S2** provides a summary of screening results for the top 68 compounds.

**Fig. 10.**
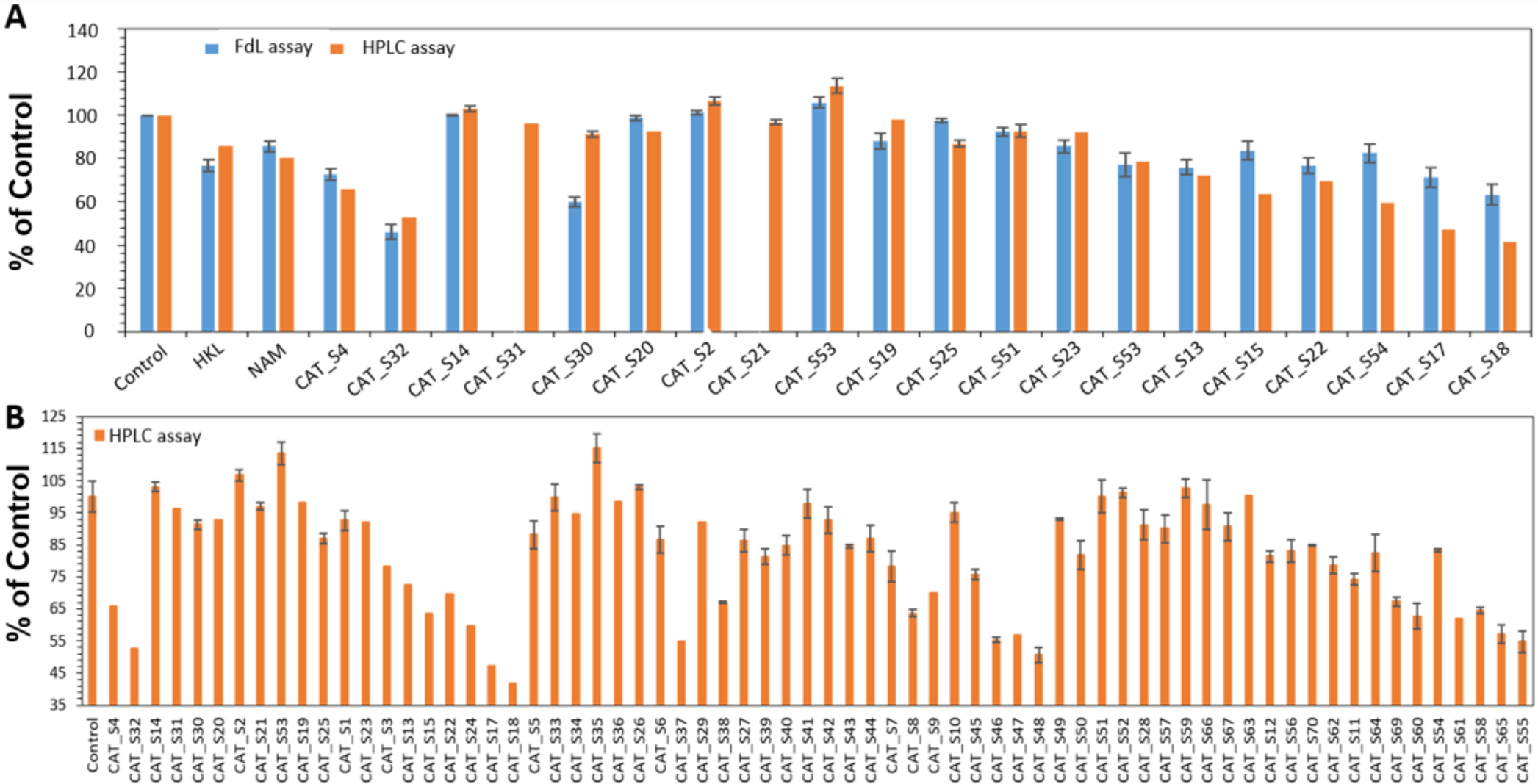
The modulation effect of hit compounds under steady state screening condition ([NAD^+^]_0_ =1 mM, [Peptide substrate] = 50 µM, [cpd] = 10 µM, [E]_0_/[AcPr]_0_ = 0.0049, [E]_0_/[NAD]_0_ = 0.000245, Time point=30 min, N=3) **(A)** Top 20 selected compounds in the presence of FDL and MnSOD peptides. (**B)** Top ranked 68 compounds using HPLC assay.

**Fig. 11.**
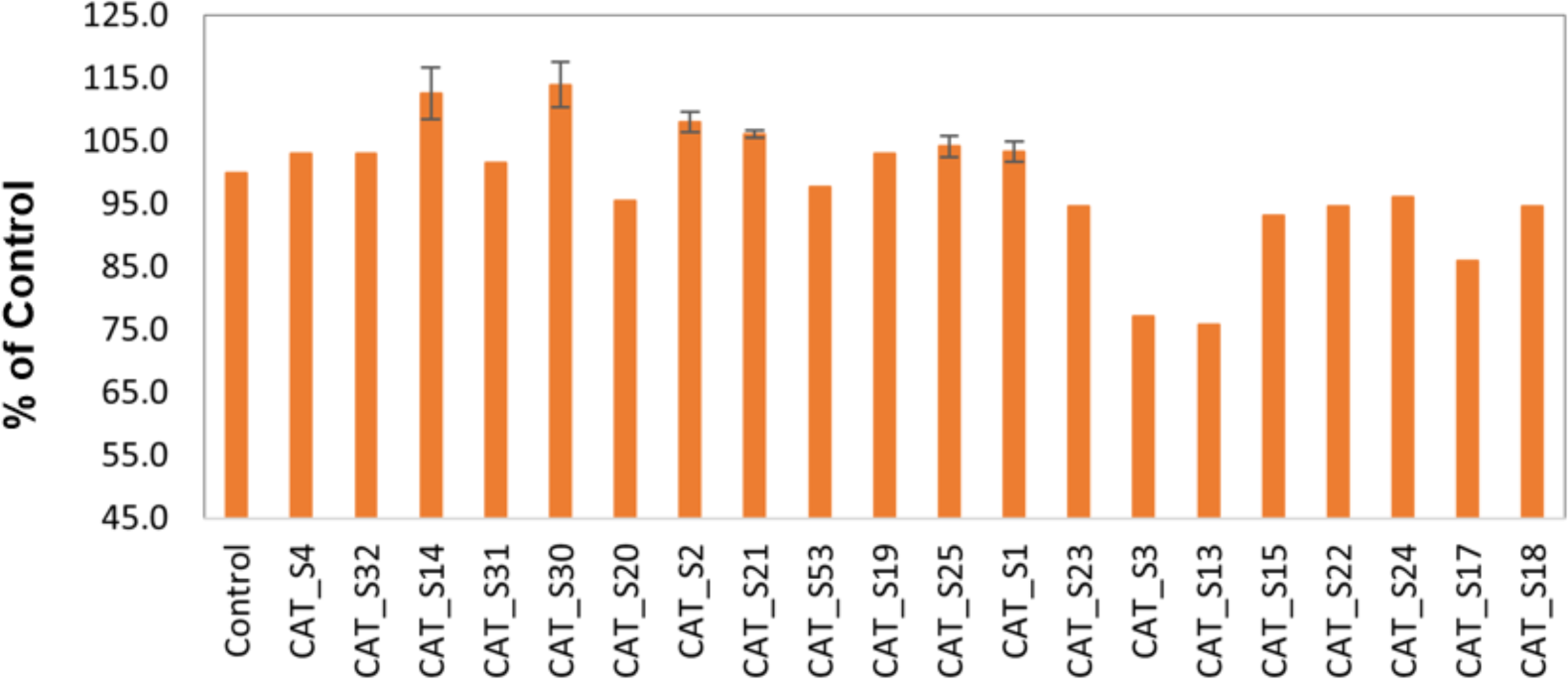
The modulation effect of the top 20 selected compounds under non-steady state screening conditions using the HPLC assay ([NAD^+^]_0_ =50 µM, [MnSOD K122] = 600 µM, [cpd] = 10 µM, [E]_0_/[NAD]_0_ = 0.1223, Time point=2 min, N=3).

Some of the tested compounds showed inhibition potency against SIRT3 in the range 0 – 58.4% at 10 µM modulator concentration (**Table S2**) under steady state conditions. Two hit compounds showed good inhibition potency (>50% inhibition) at a concentration of 10 µM. 9 out of 68 compounds were found to activate SIRT3 (3.1% - 14.0%) at 10 µM under non-steady state conditions (**Table 1**). The hit rate (the number of hits that modulate SIRT3 activity divided by the number of tested compounds) for VS is 48.6%, indicating the current VS workflow is successful.

Importantly, 6 compounds were found to activate SIRT3 under steady state conditions (**Table 1**), unlike HKL. Since they are the first reported steady state activators of SIRT3, 3 of these were extensively characterized. Dose-response curves (**Fig. 12A**) indicated that maximal activation of SIRT3 was detected in the presence of 1 μM compound CAT_S26. The potency of SIRT3 activation by 1 μM CAT_S26 and 20 μM CAT_S35 under physiologically relevant low NAD^+^ concentration (100 μM) was observed to be 199% and 213.8%, respectively (**Fig. 12B**), at 10 min reaction time.

**Fig. 12.**
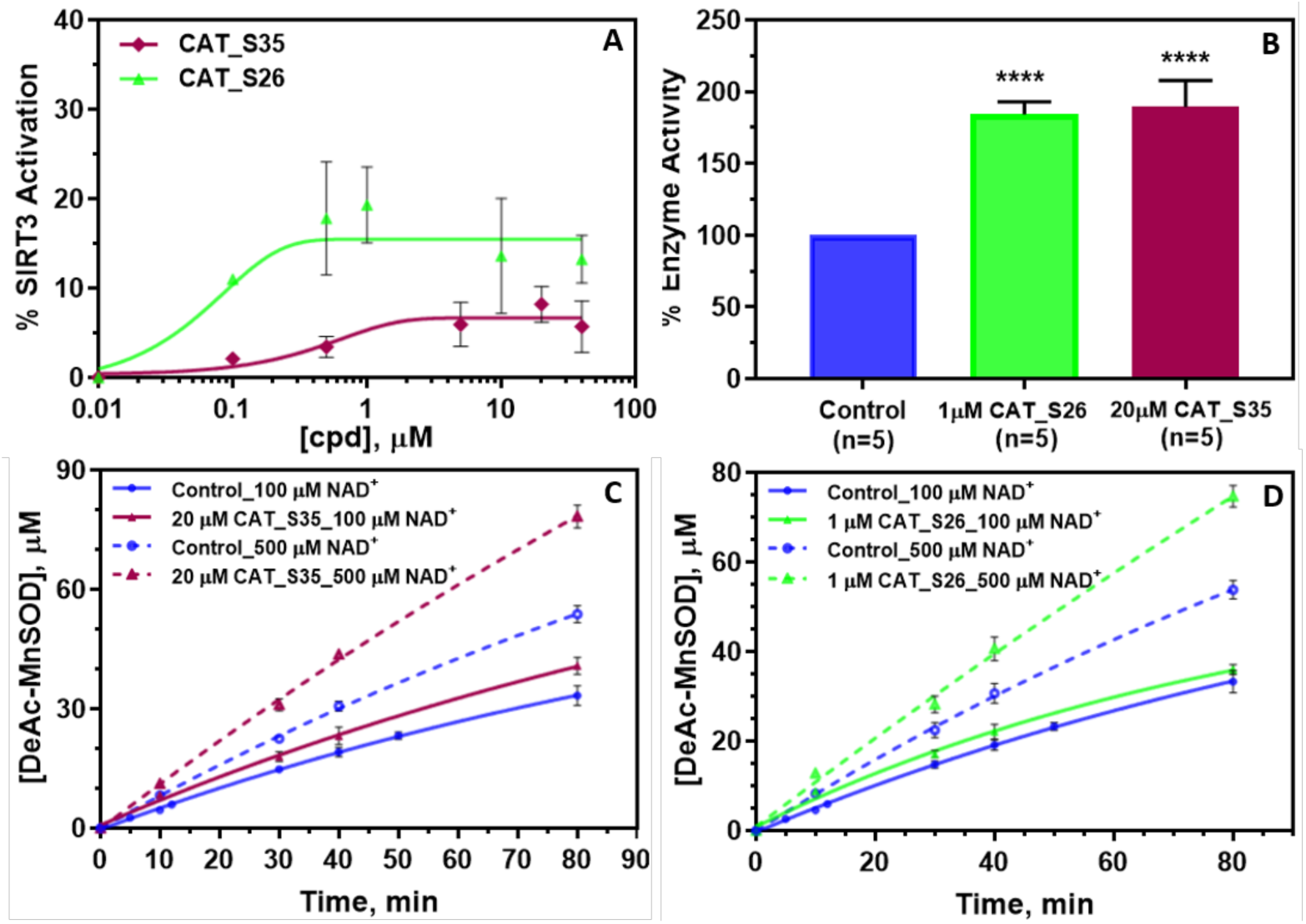
Effect of VS hit compounds on SIRT3 deacetylation activity under steady state conditions: dose-response curves and time series of enzyme activation. **(A)** Dose-response curves for 2 VS hit compounds under [E]_0_/[NAD^+^]_0_=0.00157, in the presence of 1 mM NAD^+^ and 50 μM MnSOD peptide, t=30 min, N=3. **(B)** Bar diagram of % control by 2 VS hit compounds (20 μM CAT_S35 or 1 μM CAT_S26) on SIRT3 deacetylation activity under steady state conditions where [NAD^+^]_0_ = 100 μM, [MnSOD K122] = 600 µM, [E]_0_/[NAD^+^]_0_ = 0.00157, t = 10 min, N=2. (* p < 0.001). Plots of product formation vs. time in the presence and absence of (**C**) 20 μM CAT_S35 for [NAD^+^]_0_ = 100 and 500 μM, [MnSOD K122] = 600μM, [E]_0_/[NAD^+^]_0_ = 0.00157 (N = 2); **(D)** 1 μM CAT_S26 for [NAD^+^]_0_ = 100 and 500 μM, [MnSOD K122] = 600μM, [E]_0_/[NAD^+^]_0_ = 0.00157 (N = 2).

Time series analyses were carried out to further investigate the dynamics of activation of SIRT3 by steady state activators identified through high-throughput screening. The initial rate of deacetylation of the MnSOD substrate was observed to increase in the presence of hit compounds (**Figs. 12C** and **D**) and activation persisted for over 80 min.

To summarize, our results establish that Honokiol is a non-steady state SIRT3 activator while having inhibitory effects on SIRT3 in the steady state. Steady state activators of SIRT3, such as CAT_S35, and CAT_S26, are preferable for therapeutic applications and overcome the inhibitory effects of non-steady state activators at longer times and lower enzyme concentrations (**Fig. 12**).

### Biophysical characterization of SIRT3 activation

#### Steady state characterization

Biophysical characterization of the mechanism of activator-induced modulation of SIRT3 was carried out, in order to elucidate the origin of the non-steady state activation and steady state inhibition by HKL, the steady state activation by newly reported compounds such as CAT_S35 and CAT_S26, as well as to inform the design of novel activators.

As described in (*29*) the steady state rate of deacylation can be expressed

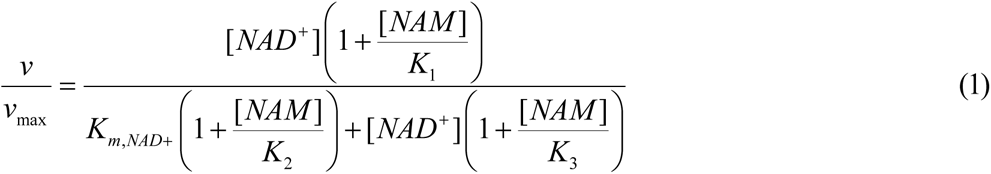

where the expressions for the steady state constants as functions of fundamental rate constants in the enzymatic reaction mechanism are presented in **Fig. S3**. Steady state rate measurements at multiple values of [NAD^+^] and [NAM], with saturating acetylated peptide, were carried out in order to estimate the parameters in this model. Saturating HKL was used to ensure the observed effects were due entirely to HKL-bound SIRT3. HKL:SIRT3 steady state rates were measured in the presence of both NAD^+^ and NAM, and the data fit into the full mixed inhibition rate expression (1), in order to elucidate its mechanism of action and determine the reasons behind its observed non-steady state activation and steady state inhibition (**Fig. 8**).

Double exponential time series fitting under conditions suitable for steady state analysis ([E]_0_/[NAD^+^]_0_ << 1, as in **Figs. 8A** and **C**) was used to identify pre-steady state and steady state phases, capture the initial rate enhancement due to HKL, and estimate the steady state rate (**Fig. S4**). The transition between the pre-steady state phase (which displays HKL-induced activation) and steady state phase (which displays HKL-induced inhibition) occurs after the concentration of deacetylated product exceeds the concentration of enzyme, i.e. after each enzyme molecule has catalyzed one reaction.

The steady state results for HKL with MnSOD substrate are presented in **Figs. 13A** and **B**, and **Table 2**. The purpose of this study is to compare the observed changes in the steady state parameters *v*_*max*_, *K*_*m*,*NAD*^+^_, *K*_1_, *K*_2_, *K*_0_, at 200 µM HKL concentration (approximately saturating) to the changes conducive to activation, which were delineated in reference (*29*) and that are depicted in **Figs. S3B** and **S5**. Note that at very high values of [NAM], the effect of differences in *K*_*d*,*NAM*_(**Fig. S6**) induced by the modulator becomes negligible, allowing isolation of differences in the NAD^+^ binding and ADP Ribosylation / base exchange kinetics.

**Fig. 13.**
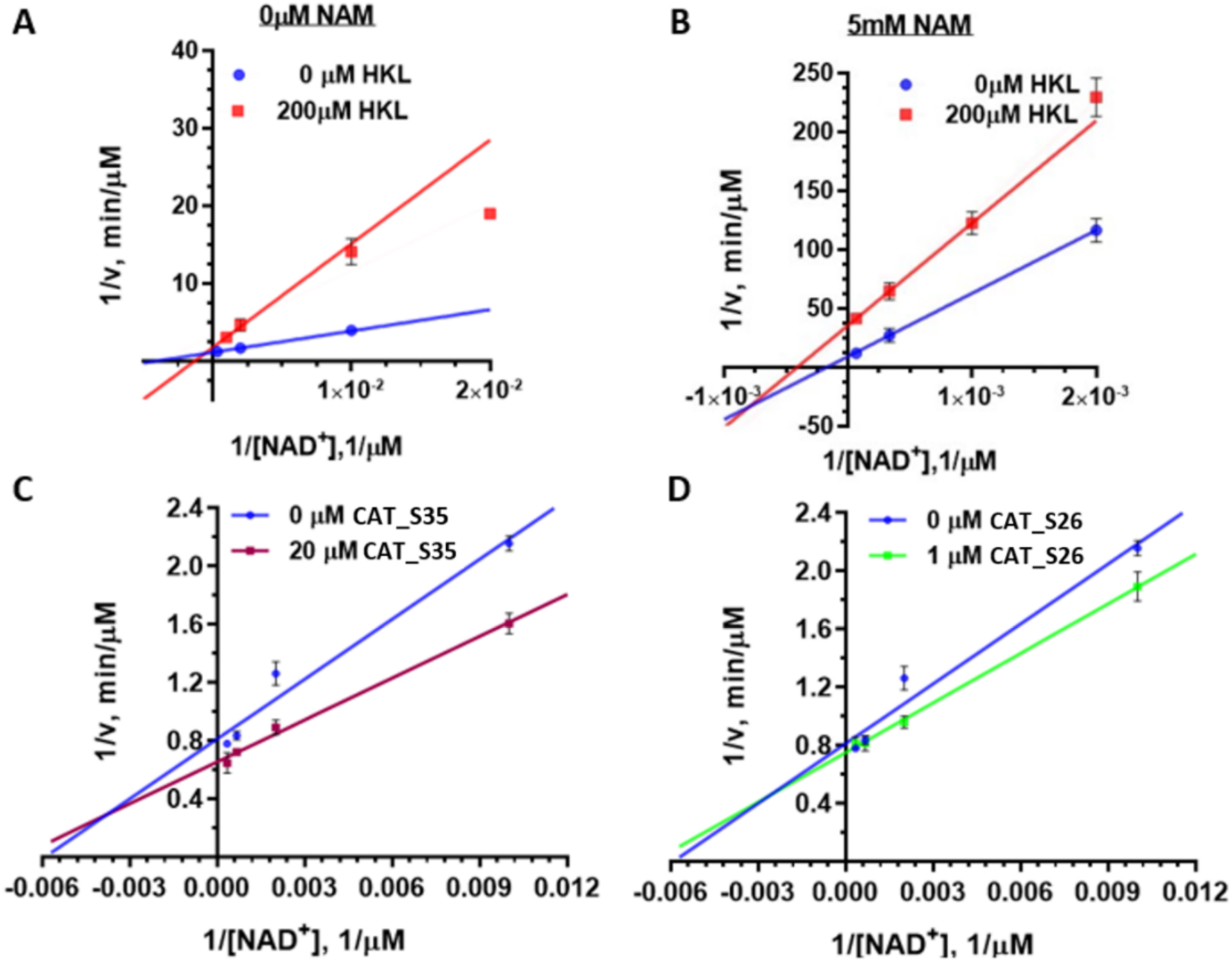
Steady state kinetic characterization of hSIRT3^102-399^ deacetylation of MnSOD substrate by non-steady state activator HKL and steady state activators. Double reciprocal plots for deacetylation initial rate measurements in the presence and absence of HKL at **(A)** [NAM] = 0 µM. **(B)** [NAM] = 5mM. **Fig. S7** shows the complete expression. Double reciprocal plots for SIRT3 deacetylation initial rate measurements in the presence and absence of **(C)** 20 µM CAT_S35 and **(D)** 1 µM CAT_S26. The initial rate was measured under [MnSOD K122] = 600 µM with different [NAD^+^]_0_ concentrations at series of time points (N=2).

**Table 2.** Model parameter estimates from global nonlinear fitting of Eq.(1) for SIRT3 in the presence and absence of 200 μM (saturating) HKL. [E_0_] = 1.85 µM.

| | 0 μM HKL | 200 μM HKL |
| --- | --- | --- |
| Vmax, μM/min | 0.8681 ± 0.012 | 0.566 ± 0.053 |
| K1, μM | 18064 ± 8279 | 3758 ± 1572 |
| Km, μM | 238.9 ± 12.92 | 756.6 ± 138.8 |
| K2, μM | 20.19 ± 1.612 | 33.23 ± 6.886 |
| K3, μM | 528 ± 105 | 108.1 ± 28.98 |
| Goodness of Fit |  |  |
| Degree of Freedom | 11 | 10 |
| R square | 0.9988 | 0.9947 |
| Absolute Sum of Squares | 0.001052 | 0.0006615 |
| Sy.x | 0.009782 | 0.008133 |

The steady state results for newly reported steady state activators (CAT_S35 and CAT_S26) are presented in **Figs. 13C** and **D**, and **Table 3**. For these activators, expression (1) was fit to the data with [NAM]=0. The results closely match the predicted properties in **Figs. S3B** and **S5C** for steady state mechanism-based sirtuin activators.

**Table 3.** Model parameter estimates from global nonlinear fitting of Michaelis-Menten for SIRT3 in the presence and absence of 1 μM CAT_S26 and 20 μM CAT_S35. [E_0_] = 1.568 µM.

| | Control | 20 μM CAT_S35 | 1 μM CAT_S26 |
| --- | --- | --- | --- |
| Vmax, μM/min | 1.381 ± 0.114 | 1.58 ± 0.0617 | 1.318 ± 0.029 |
| Km, μM | 270.8 ± 94.8 | 172.8 ± 32.7 | 142.6 ± 16.3 |
| Km/Vmax, min | 211.4 ± 56.6 | 109.4 ± 16.4 | 108.2 ± 9.9 |
| **Goodness of Fit** |  |  |  |
| Degrees of Freedom | 2 | 2 | 2 |
| R square | 0.9539 | 0.9827 | 0.9932 |
| Absolute Sum of Squares | 0.01999 | 0.008505 | 0.002213 |
| Sy.x | 0.09997 | 0.06521 | 0.03326 |

#### Measurement of binding affinities

Additional biophysical analysis in the form of binding affinity measurements was carried out to further elucidate the mechanism of SIRT3 activation by HKL. The binding affinities of ligands to complexes in the catalytic mechanism of SIRT3/MnSOD peptide substrate were measured using microscale thermophoresis (MST) (**Table 4**, **Figs. S8**, **S9** and **S10**). In order to carry out binding affinity measurements on the reactive complex of enzyme, acetylated peptide substrate and NAD^+^, we synthesized the catalytically inert molecule Carba-NAD and used this NAD^+^ analog for those MST studies.

**Table 4.** Summary of *K_d_* values reported in current study. The microscale thermophoresis (MST) was used to determine the binding affinities of HKL or hit compound CAT_S35 in the presence of cofactors, including reactants (Ac-MnSOD or Ac-p53-AMC), products (DeAc-MnSOD + OAADPr).

| Exp. Detail | Target Molecule | $K_d$, μM |
| --- | --- | --- |
| SIRT3 + 625 nM Ac-MnSOD | Carba-NAD | 7.3 ± 0.7 |
| SIRT3 + 625 nM Ac-MnSOD + 6.25 μM Honokiol | Carba-NAD | 6.6 ± 2.1 |
| SIRT3 + 625 nM Ac-MnSOD | Honokiol | 3.0 ± 0.02 |
| SIRT3 + 625 μM Ac-MnSOD + 1.25 mM NAM | Honokiol | 0.7 ± 0.17 |
| SIRT3 + 100 μM DeAc-MnSOD | Honokiol | 0.97 ± 0.21 |
| SIRT3 + 100 μM DeAc-MnSOD + 1.25 mM OAADPR | Honokiol | 0.29 ± 0.09 |
| SIRT3 + 625 μM NAD<sup>+</sup> + 100 μM DeAc MnSOD | Honokiol | 0.18 ± 0.02 |
| SIRT3 + 50 μM Ac-p53-AMC + 6.25 μM Honokiol | Carba-NAD | 7.1 ± 0.5 |
| SIRT3 + 50 μM Ac-p53-AMC | Carba-NAD | 5.5 ± 1.1 |
| SIRT3 | CAT_S35 | 0.0266 ± 0.0265 |

In addition to measuring the effect of HKL on NAD^+^ binding affinity, we measured its effect on acetylated peptide binding affinity, NAM binding affinity in the presence of acetylated peptide, deacetylated peptide binding affinity, O-Acetylated ADP Ribose binding affinity in the presence of deacetylated peptide (i.e., the product complex), and NAD^+^ binding affinity in the presence of deacetylated peptide (**Table 4**, **Figs. S8**, **S9** and **S10**). The latter measurements in the presence of deacetylated peptide were made to determine whether product inhibition plays any role in the observed kinetics. Importantly, the binding affinity of hit compound CAT_S35 with apo enzyme SIRT3 was also measured using MST. The nM range K_d_ value of CAT_S35 indicates the hit compound is a very potent binder to SIRT3 (**Table 4** and **Fig. S8F**).

## DISCUSSION

The catalytic cycle of sirtuin enzymes (**Fig. S1**) proceeds in two consecutive stages (*2*). In the first stage (ADP ribosylation), the NAM moiety of NAD^+^ is cleaved through nucleophilic attack of the protein substrate’s acyl-Lys side chain to create a positively charged O-alkylimidate intermediate (*2, 7*). NAM-induced reversal of the intermediate (the “base exchange” reaction) results in regeneration of NAD^+^ and acyl-Lys protein. The thermodynamics of this reversible reaction affects both the potency with which NAM inhibits sirtuins as well as *K_m,NAD^+^_* (the Michaelis constant for NAD^+^). Stage 2 of sirtuin catalysis includes the rate-determining step and involves four successive steps that conclude in deacylation of the protein substrate’s Lys side chain and generation of the coproduct, O-Acetyl ADP Ribose (*2, 4, 43*).

Using **Eq. (1)**, it is possible to determine the initial rate of deacylation for specified (e.g., intracellular) concentrations of NAD^+^ and NAM, if the rate constants are known. At constant [NAM], **Eq. (1)** is usually represented graphically as double reciprocal plots, where the slope of the plot (1/v *vs* 1/ [NAD^+^]) at [NAM] = 0 is *K*_*m*,*NAD*^+^_/*v*_*max*_, for which the expression is (*44*):

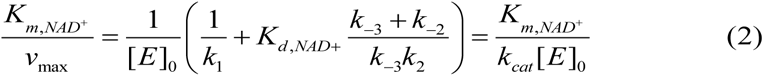

where k_1_ denotes the rate of NAD^+^ binding, k_2_ and k_-2_ respectively denote the ADP ribosylation forward and reverse (base exchange) rates, and k-_3_ denotes the rate of NAM dissociation (usually fast such that k_-3_ and k_-2_ can be eliminated from the expression). The steady state parameter α, which measures the extent of competitive inhibition by NAM, the endogenous inhibitor, against the NAD^+^ cofactor, can be written in terms of the ratio of *K*_*d*,*NAD*^+^_ and *K*_*m*,*NAD*^+^_ (*44*):

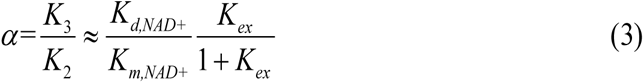

where *K*_*ex*_ = *k*_−2_/*k*_2_ and the approximations are defined in **Fig. S3**. (The corresponding expressions for the peptide substrate are analogous, with peptide replacing NAD^+^ in the equations above.)

For enzyme activation to be possible, the modulator must co-bind with NAD^+^ and acylated peptide. Moreover, non-allosteric activators must bind in the vicinity of the active site and modulate local conformational degrees of freedom to enhance enzyme activity. In our previous study (*29*), we presented a theory of mechanism-based enzyme activation, for the purpose of developing a foundation for the characterization and design of non-allosteric enzyme activating compounds. **Fig. S11** illustrates the theoretical model for mechanism-based sirtuin enzyme activation schematically. This theory provides a unifying framework under which such compounds can be characterized, and hits can be evolved into leads. We apply this theory to the experimental data we collected for HKL and the activators discovered herein, identify the reasons that HKL does not activate SIRT3 under steady state conditions, and explain how the steady state activators discovered in the present work overcome these limitations.

### Characterization of SIRT3 modulation by the non-steady state activator Honokiol

The data reported above indicate that HKL co-binds with the peptide substrate and cofactor of SIRT3. Under the conditions tested (saturating HKL), the net effect of SIRT3 on MnSOD peptide substrate was pre-steady state activation followed by steady state inhibition (**Figs. 8** and **13A**). The mechanistic model for non-allosteric activation (**Fig. S1**) is required to elucidate the effects of HKL on the kinetic parameters of the enzymatic reaction, explain HKL’s activation of SIRT3 activity under physiological non-steady state conditions, and understand how its properties as an activator differ from those of the novel steady state activators discovered herein.

The structural and computational results in **Figs. 1-5** demonstrate that HKL cobinds with SIRT3 substrates near the active site and induces a change in the conformation of the cofactor binding loop due to the latter’s flexibility. It is therefore not an allosteric modulator. The induced changes in the cofactor binding loop conformation affect the binding of NAD^+^ as well as that of the AADPR coproduct. This conformational change is predicted to have a significantly favorable effect on AADPR coproduct binding affinity (**Fig. 5**), whereas its effect on NAD^+^ binding affinity is predicted to be less significant due to the presence of NAM in the C pocket where Phe157 resides upon loop closure (**Fig. 1**). Moreover, due to the interaction of the modulated loop conformation with the nicotinamide moiety of NAM, the resulting change in NAD^+^ binding geometry (**Fig. 4**) is expected to alter the ADP ribosylation / base exchange rate constants of the reaction.

We now consider the experimentally measured effects of HKL on SIRT3 ligand binding affinities and steady state kinetics. According to **Table 4**, **Figs. S8**, **S9** and **S10**, *K*_*d*,*NAD*^+^_ is roughly unchanged or slightly reduced for the MnSOD substrate. Complementary information is obtained from the effect of HKL on steady state kinetic parameters. α decreases several fold in the presence of HKL (**Table 2**). This decrease in α is due to the fact that *K*_*m*,*NAD*^+^_ increases in the presence of HKL while *K*_*d*,*NAD*^+^_ does not increase / decreases slightly (see **Eq. (3)**). HKL’s effect on *K*_*d*,*NAD*^+^_ is consistent with this reduction in *⍺* ∗ *K*_*m*,*NAD*^+^_. According to **Eq. (3)**, *⍺* ∗ *K*_*m*,*NAD*^+^_ is highly correlated with *K*_*d*,*NAD*^+^_ and the methods are complementary, since MST employs a chemically modified NAD^+^ analog. A slight decrease in *K*_*d*,*NAD*^+^_ but increase in *K*_*m*,*NAD*^+^_ can explain the observed non-steady state activation of MnSOD substrate deacetylation. As described in the SI, a complete analysis of the HKL characterization data in the context of the non-allosteric activation model (**Fig. S11**) indicates that non-steady state activation of SIRT3 by HKL may also be enhanced by its effect on the rate of ADP ribosylation through its induction of loop closure.

In the presence of a non-steady state activator like HKL, the SIRT3-catalyzed deacetylation reaction goes through 3 phases: i) the pre-steady state phase during which each enzyme molecule catalyzes a limited number of reaction cycles, and which is faster in the presence of HKL (**Fig. 8**); ii) the steady state phase during which each enzyme molecule may catalyze many reaction cycles, and which is slower in the presence of HKL; iii) the post-steady state phase which is faster in the presence of HKL. Depending on the initial ratio of enzyme to limiting substrate in the system, the net effect of HKL over the course of the reaction can be either inhibitory or activating, with activation occurring if this ratio is above a threshold value (**Fig. 9**).

The role of product inhibition in non-steady state activation by HKL is apparent from **Fig. 9**. It can be seen that activation occurs at times prior to which the amount of product formed is roughly equal to the enzyme concentration [E]_0_. The dose-response curves in **Fig. 8A** include measurements from reaction phase i. The dose-response curves for 50 µM NAD^+^ (1.5 min) and 100 µM NAD^+^, 100 µM NAM (10 min) (**Fig. 8A**) include contributions from phases i and ii of the reaction, whereas the dose-response curves for 50 µM NAD^+^ (30 min) and 100 µM NAD^+^ (10 min) have dominant contributions from phase ii (steady state phase). **Fig. 8B** demonstrates how a higher concentration of enzyme results in net activation. The results from numerical simulation of SIRT3 reaction dynamics for various values of [E]_0_/[NAD^+^]_0_ (**Fig. 9**) are fully consistent with the analysis above of the effects of HKL on the three phases of the reaction, and the pre-steady state activation / steady state inhibition observed experimentally. In **Fig. 9**, Top ([E]_0_/[NAD^+^]_0_ << 1), note the pre-steady state activation by the modulator followed by a long steady state phase and finally a post-steady state phase – closely matching the experimental results in **Fig. 8A**. In **Fig. 9**, Bottom (higher [E]_0_/[NAD^+^]_0_), note the net activation, as well as a reduction in the net extent of activation at intermediate times during the steady state – again, closely matching the experimental results under these conditions in **Fig. 8B**.

### Steady state SIRT3 activating compounds

Importantly, our drug discovery workflow (**Scheme 1** in Supplementary Methods) identified the first reported steady state activators of SIRT3 (**Table 1**), which improve qualitatively upon the properties of Honokiol and are ideal candidates for SIRT activator drug development. The steady state activators CAT_S35 and CAT_S26 were characterized and found to provide a 184.5% (at 20 μM) and 194.4% (at 1 μM) increase in SIRT3 activity, respectively, under steady state conditions at the physiologically relevant [NAD^+^]_0_=100 μM (**Fig. 12B**), with the catalytic efficiency (k_cat_/K_m_) of SIRT3 being nearly doubled by both of these compounds (**Table 3**). The MST binding affinity result for compound CAT_S35 falls in the nM range which indicates that compound CAT_S35 is a strong binder to SIRT3 (**Fig. S8F**). The steady state parameter estimation (**Table 3**) shows that the catalytic efficiency is improved primarily through reduction in K_m_, as originally proposed in our prior theoretical study (*29*), which predicted that this is possible to achieve through modulation of the conformation of the flexible loop induced by active site binding of properly designed small molecules. The observed results in **Table 3** and **Fig. 13** (Michaelis-Menten plots) closely match the predicted properties in **Figs. S3B** and **S5C** for steady state mechanism-based sirtuin activators. Dose-response curves for compound CAT_S26 (**Fig. 12A**) also show that these compounds achieve over 50% of their maximal activation effect (AC50) at 100 nM and < 1 uM concentrations respectively, which is promising for further development of these hits into drug-like leads.

By approximately doubling the deacetylation rate of SIRT3 at 100 μM NAD^+^, which is similar to the NAD^+^ concentration in aged mitochondria since mitochondrial NAD^+^ is ∼ 200 µM in young age and can drop by 50% or more in old age (*45, 46*), the level of SIRT3 activation by these compounds is sufficient to largely recover the mitochondrial deacetylation activity of SIRT3 characteristic of young age. Moreover, steady state activators like these compounds will be less affected by variations in physiological conditions pertinent to SIRT3 activity, such as enzyme expression level, than Honokiol. As shown above (**Fig. 8**), non-steady state activators such as Honokiol require elevated levels of SIRT3 to exert a potent and sustained enhancing effect on enzyme activity, with inhibition ensuing at low enzyme levels. Indeed, Pillai et al. (*24*) established the enhancement of SIRT3 activity on the MnSOD substrate under larger ratios of enzyme to substrate concentrations and demonstrated that HKL also induces an increase in SIRT3 expression level in mice.

SIRT3 activation by these steady state activators was demonstrated under both saturating NAD^+^ (**Fig. 12A**) and saturating peptide substrate (**Figs. 12B-D**) conditions. The activators thus improve the catalytic efficiency irrespective of whether it is the on rate k_1_ and the dissociation constant K_d_ of the cofactor or peptide substrate that appear in **Eq. (2)**, which as noted also applies to saturating NAD^+^ conditions if NAD^+^ in the expression is replaced by acetylated peptide. Therefore, this result demonstrates that activation by these compounds is likely not only a consequence of improvement in cofactor and/or substrate binding affinity, but according to **Eq. (2)** also likely involves an enhancement of the rate of ADP ribosylation (k_2_), since the off rate of NAM (k_-3_), the base exchange rate (k_-2_) and k_cat_ do not appear in the expression for catalytic efficiency under the commonly applied assumption of fast NAM dissociation (27). By contrast, since the catalytic efficiency of SIRT3 decreases in the presence of HKL (**Table 2**), the latter’s effect on the ADP ribosylation rate is likely not as favorable. In addition, the observed activation by these compounds under steady state conditions demonstrates that each molecule of SIRT3 bound to such a compound can catalyze multiple reaction cycles at a faster rate than SIRT3 alone, unlike HKL-SIRT3, and hence such compounds may not induce as tight binding of the AAPDR coproduct as HKL does (**Fig. 5**). Given the crystallographic structures of HKL-SIRT3 complexes (**Fig. 3**), and the prediction of similar binding modes for these compounds interacting with the flexible loop, the balance of effects required for non-allosteric steady state activation apparently occurs due to stabilization of optimal cofactor binding loop conformations by the compounds.

### Comparison of receptor-ligand interactions for top activator hits across receptors

The residues participating in interactions with modulators, especially in the flexible loop area, were analyzed in further detail for the top validated activator compounds (**Table 5**) across these receptors. In addition, the HKL:SIRT3:AcPr:Carba-NAD receptor (HKL-SIRT3_2.4Å) was used for redocking the top validated compounds, in order to compare the interactions and the predictive accuracy of docking to ternary receptors co-crystallized in the presence of the known activator HKL vs the ternary (open loop) and intermediate (closed loop) complex receptors 4FVT and 4BVG, respectively. For certain non-steady state activators (CAT_S14, CAT_S25, and CAT_S30), Pro 155 and Tyr 204 were found in common to interact with the compounds through H-bonds under the closed loop conformation. On the other hand, for certain steady state activators (CAT_S35, CAT_S53, and CAT_S59), H-bond interactions were detected with Phe 157, Arg 158, Ser 159, Glu 177, Glu 198, Gln 228, and Glu 323 for the closed loop conformation, and only Glu 198 is found common in the selected list of compounds for the open loop conformation of SIRT3. The same residues also interact with top docked poses of HKL in both structures. Importantly, Pro 155, Phe 157, Glu 177, are residues in the loop region, with significantly higher residue mobility than the rest of the protein. Proper definition of the docking site, docking template, and ligand query play an important role in achieving the high hit rate (20% for activators and 48.6% for inhibitors and activators) for the current VS protocol.

**Table 5.**
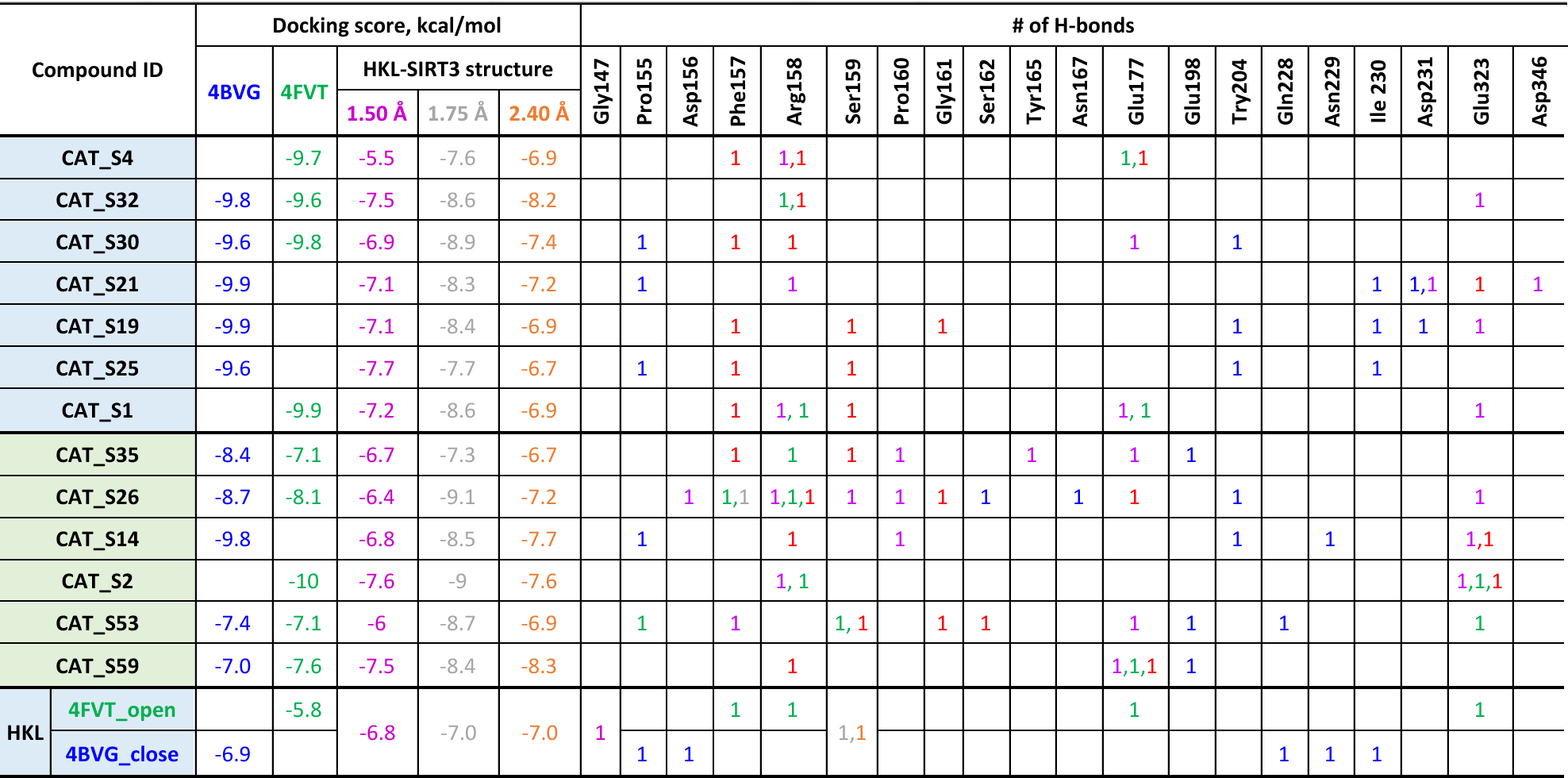
Summary of the ligand-protein interactions observed in docking for top hit compounds. The compounds showing activation effect under non-steady state condition are highlighted in blue; under steady state condition were highlighted in yellow; under both conditions were highlighted in green. 5 receptors were applied in the docking study, where 4BVG is in blue, 4FVT in green, HKL-SIRT3-1.50Å structure (HKL:SIRT3:AcPr) in purple, HKL-SIRT3-1.75Å structure (SIRT3: AcPr:Carba-NAD, HKL unresolved) in gray, HKL-SIRT3-2.40Å structure (HKL:SIRT3:AcPr:Carba-NAD) in orange. The docking score and # of H-bonds for hit compounds vs. different receptors were marked using same color code respectively.

In summary, we have applied a biophysical framework for activation of sirtuin enzymes to develop and apply an integrated computational and experimental drug design methodology for non-allosteric activators of the SIRT3 enzyme. Using this methodology, we identified novel SIRT3 activators, including the first reported compounds that activate SIRT3 under steady state conditions, and we presented and analyzed results from computational, kinetic and thermodynamic studies on SIRT3 activators.

We have shown that the compound Honokiol, previously reported to be a SIRT3 activator (*24*), does activate the enzyme under physiologically relevant non-steady state conditions but does not activate it under steady state conditions. By applying a) crystallography and computational modeling; b) experimental non-steady state, steady state and thermodynamic assays, and using the theory of mechanism-based enzyme activation (*29–31*) to interpret the data, we explained the mechanism behind Honokiol’s enhancement of SIRT3’s non-steady state deacetylation rate, demonstrated it is a non-allosteric modulator, and established a foundation for further hit-to-lead evolution of Honokiol and related compounds in order to convert them into more potent, steady state activators. Building upon these results, a 1.2 million compound library was screened virtually and Honokiol-like hit compounds were successfully identified.

Importantly, our drug discovery workflow also identified the first reported steady state activators of SIRT3. These steady state activators, which overcome limitations in Honokiol’s mechanism of action, were shown to increase up to two-fold the catalytic efficiency of SIRT3 and its deacetylation rate under NAD^+^ depletion conditions characteristic of old age. As such, compounds of this type have the potential to rejuvenate the activity of the major mitochondrial sirtuin to levels observed in youth. In particular, these effects were observed with the important SIRT3 substrate MnSOD, which plays a critical role in combating oxidative stress in mitochondria and whose upregulation has been demonstrated to rejuvenate aged cells including human oocytes (*47*). Several of these compounds display nanomolar AC50s and did not display toxic effects when applied to mouse oocytes at twice their concentrations of maximal activation (data not shown), potentially presenting a broad therapeutic window; as such, these compounds are suitable for *in vitro* preclinical studies using mouse models of age-related conditions, for example in the context of *in vitro* fertilization.

The present study serves as a successful proof of concept example for the current novel strategy for the field of non-allosteric activator discovery. As such, the present study constitutes an important foundation for the development of a new class of drugs for the treatment of age-related diseases that operate through a wholly new mode of action not shared by any existing drug.

Further characterization of these compounds, based for example on the methodologies presented in references (*29–31*), will facilitate the rational design of even more potent steady state activators. In addition to enabling further improvement of the identified novel steady state SIRT3 activators, these methodologies enable the evolution of hits that activate enzymes under only non-steady state conditions (such as several of the novel hits identified herein) into steady state activators. This is achieved through decomposition of the observed kinetic and thermodynamic effects of a modulator into biophysical parameters of the enzymatic reaction, and identification of those parameters that require optimization through hit-to-lead evolution (*44*). Such rational activator design workflows, whether applied to non-steady state or steady state activator hit compounds, would bear more similarity to enzyme directed evolution (*31*) than to traditional drug discovery workflows. Future work can apply enzyme engineering-inspired methods to hit-to-lead evolution of non-allosteric activators, in conjunction with structure-based computational methods analogous to those developed for computational enzyme design (*48*).

## MATERIALS AND METHODS

### Chemicals and Reagents

MnSOD (KGELLEAI-(KAc)-RDFGSFDKF) was synthesized by GenScript (Piscataway, NJ). Carba-NAD was synthesized by Dalton Pharma Services (Toronto, ON, Canada). FDL (QPKK^AC^-7-amino-4-methylcoumarin) peptide, also called p53-AMC peptide, was purchased from Enzo Life Sciences (Farmingdale, NY). Honokiol was purchased from Sigma-Aldrich (St. Louis, MO). Honokiol solubility was measured by Ascendex Scientific (Bristol, PA).

### Expression and purification of hSirt3^102-399^

hSirt3^102-399^ was expressed in *E.coli* Arctic Express (DE3) cells (Agilent Technologies, Wilmington, DE), as previously described (*30*). A single bacterial colony was inoculated in LB media containing ampicillin and gentamycin at 37°C. For protein purification, we inoculated overnight grown culture into 200 ml LB medium and grown at 30°C, 250 rpm for 4 hours. We induced Sirt3 expression by adding 1 mM IPTG at 15°C. After 24 hours, the cells were harvested, and the pellet was re-suspended in A1 buffer (50 mM NaH_2_PO_4_, 300 mM NaCl, 10 mM imidazole, pH 8.0) and sonicated. A clear supernatant was obtained by centrifugation at 14000 × g for 25 min at 4°C then loaded onto a 5 ml HisTrap HP column, pre-equilibrated with A1 buffer, attached to an AKTA pure FPLC system (GE Healthcare, Pittsburgh, PA). The column was then washed with 50 ml of each buffer A1, A2 (50 mM NaH_2_PO_4_, 300 mM NaCl, 75 mM imidazole, pH 8.0), A3 (20 mM Tris-HCl, 2M urea, pH 6.8), and again with buffer A2. Protein was eluted with buffer B1 (50 mM NaH_2_PO_4_, 300 mM NaCl, 300 mM imidazole, pH 8.0). The protein fractions were pooled, dialyzed against dialysis buffer (25 mM Tris, 100 mM NaCl, 5 mM DTT, 10% glycerol, pH 7.5) at 4°C overnight. The purity of final protein was >85% as assessed by SDS-PAGE.

### SIRT3 Crystallization and Structure Solution

SIRT3 complexes were crystallized by the vapor diffusion method. SIRT3 (118–399) (10.3 mg/ml) was crystallized in complex with LEAI(K-AC)RFFGS peptide (C10 peptide, 3 mM) and honokiol (1 mM) in 25% PEG 3350, 0.2 M Li_2_SO4 (or 0.2 M NaCl), and 0.1M HEPES, pH 7.5 as reservoir. Following formation of the ternary complex, crystals were soaked with Carba-NAD (10 mM). X-ray diffraction data were collected at beamline BL-13-XAVOC at the ALBA synchrotron in Cerdanyola des Valles, Barcelona, Spain. Data from samples that appeared to be of the best resolution and data quality were indexed and integrated using autoprocessing software at the beamline (XDS); merged, sealed and averaged using AIMLESS (CCP4 suite).

Structures were solved by the molecular replacement approach using the program Phaser from CCP4 suite and the SIRT3 molecule from SIRT3/ Fluor-de-Lys/Piceatannol complex refined against 2.3 Å data (PDB code 4HD8) as a search model. Structures were refined iteratively using the program Refmac(*49*) using the maximum likelihood target functions, isotropic individual B-factor refinement, and models were manually rebuilt in Coot(*50*), with 5% of the reflections excluded from refinement for *R*_free_ calculation(*51*). The ligand restraints for both the FDL and Carba-NAD molecules were generated with the program Acedrg from CCP4 suite. The protein moiety, FDL and Carba-NAD moiety are clearly visible and ordered in the final electron density of the refined model. The PDB IDs for the co-crystal structures mentioned throughout the paper will be released later.

### Steady state kinetic characterization of Honokiol on hSirt3^102-399^ deacetylation activity for MnSOD and FDL peptides

The steady state kinetic parameters were determined using varying subsaturating concentrations of NAD^+^ with 600 µM MnSOD peptide substrate (saturating peptide assays), in the presence and absence of HKL (0-200 µM), in the following buffer: 50mM Tris-HCl, 137mM NaCl, 2.7mM KCl, and 1mM MgCl_2_, pH 8.0 and 5% DMSO. Reactions were started by adding 5U hSirt_3_^102-399^ and then incubated at 37°C. Reactions were terminated using TFA at desired time points. The peptide product and substrate were resolved using HPLC.

For reactions with FDL (p53-AMC peptide), the same buffer was used with 100 µM FDL in the presence of different [NAD^+^]_0_. These reactions were terminated at specified time by using 1X developer 2mM NAM solution and fluorescence measured on a TECAN microplate reader.

### Effect of Honokiol and hit compounds on hSirt3^102-399^ deacetylation

To access the modulation effect of HKL or hit compounds on hSirt3, enzymatic reactions were performed under either 1 mM NAD^+^ and 50 µM MnSOD peptide and 5U hSirt_3_^102-399^ with 30 minutes incubation (steady state condition) or 50 µM NAD^+^ and 600 µM MnSOD substrate and 50U hSirt_3_^102-399^ with 2 minutes incubation (non-steady state condition) in presence of 10 µM hit compounds including HKL. The reactions were incubated at 37°Cand terminated by addition of TFA. The peptide product and substrate were resolved using HPLC. For reactions with FDL (p53-AMC peptide), similar conditions were applied as mentioned above.

### Separation of MnSOD and FDL peptides

An Agilent 1260 infinity HPLC system and a ZORBAX C18 (4.6×250 mm) column were used to separate deacetylated peptide and acetylated substrate using gradient comprising 10% acetonitrile in water with 0.005% TFA (solvent A) and acetonitrile containing 0.02% TFA (solvent B) 1 ml/min flow rate. Upon sample injection, the HPLC was run isocratically in solvent A for 1 min followed by a linear gradient of 0-51 %B over 20 min. After separation, the columns were washed with 100% solvent B and then equilibrated with 100% solvent A. The detector was set at 214 nm. The percent product produced was calculated by dividing the product peak area over the total area. GraphPad Prism software was used (GraphPad Software, Inc, CA) to fit the data.

### hSirt3^102-399^- ligand binding assay by Microscale Thermophoresis

Microscale thermophoresis experiments were performed by 2bind GmbH (Regensburg, Germany). In brief, the protein was conjugated with DY-647P1-NHS-Ester in 1x PBS pH 7.4, 0.05 % Pluronic F-127, and the labeled protein was buffer exchanged into 47 mM Tris-HCl pH 8.0, 129 mM NaCl, 2.5 mM KCl, 0.94 mM MgCl_2_, 5% DMSO, 0.05% Tween-20. We used 2 nM labeled hSirt3 and titrated with varying concentrations of the modulators in the absence and presence of various concentrations of substrates, products and intermediates. The thermophoresis was measured (excitation/emission 653/672 nm) using a Monolith NT. 115 Pico (NanoTemper Technologies) at 25°C. Dissociation constants (*K_d_*) were determined using GraFit7 (Erithacus Software) by nonlinear fitting.

### Preparation of SIRT3.Ac-Pr.NAD^+^ substrate complex for protein-ligand docking

#### Modeling the SIRT3 complex structure

The SIRT3 receptor structure for docking was prepared by converting Carba-NAD in pdb structure 4FVT into NAD^+^. An additional receptor structure for docking was prepared by replacing the cofactor binding loop in 4FVT with that from 4BVG. The complex structures were subjected to the Protein Preparation Wizard module of Schrodinger Suite. The module assigns bond orders and adds hydrogens based on Epik calculations for proper pKa values. Het molecules except for native ligands were eliminated and optimized structure by the Prime module of Schrodinger suite. The side chain structure of His248 was modified to HIE, because of the proximity of the histidine N to the Ribose oxygen of NAD^+^.

#### MD simulation protocol

The SIRT3-ALY-NAD^+^ complex was subjected to molecular dynamics simulation as follows. The complex is kept into a suitably sized box, of which the minimal distance from the protein to the box wall was set to 10Å. The box is solvated with the SPC (simple point charge) water model. The counterions are added to the system to provide a neutral simulation system. The whole system was subsequently minimized, charges of the atoms of NAD^+^ and ALY were calculated by using OPLS2005 Force field. The charges of the atoms of NAD^+^ and ALY were calculated by using OPLS2005 Force field. Covalent and non-bonded parameters for the atoms in the ligands were assigned, by analogy or through interpolation, from those already present in the OPLS2005 force field. MD simulation is then carried out using the Desmond package (version 2.3) with constant temperature and pressure (NPT) and periodic boundary conditions. The OPLS2005 force field was applied to the proteins. The default Desmond minimization and equilibration procedure was followed. Simulations were kept at constant pressure (1 bar) and temperature (300 K) maintained with a Berendsen barostat and thermostat, respectively. SHAKE was applied to all systems allowing a 2-fs time-step. Long-range interactions were treated with the Particle Mesh Ewald method for periodic boundaries using a nonbonded cut-off of 9.0 Å and the nonbonded list was updated frequently using the default settings. Coordinates and energies for the OPLS2005 force field simulations were recorded for total of 15ns. The structure of the equilibrium state (15ns) was extracted from the MD. Het molecules except for native ligands were eliminated (water and counter ions for MD). Minimization was performed using the OPLS3e force field to produce the MM-optimized reactant structure.

#### QM-MM optimization of the SIRT3 complex

The minimized SIRT3-ALY-NAD^+^ complex structure optimized using the OPLS2005 force field was further optimized at the QM/MM (DFT-B3LYP/6-31G*: OPLS2005) level, using the QSite module as implemented in Schrodinger Suite. The quantum mechanics (QM) region of reactant includes the NAD^+^, acetyl lysine (entire residue of the peptide), Gln 228 (entire residue of the receptor) and side chain (CH2C3N2H3) of His 248, for a total of 132 atoms. The QM region was calculated by using the density functional theory with the B3LYP exchange-correlation functional and 6-31G* basis set. The remainder of the system (MM region) was treated by using the OPLS2005 force field. A total of 4023 atoms in the system were included in QM/MM simulations. To avoid over polarization of QM atoms by MM atoms at the boundary, Gaussian charge distributions were used to represent the potential of the atoms within two covalent bonds of the QM/MM cut-site using the Gaussian grid method for hydrogen cap electrostatics in QSite. MM point charges were employed for the rest of the MM region.

### Protein-ligand docking

Molecular docking of compounds to SIRT3 ligand complexes was carried out using AutoDockVina as previously described (*30*). AutoDockTools was used to prepare the Sirt3 receptor in its native format by adding polar hydrogens and assigning appropriate Gasteiger charges to all atoms as well as identifying the coordinates of the target box for enclosing the QM/MM optimized SIRT3.Ac-Pr.NAD^+^ substrate for global docking with a cubic grid box of 126×126×126 point dimensions of 0.5 resolution. Prior to docking, the compounds were geometry optimized in gas phase by Hartree-Fock method using the 6-311G+ basis set.

### Loop modeling and binding affinity calculations

Modelling of residues in the co-factor binding loop region that were missing from crystallographic data was carried out using Random Coordinate Discrete (RCD+) method implemented in the Fast Loop Modelling server that applies ab initio modelling with coarse-grained conformational search (*52, 53*). The loops generated were subsequently ranked using fast distance-orientation dependent energy filter and are afterwards refined with the Rosetta energy function to obtain accurate all-atom predictions to the models. An ensemble of closed loops up to 10,000 loops for the residues from Ser159 – Leu168 are generated after specifying the anchor residues, using coarse-grained contact potential (*54*) that associates inter-residue distance and orientation applied in this method. Native loop prediction scenario was selected to model the missing region of the loop. The lowest energy loop model is selected as the final structure among top 20 models obtained as the result.

For modeling of the effect of alternate conformations of loops on ligand binding affinities, loop replacement was carried out using the following steps as previously described (*30*):

### Sequence-structure based superimposition

- Graft the coordinates for the co-factor loop region.
- Run protein preparation wizard on the modelled product complex (open/closed loop conformation).
- Predict/repack the side chains for all residues within 7.5 Å of the grafted loop region using Monte Carlo approach together with backbone sampling.
- Carry out prime energy refinement only on those residues which were repacked keeping the others fixed.

### Molecular dynamics simulation for conformational and binding affinity analyses on protein-ligand complexes

#### Force field

The Amber99SB force field (*55, 56*) was used for all the molecular mechanics calculations, as previously described (*30*). Extra parameters were adapted from the following sources: parameters for Zn developed by Luo’s group (*57*); parameters for NAD^+^ developed by Walker et al (*58*) and Pavelites et al (*59*) ; parameters for acetylated lysine developed by Papamokos et al (*60*). Amber14 tools (Antechamber program) are employed for parametrizing the non-standard residues and tleap program for creating topology and force field parameters for the standard residues.

#### Fast long-range electrostatics

Electrostatic energy was calculated using the Particle Mesh Ewald (PME) method (*61*).

#### Explicit vs implicit solvation

Explicit solvent was employed for sampling, but implicit solvent was used for subsequent energy calculations. Water molecules were stripped from the molecular dynamics trajectories in preparation for MM/PBSA and MM/GBSA re-runs.

#### MD simulation

All molecular dynamics simulations were performed with periodic boundary conditions to produce isothermal-isobaric ensembles (NPT) at 300 K using the NAMD program (*62*), as previously described (*30*). The integration of the equations of motion was conducted at a time step of 2 femtoseconds. The covalent bonds involving hydrogen atoms were frozen with the SHAKE algorithm (*63*). Temperature was regulated using the Langevin dynamics with the collision frequency of 1 ps^−1^. Pressure regulation was achieved with isotropic position scaling and the pressure relaxation time was set to 1.0 picosecond.

### High-throughput batch docking

A database containing 1.2 million compounds (Chembridge) was screened against two SIRT3 structures (PDB: 4FVT and 4BVG) (*64, 65*). Our prior knowledge of the bioactive conformations of HKL and its binding sites in SIRT3 were applied to define the docking site and the selection of the protein conformation. The structure-based docking was used in a sequential process to implement VS (**Scheme 1**). In brief, these 1.2 million compounds were first imported into a database and converted from 2D ⭢ 3D using relevant modules in MOE. Assign partial charges and perform a quick round of energy minimization using the MMFF94x forcefield. When appropriate, the most likely tautomeric form was generated and automatically chosen.

These prepared ligand structures were converted back into the input file format preferred by AutoDock Vina. Dockings were performed using AutoDock Vina on an AWS cloud server. The grid center and volume were defined as follows. For 4FVT, Center (x, y, z) = 23.19, 31.19, −14.31; Volume =29.6 × 34.8 × 33.8. For 4BVG, Center (x, y, z) = 22.24, 30.39, −7.98; Volume = 25.0 × 25.0 × 25.0. 20 possible top conformations for each ligand were collected.

### Post-docking analysis

The docked ligands were then ranked by docking scores in AutoDock Vina (predicted affinity in kcal/mol). The top 1,000 compounds were then screened for (1) non-covalent interactions with the residues in the active site of 4FVT and 4BVG; (2) formation of H-bonds with residues lining binding site; (3) interaction within specific residues within the co-factor binding loop of SIRT3; (4) predicted ADME properties, such as LogP, LogD and (5) predictions about synthetic feasibility. Only the best ranked pose of each compound from the top 1,000 were included in the list.

This list of top 1,000 compounds was then subjected to further post-docking analysis using MOE. The ligands were imported into a database and converted into the native format utilized by MOE after addition of hydrogen atoms and recalculation of partial charges for each structure. A small percentage of these structures were carefully examined to ascertain that this processing had not introduced any undesirable artifacts in them. Next a truncated post-docking energy minimization of ligands and hydrogen atoms from adjacent residues was performed to further optimize the interactions between docked poses and site residues, using a MOE script known as ‘analysis_dock.svl’. The energy optimization step provides a new value for approximate calculated Binding Energy of the compounds (in MOE). The optimized structures were then used for all subsequent post-docking analysis steps.

We screened for the presence of H_Bonds between optimized poses and residues lining the binding site. Based on the previously docked structure of Honokiol, the following residues were considered to be part of the site: 4FVT (Gly 145, Ser 149, Gly 153, Gly 163, Leu 168, Tyr 171, Phe 180, Lys 195, Glu 198, Tyr 204, Gln 228, and His 248) and 4BVG (Asp 156, Pro 176, Ile 179, Phe 294, Ser 321, Glu 323, and NLE7 from the co-crystallized Ac-ACS peptide chain). The docked pose of Honokiol was also used to measure the relative ‘receptor buriedness’ of the optimized poses in both receptor structures. The BHB (buriedness, hydrogen bonding, and binding energy) score for each compound was calculated using the general formula first described by Feher M et al (*66*).

### Evaluating the agreement of the Model to the X-ray crystallography data

After docking of the ligands in the binding pocket, the density map was searched for clusters of density that might contain a ligand. Careful inspection of the model against the map was performed using Coot molecular graphics software (*67*). This program was used to optimize the ligand’s fit to the density and removing the atomic clashes in the model. The starting position of the docked ligand was transformed by applying a set of random rotations and small translations. The resulting positions were then submitted to rigid-body refinement and scored based on density fit and the best pose was real-space refined.

The quality of fitting of the refined model to the experimental X-ray crystallographic map was determined using the validation process of the structure. Assessing the fit of a model to the density is affected by factors like map noise and resolution of the data. Although factors like R_free_ (*68*) provide reasonable measurements of overall bias of the model, for validation of the ligands fit to the map we need local indicators. Local parameters like real-space residual (RSR) (*69*) or real-space correlation coefficient (RSCC) (*70*) are more useful for such evaluations (**Eqs. 4** and **5**).

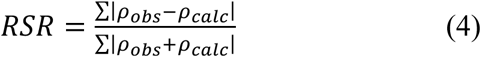

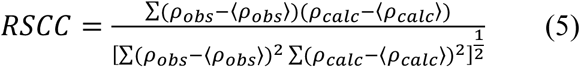

Where *ρ*’s are density values at grid points that cover the ligand, obs and calc refer to experimental and model densities respectively. The atomic coordinates and reflection map were deposited in wwPDB validation server (will be released with PDB IDs).

With the ligand’s coordinates, to make sure that the chemistry is reasonable based on similar existing structures in wwPDB Chemical Component Dictionary, Molecular Geometry and Unified Library (Mogul) (*71*) analysis was performed. Two NAD^+^ ligands were matched and HKL could not be matched to an existing wwPDB chemical component dictionary definition. Two-dimensional graphical depiction of Mogul quality analysis of bond lengths, bond angles, torsion angles, and ring geometry for two NAD^+^ ligands are shown in **Table S2**. Based on wwPDB validation report criteria, for torsion angles, if less than 5% of the Mogul distribution of torsion angles is within 10 degrees of the torsion angle in question, then that torsion angle is considered an outlier. Any bond that is central to one or more torsion angles identified as an outlier by Mogul is highlighted in the graph. For rings, the root-mean-square deviation (RMSD) between the ring in question and similar rings identified by Mogul was calculated over all ring torsion angles. If the average RMSD was greater than 60 degrees and the minimal RMSD between the ring in question and any Mogul-identified rings was also greater than 60 degrees, then that ring was considered an outlier. The outliers are highlighted in purple. The color gray indicates Mogul did not find sufficient equivalents in the Cambridge Crystallographic Data (CSD) to analyze the geometry.

### Generating ROC curve of 1.2 million compounds docking scores

For generating the ROC curve the following data were processed:

- Docking scores for 4BVG receptor including about 1.2M docked compounds.
- Docking scores for 4FVT receptor including about 1.2M docked compounds.
- Docking scores for a subset (68 compounds) of the above 1.2 million compounds against two above receptors, as well as modulation effects on SIRT3 of these compounds, as determined experimentally with the HPLC assay.

The experimental HPLC scores for the 68 compounds were obtained. Based on these data, we determined if a compound is a SIRT3 modulator or not. As shown in **Table S4**, a SIRT3 modulator is one where the HPLC score is above a threshold in absolute value, i.e. |HPLC-100|>20.

For these 68 compounds, we determined the % of modulators (positives) in three bins based on the docking scores. The percentages of modulators for these three bins are 21.1%, 56.3% and 39.4% respectively (**Table S5**). The probability of each of the 1.2M compounds being a positive was set to the respective percentage corresponding to the bin into which its docking score fell. TPR and FPR for the range of cutoffs of the docking score were calculated (docking score refers to the better score between the two receptors).

## Supplementary Information

**Fig. S1.**
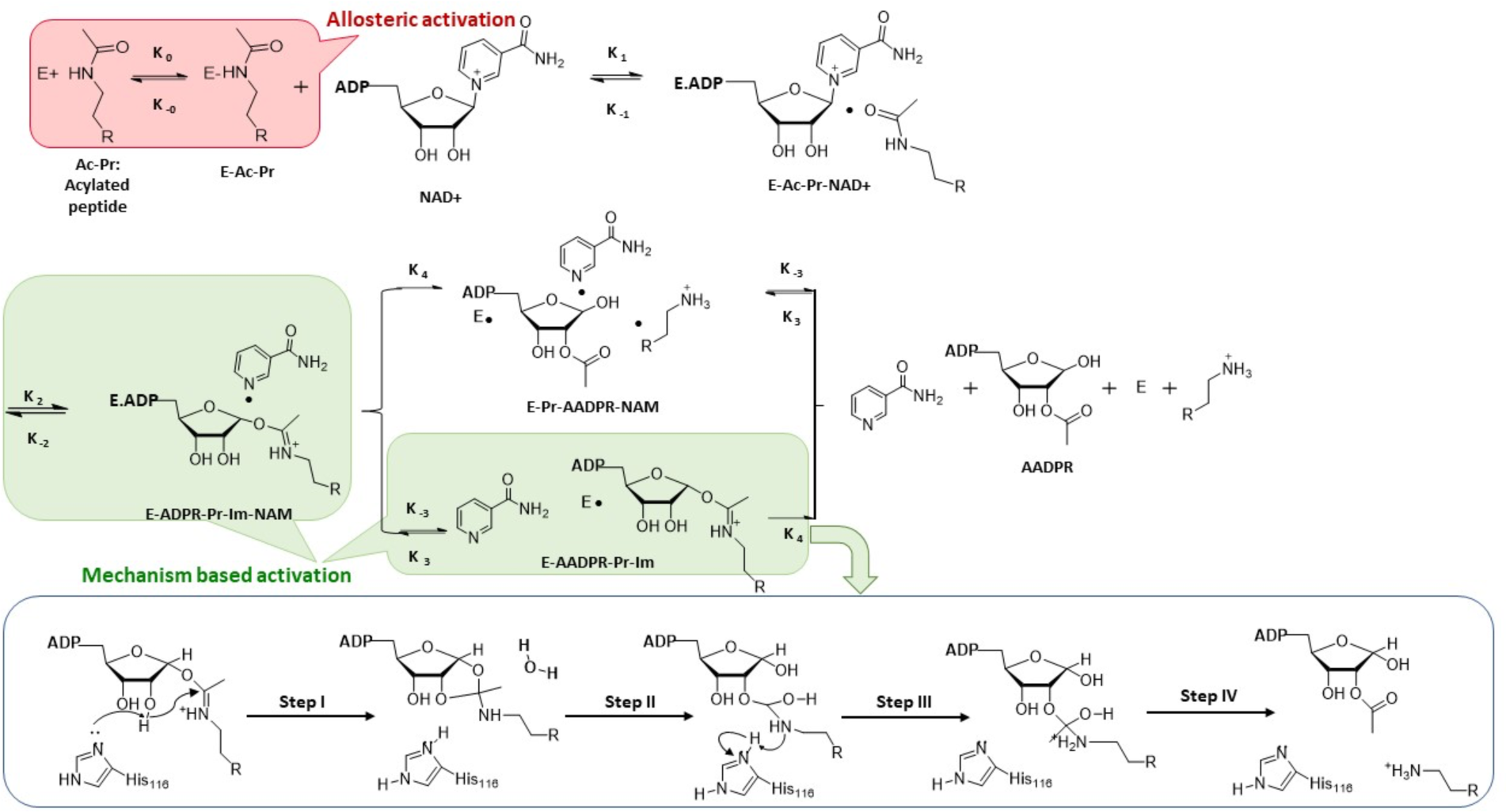
Chemical mechanism of sirtuin-catalyzed deacylation and modes of sirtuin activation (*1*).

**Fig. S2.**
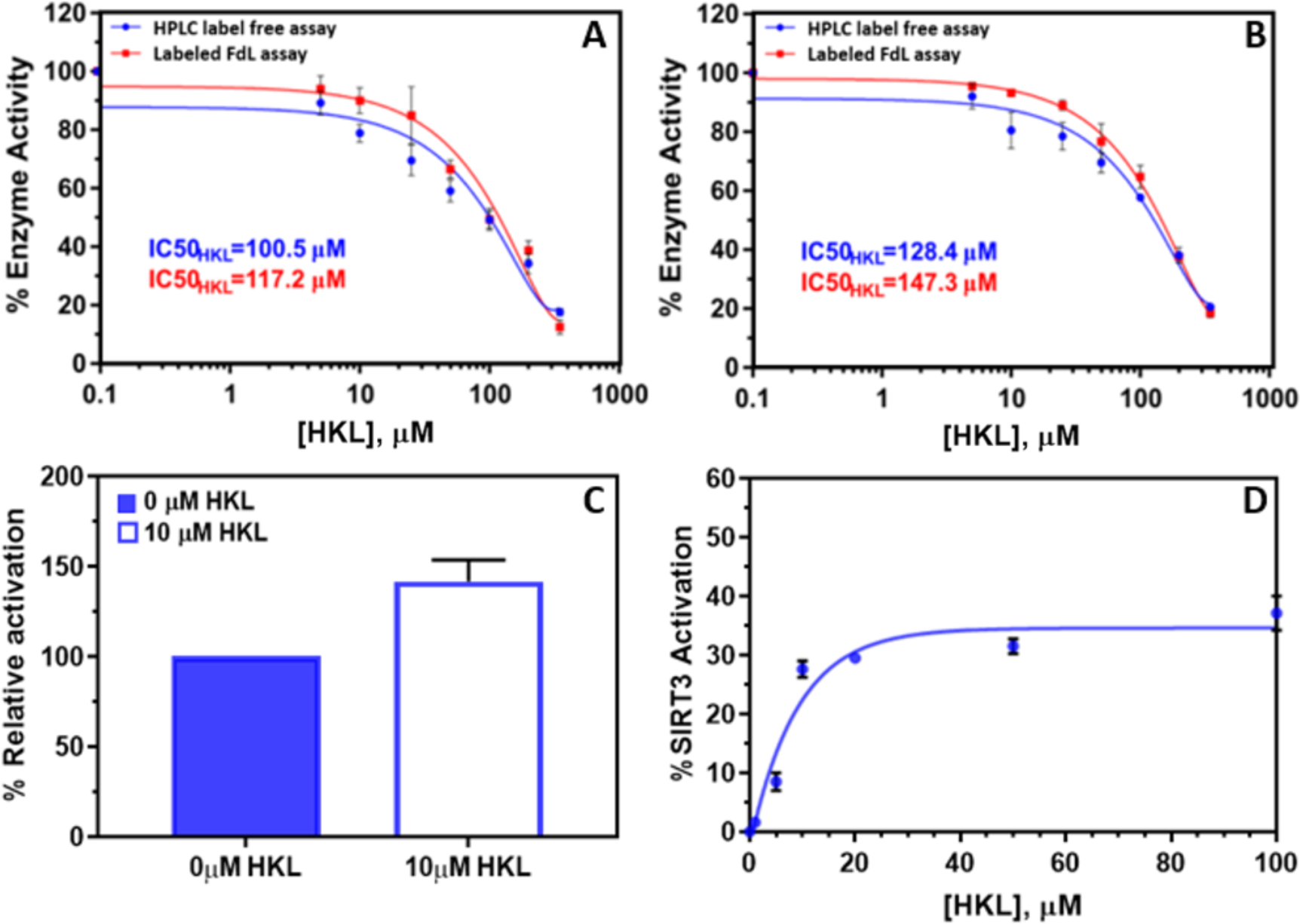
Effect of HKL on Sirt3 deacetylation activity for FDL (fluorophore-labeled p53-AMC peptide) substrate. **(A, B)** Dose-response curves were measured under conditions where [E]_0_/[S]_0_ << 1, where [S]_0_ denotes the initial concentration of the limiting substrate, in order to maximize the contribution, the steady state phase of the reaction to the curve. **(A)** Saturating FDL (250µM) and non-saturating NAD^+^(25µM) (N=2); **(B)** saturating NAD^+^ (3000µM) and non-saturating FDL (30µM) (N=2) in the presence of different concentrations of HKL range from 0-350 µM using label-free HPLC (red) and labeled FDL assay (blue). Reactions were terminated after 30 min. **(C)** Activation of SIRT3 with p53-AMC substrate in the presence of 10 mM HKL at [E]_0_/[NAD^+^]_0_=0.0175 ([E]_0_=0.349 µM, [NAD^+^]_0_=20 µM, [p53-AMC] _0_=10 µM, t=5 min, N=2. **(D)** p53-AMC substrate deacetylation measured under higher enzyme concentration conditions, [E]_0_/[NAD^+^]_0_=0.3, in the presence of 50 µM NAD^+^ and 10 µM FDL, t=5 min (N=3). The error bars that are not visible are too small to view at this scale.

**Fig. S3.**
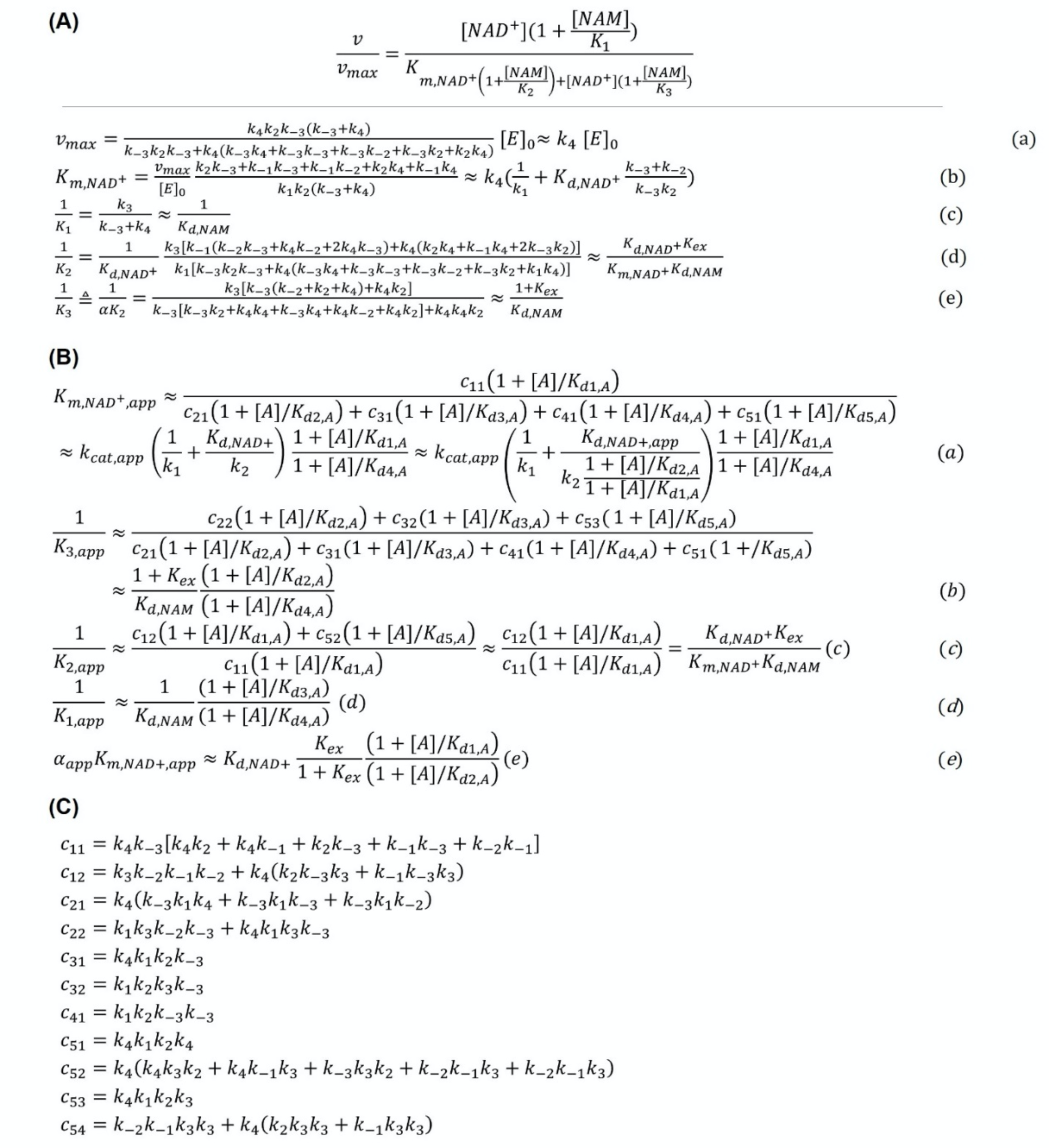
(A) Steady state parameters of sirtuin-catalyzed deacylation as functions of rate constants in the reaction mechanism. (*1*). *K_ex_* = k_-2_/k_2_, and the approximations refer to the case where *k_4_*≪*k_j_, j≠* 4. **(B)** Expressions for **(A); (C)** Expressions for c_ij_’s in **(B).**

**Fig. S4.**
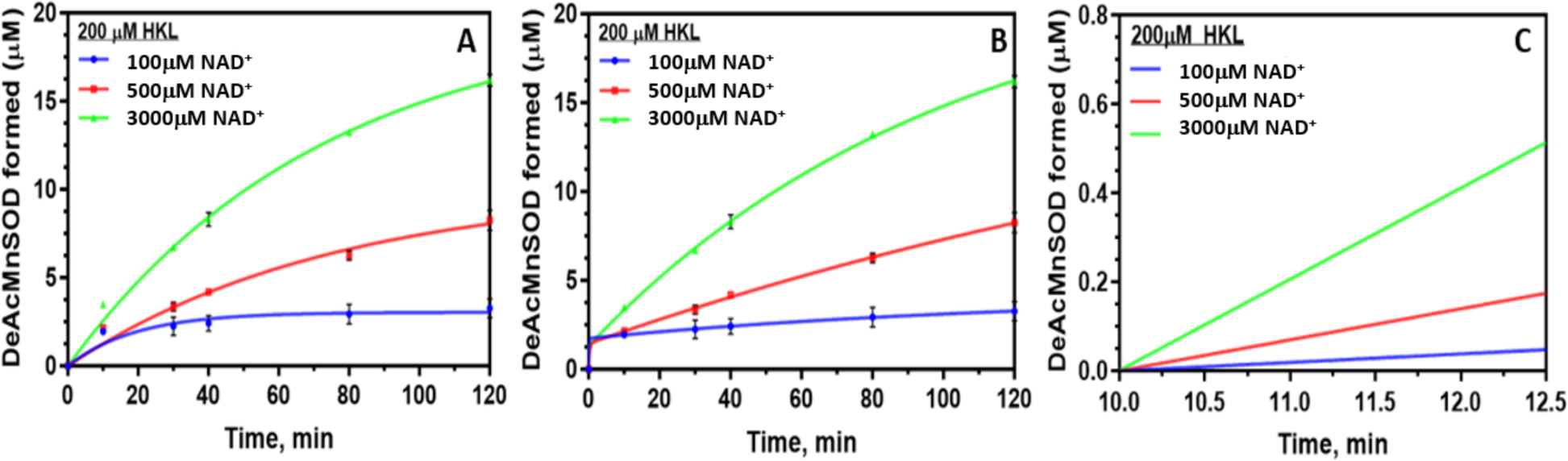
Sirt3 deacetylation activity in the presence of HKL for different cofactor (NAD^+^) concentrations: steady state rate determination. Plots of DeAc-MnSOD formed vs. time with **(A)** single exponential fitting, and **(B)** double exponential fitting. Corresponding linear rate fitting plots for **(C)** double exponential (steady state rates) starting at 10 min. [NAM] = 100µM. Note 100 µM NAM is on the same order of the magnitude as the physiological concentration of NAM.

**Fig. S5.**
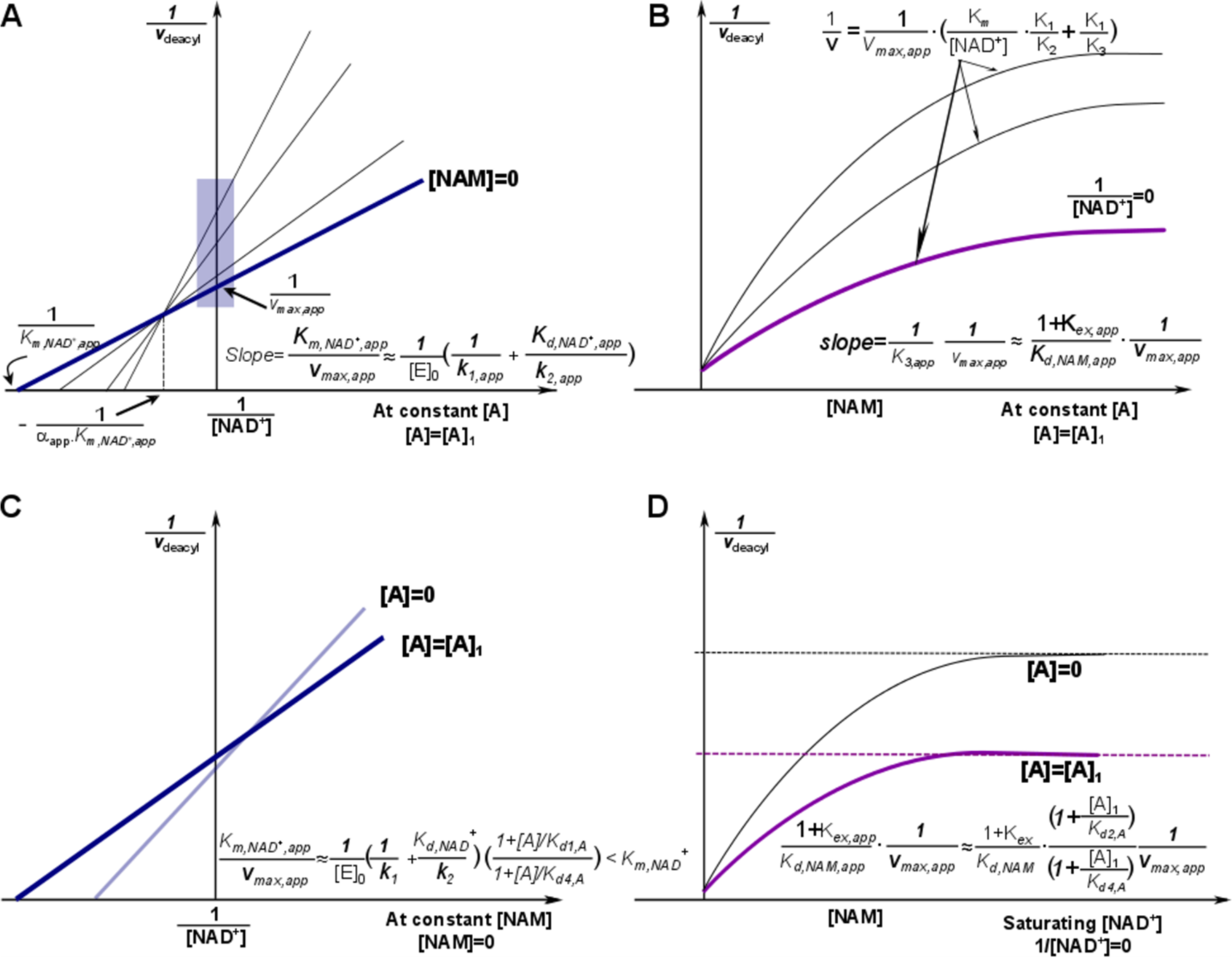
Mechanism-based activation of sirtuin enzymes: model-predicted steady state and dose-response properties (*1*). **(A)** Double reciprocal plots for deacylation initial rate measurements in the presence of activator. Data used to construct the Dixon plot at saturating [NAD^+^] depicted in (B) are highlighted by the blue box on the y axis. **(B)** Dixon plots for initial rates of deacylation in the presence of activator. Predicted plateaus in these curves are indicated by the arrows. **(C)** Comparison of double reciprocal plots at [NAM] = 0 µM with and without activator. **(D)** Dixon plots at 1/[NAD^+^] = 0 compared with and without activator. “A” indicates a mechanism-based sirtuin activating compound. The slight increase in k_cat_ implied in C is not a consequence of the model predictions (*1*) and is shown to account for the possibility of activation at low [NAD^+^] irrespective of whether such an effect occurs.

**Fig. S6.**
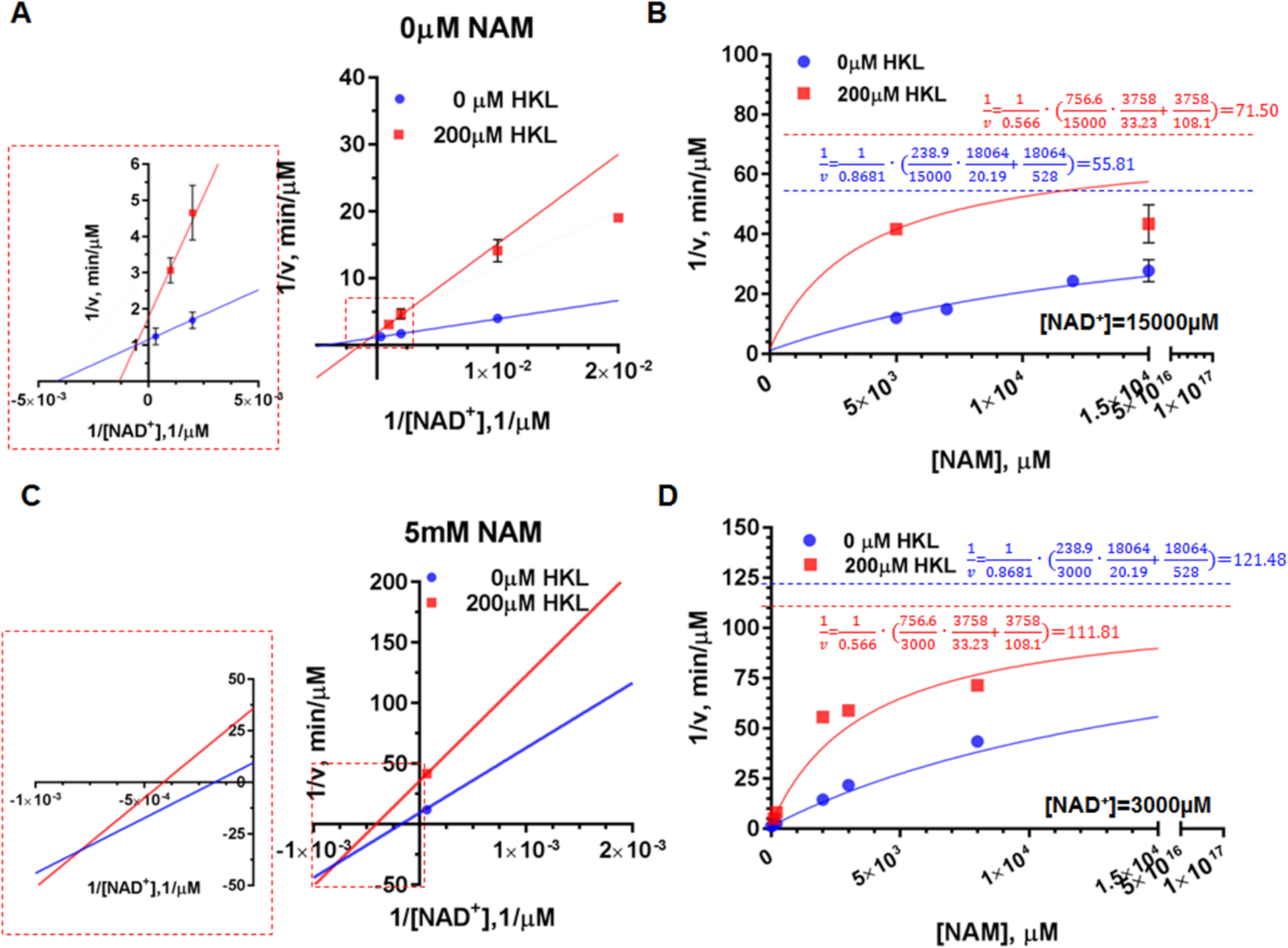
Mechanism-based modulation of hSirt3^102-399^ deacetylation of MnSOD substrate by HKL: effect of NAD^+^ and NAM. **(A)** Double reciprocal plots for deacetylation initial rate measurements in the presence and absence of HKL at [NAM] = 0 µM. The enlargement of intersection point is provided as inset. **(B)** Comparison of Dixon plots at [NAD^+^] = 15000 µM in the presence and absence of HKL. **(C)** Double reciprocal plots for deacetylation initial rate measurements in the presence and absence of HKL at [NAM] = 5mM. The enlargement of intersection point is provided as inset. **(D)** Comparison of Dixon plots at [NAD^+^] = 3000 µM in the presence and absence of HKL. Note that the model omits the term which is quadratic in [NAM] in the steady state model, as is standard, and the dotted lines in the Dixon plots represent the asymptotic values that the model-predicted rates converge to if this term is omitted. **Fig. S5** depicts the corresponding predicted plots for a mechanism-based steady state activator (HKL is a non-steady state activator).

**Fig. S7.**
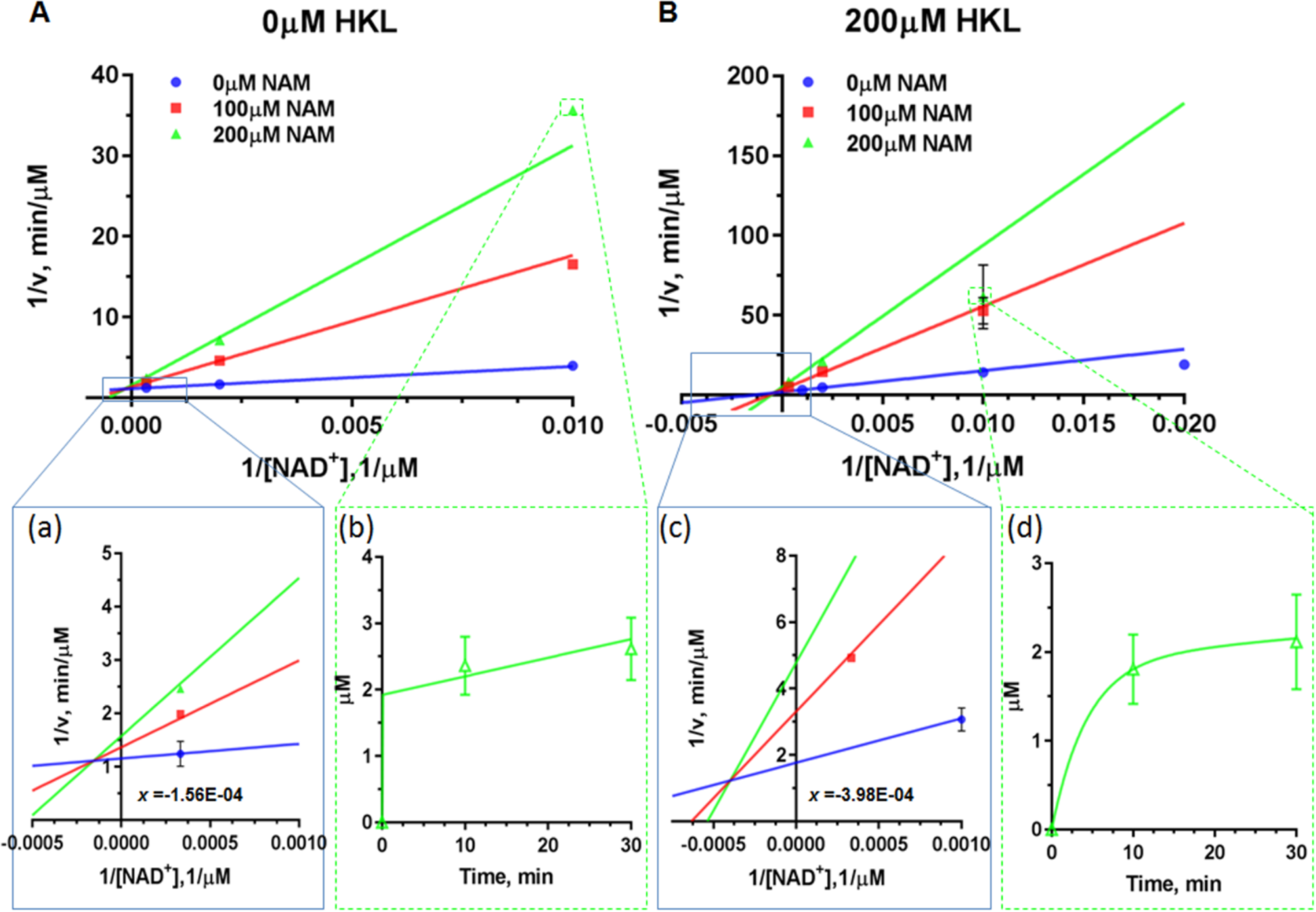
Steady state kinetic characterization of deacetylation in the presence and absence of the non-steady state activator HKL. Double reciprocal plots for deacetylation initial rate measurements of saturating substrate peptide (MnSOD K122) in the presence of different concentrations of NAM with **(A)** 0 µM HKL; **(B)** 200 µM HKL. Enlargements of the intersection points are provided as insets (a) and (c). The time series plots of µM product formed vs. time (for selected small time points; see **Fig. S4** for full time course) are provided as insets (b) and (d). The lines are the results of global fitting; the lines were not fit independently to the data points in the individual panels. The error bars that are not visible are too small to view at this scale. The insets in **Fig. S7** show examples of the time series data collected. These insets demonstrate that deacetylation of this substrate displays a two-phase behavior (pre-steady state/steady state phase) with a relatively slow first phase. Due to this behavior, as noted above we used a two-phase rather than one-phase exponential time series fitting. Irrespective of the first phase time constant, the rate in the second phase is almost identical. Hence the uncertainty in steady state rates is very low. The rate is generally almost constant over the measured times in the second phase.

**Fig. S8.**
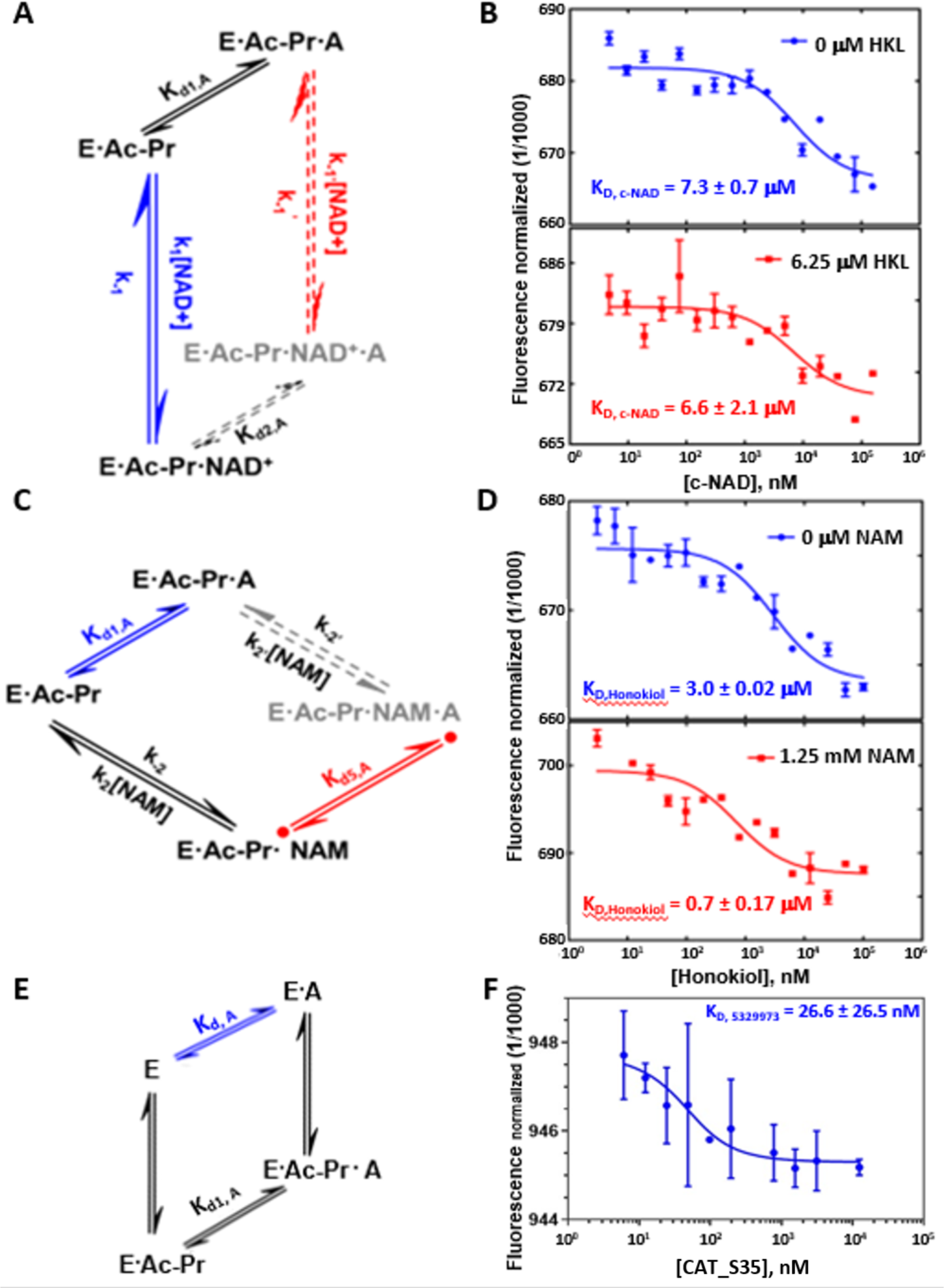
Binding affinity measurements for complexes in the sirtuin reaction mechanism. **(A, C, E)** Pathways in the sirtuin reaction network from **Fig. S11**. E, enzyme; Ac-Pr, acetylated peptide substrate; NAD, nicotinamide adenine dinucleotide; A, modulator (HKL or hit compound CAT_35); NAM, nicotinamide adenine mononucleotide. **(B)** Carba-NAD binding in the ternary complex: effect of mechanism-based modulator HKL. Binding of carba NAD (c-NAD) to Sirt3.Ac-MnSOD complex, in presence and absence of 6.25 µM HKL measured using MST. **(D)** HKL binding: effect of NAM. Binding of HKL to the Sirt3.Ac-MnSOD complex, in presence and absence of NAM, measured using MST. (**F**) Hit compound CAT_35 binding to the apo enzyme Sirt3.

**Fig. S9.**
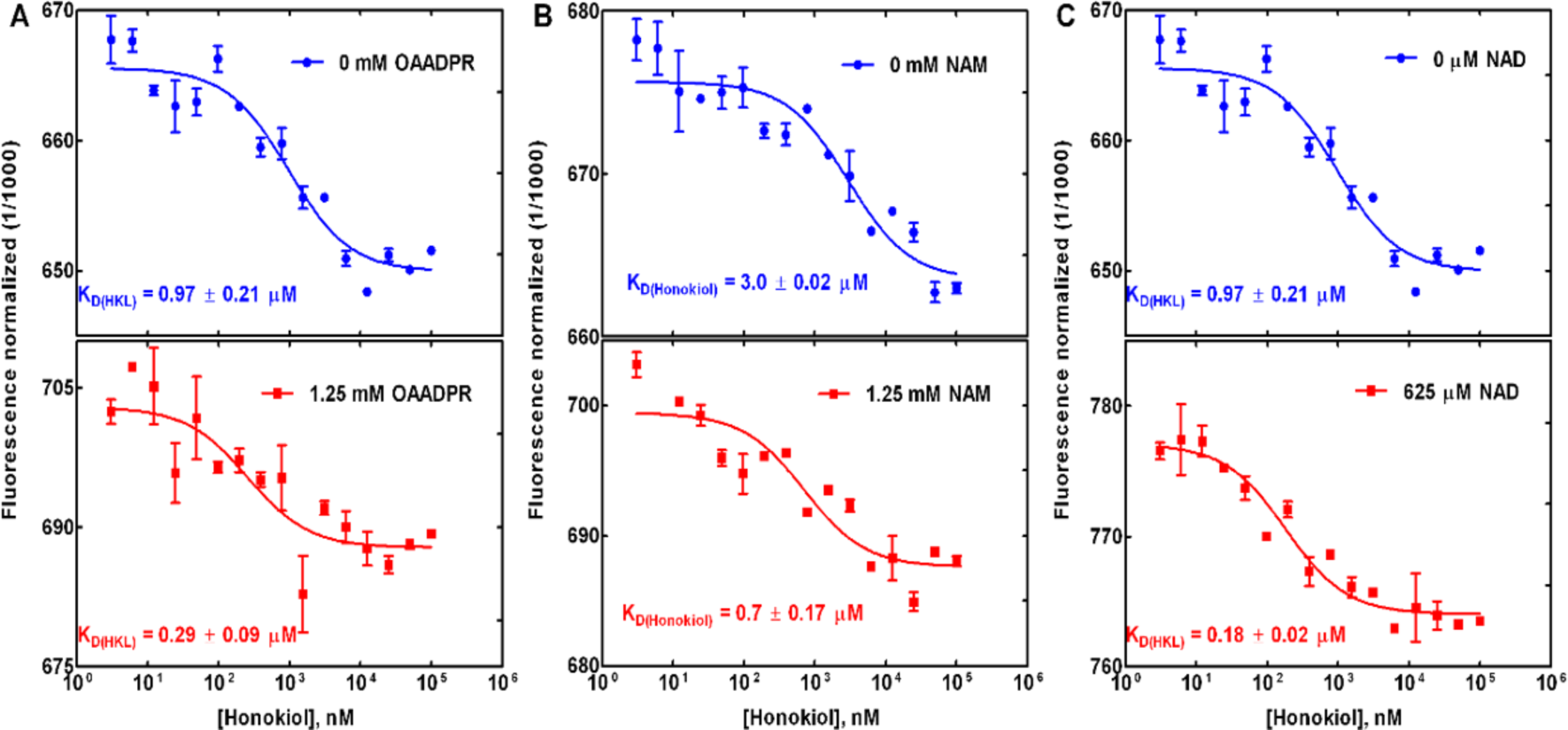
Binding affinity measurements of **(A)** Honokiol binding in the product complex: effect of 2’-O-acetylated ADP ribose coproduct; **(B)** Honokiol binding in the coproduct complex: effect of NAM; **(C)** Honokiol binding in the product complex: effect of NAD^+^. Note that measuring the binding affinities for any of the 3 sides of such faces determines the fourth binding affinity. In particular, the binding affinities of HKL or Carba-NAD were measured in the ternary complex and product complex. Cooperative binding between HKL and the relevant ligand were studied in each case. It was observed that HKL can have synergistic binding interactions with several ligands. For binding affinity studies on complexes that appear in the model schematic **Fig. S11**, the relevant faces of the cube are also displayed in the respective figures. Note that measuring the binding affinities for any of the 3 sides of such faces determines the fourth binding affinity. In particular, the binding affinities of HKL or Carba-NAD were measured in the ternary complex and product complex. Cooperative binding between HKL and the relevant ligand were studied in each case. It was observed that HKL can have synergistic binding interactions with several ligands.

**Fig. S10.**
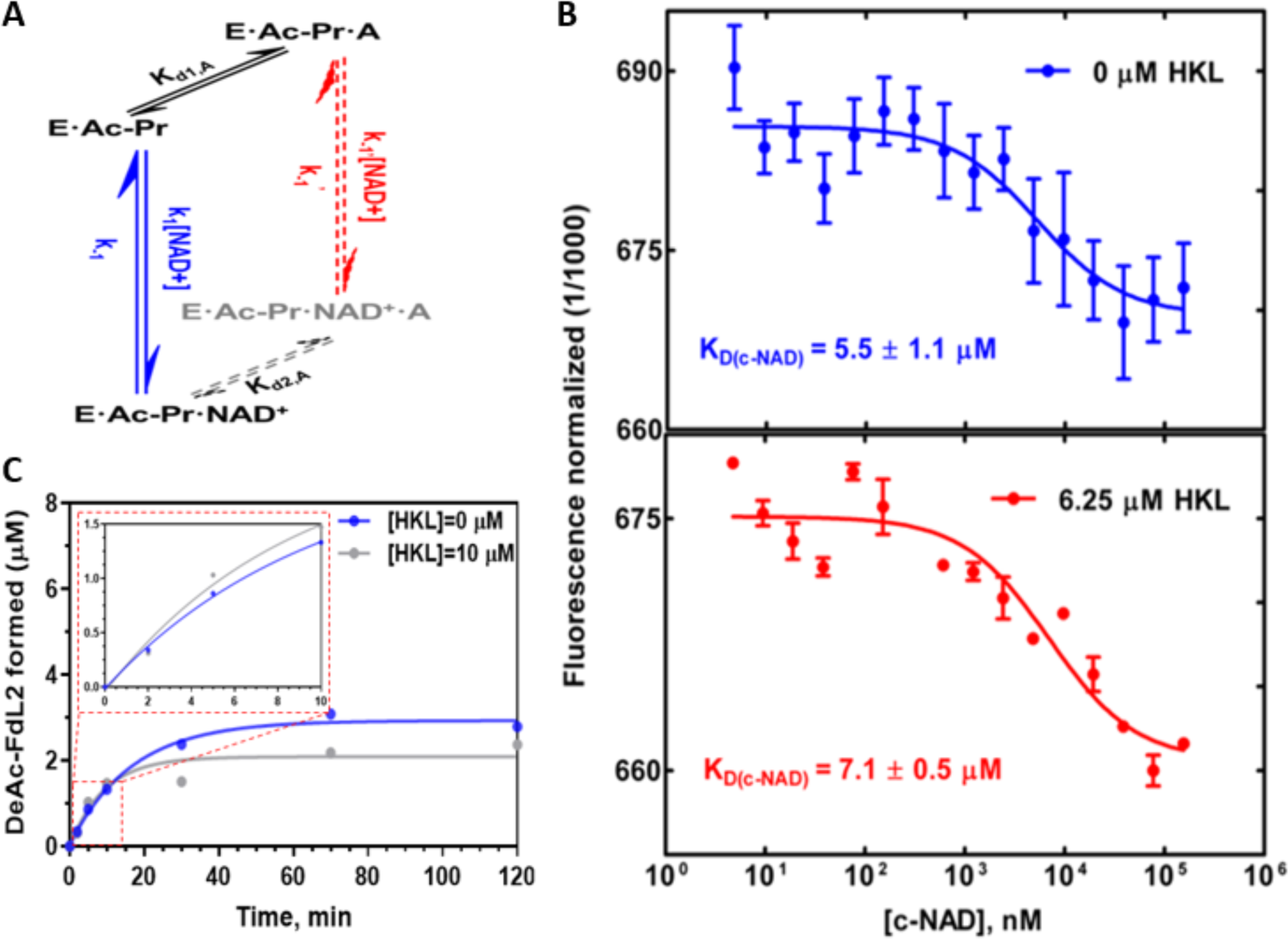
Effect of HKL on ligand binding affinity and deacetylation rate for the p53-AMC substrate. **(A, B)** Binding affinity measurements of Carba-NAD binding in the ternary complex p53-AMC substrate -- effect of HKL**. (A)** Pathways in the sirtuin reaction network from **Fig. S11**. E, enzyme; Ac-Pr, acetylated peptide substrate; NAD, NAM adenine dinucleotide; A, modulator (HKL). **(B)** Binding of Carba-NAD (c-NAD) to SIRT3.Ac-p53-AMC complex, in presence and absence of 6.25 µM HKL measured using MST. **(C)** SIRT3 activation by HKL: Plot of product formation vs. time ([NAD^+^]_0_ =50 µM, [FDL] = 10 µM, [E]_0_/[NAD^+^]_0_=0.3). As in the case of the MnSOD substrate, activation was observed under non-steady state condition and inhibition was observed under steady state condition. For the p53-AMC substrate, a standard mixed inhibition steady state model was fitted instead of **Eq. (1)** since high values of [NAM] were not sampled. **Table 4** summarizes all the *K_d_* values reported here.

**Fig. S11.**
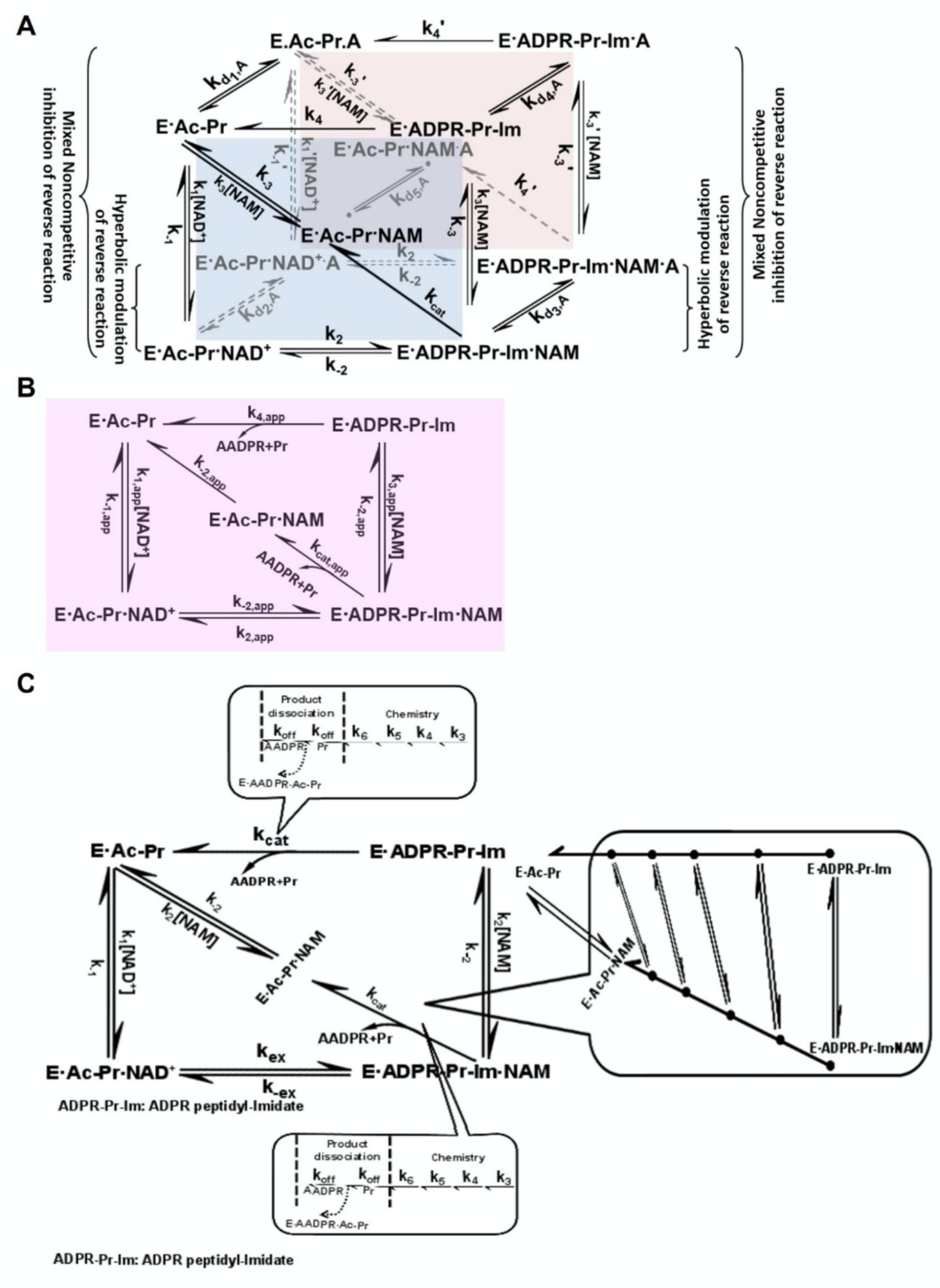
General model for mechanism-based sirtuin enzyme activation (*1*). **(A)** The front face of the cube (blue) depicts the reaction diagram for sirtuin-catalyzed deacylation in the presence of cofactor and nicotinamide concentrations [NAD^+^] and [NAM]. “A” denotes activator. The back face of the cube (pink) depicts the corresponding reaction diagram in the presence of saturating [A]. Conventional modes of enzyme activity modulation are annotated in the figure to elucidate the relationship between mechanism-based enzyme activation and these conventional modes. **(B)** At any [A], there exist apparent values of each of the rate constants in the reaction mechanism, producing the apparent reaction diagram (purple). These are denoted by “app” in the Figure. There are also corresponding “app” values of the steady state, Michaelis, and dissociation constants. **(C)** Extended model for sirtuin catalyzed deacetylation including separate representation of elementary deacetylation chemistry and product release steps.

**Table S1.** Carba-NAD and HKL ligands structure quality statistics after structure refinement fit to 2.4Å resolution crystal structure of human SIRT3 (HKL-SIRT3-2.4Å). The fit of the model to the electron density was quantified by the RSCC and RSR values. These values were obtained from the wwPDB validation report.

| Refinement statistics | RSCC value | RSR value |
| --- | --- | --- |
| Carba-NAD (chain A binding pocket) | 0.66 | 0.51 |
| Carba-NAD (chain C binding pocket) | 0.69 | 0.44 |
| HKL (chain C binding pocket) | 0.73 | 0.47 |

**Table S2.**
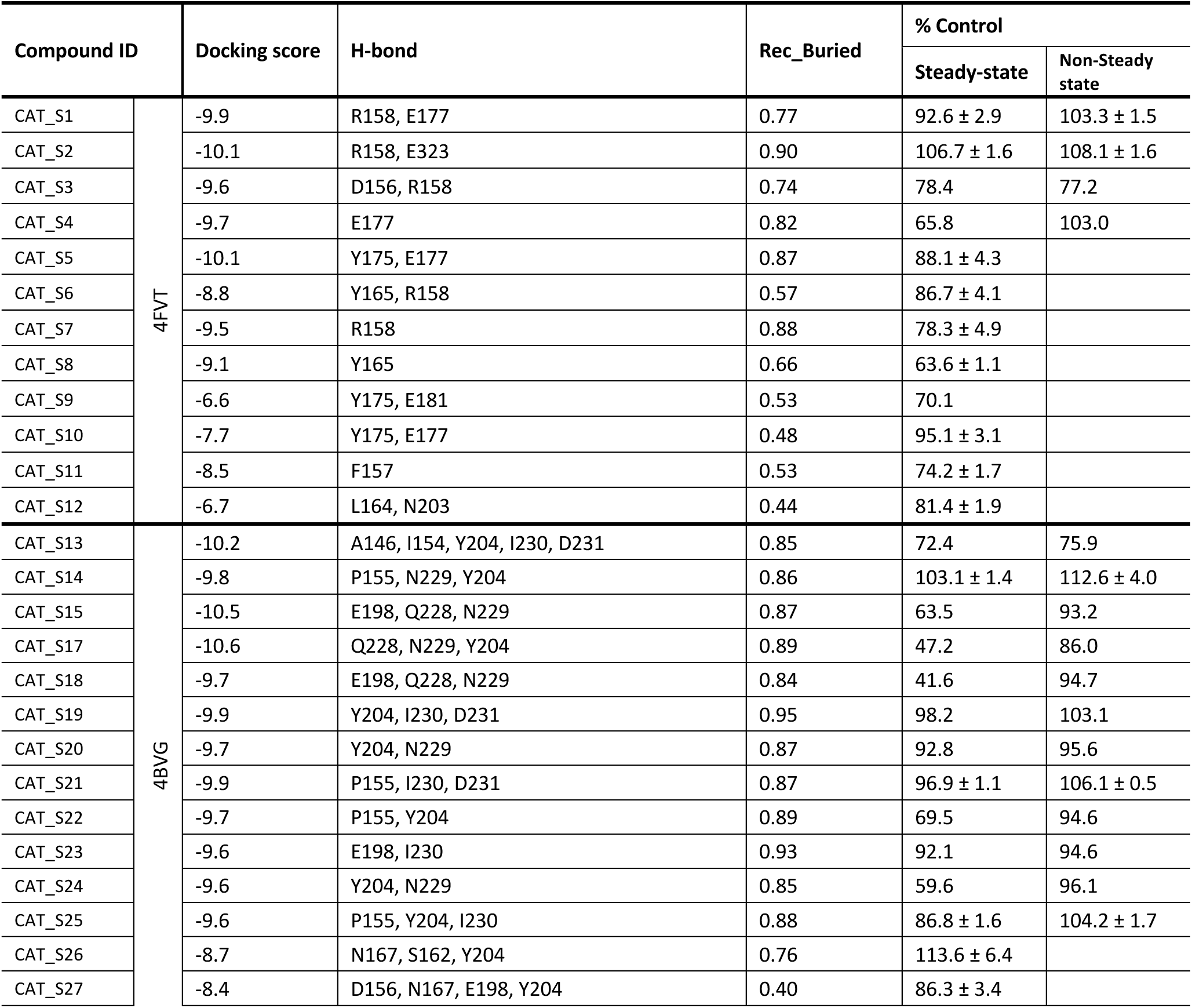
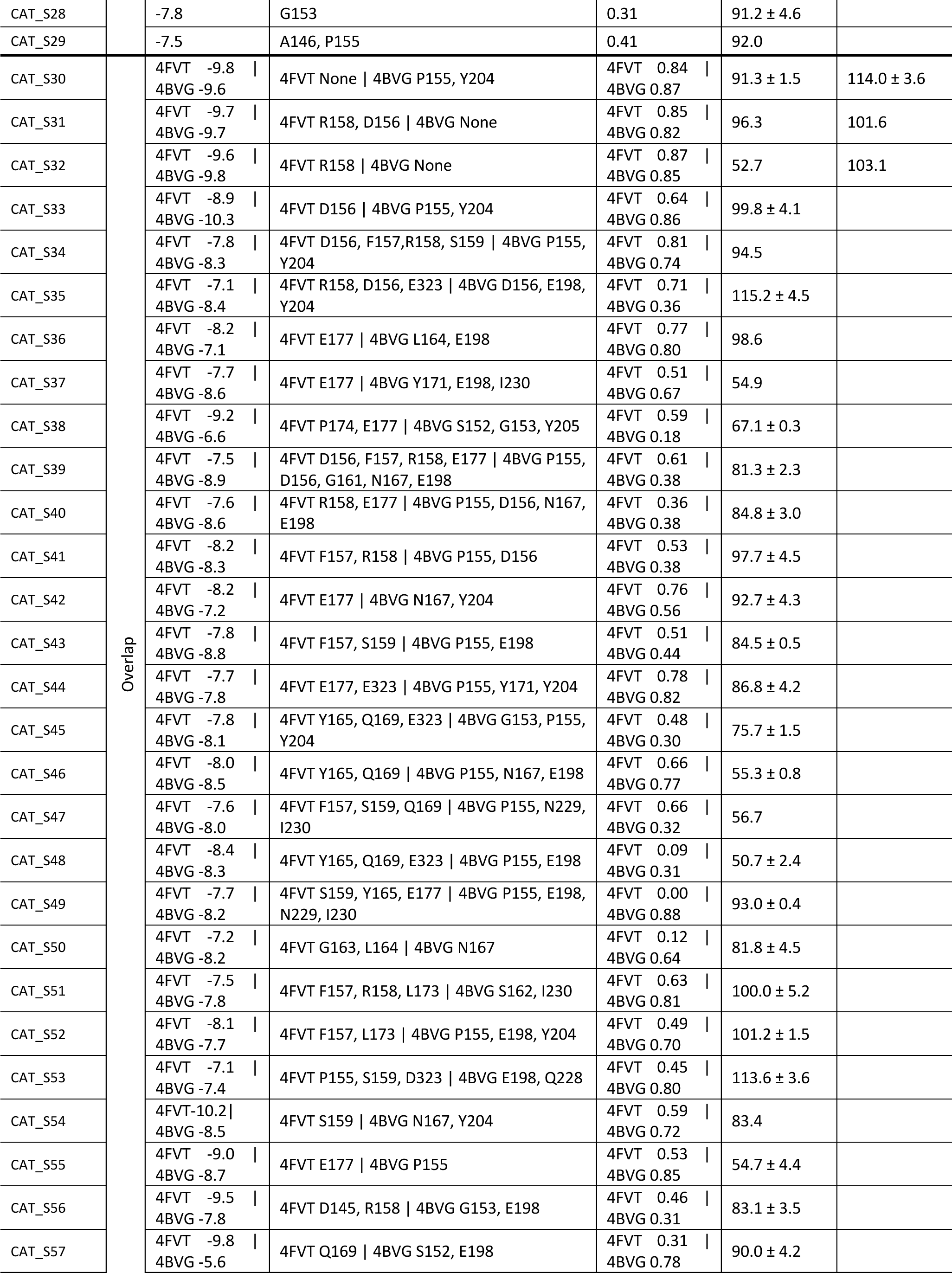
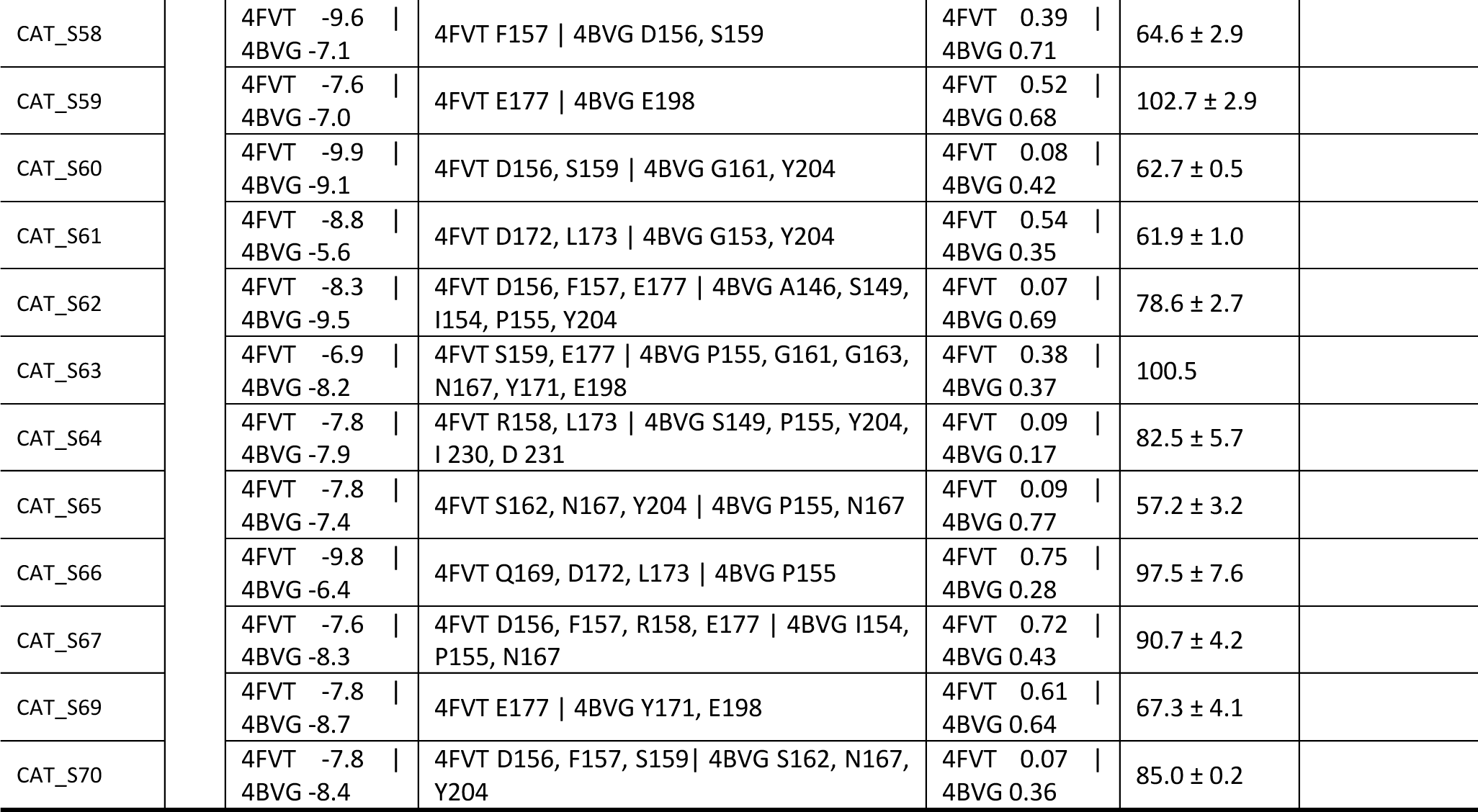
Docking scores of, and *in vitro* Sirt3 modulation by, the top 68 compounds selected through virtual screening. 1.2 million compound library were docked against two QM-MM geometry optimized SIRT3 complexes (4FVT and 4BVG) using AutoDock Vina. The docking scores, H-bond interaction, and Rec_Buried scores are presented. The modulation effects of top hits were validated and % activities are reported.

**Table S3.**
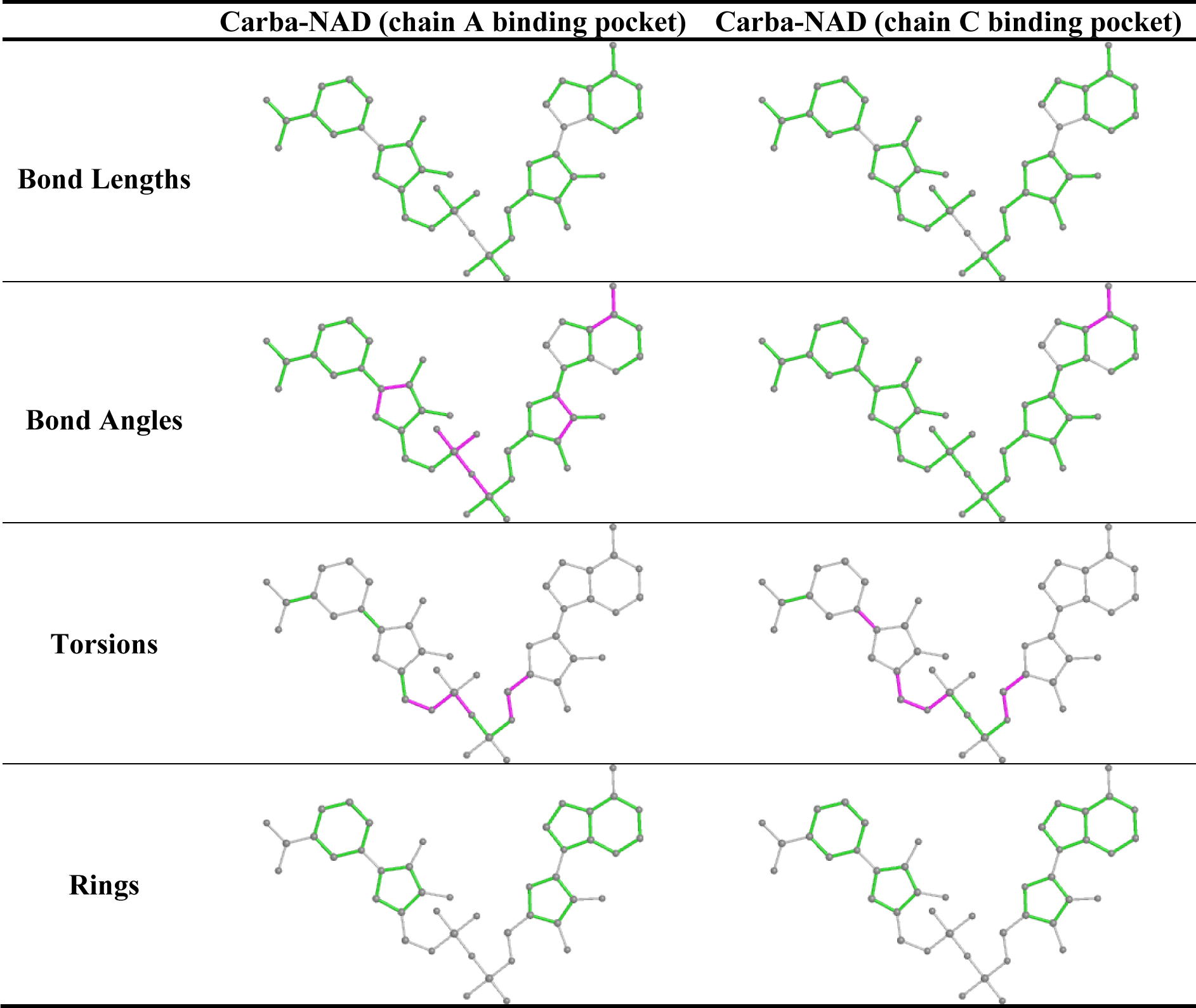
Two-dimensional graphical depiction of Mogul quality analysis of bond lengths, bond angles, torsion angles, and ring geometry for 2 Carba-NAD ligands in the chain A and chain C binding pockets.

**Table S4.**
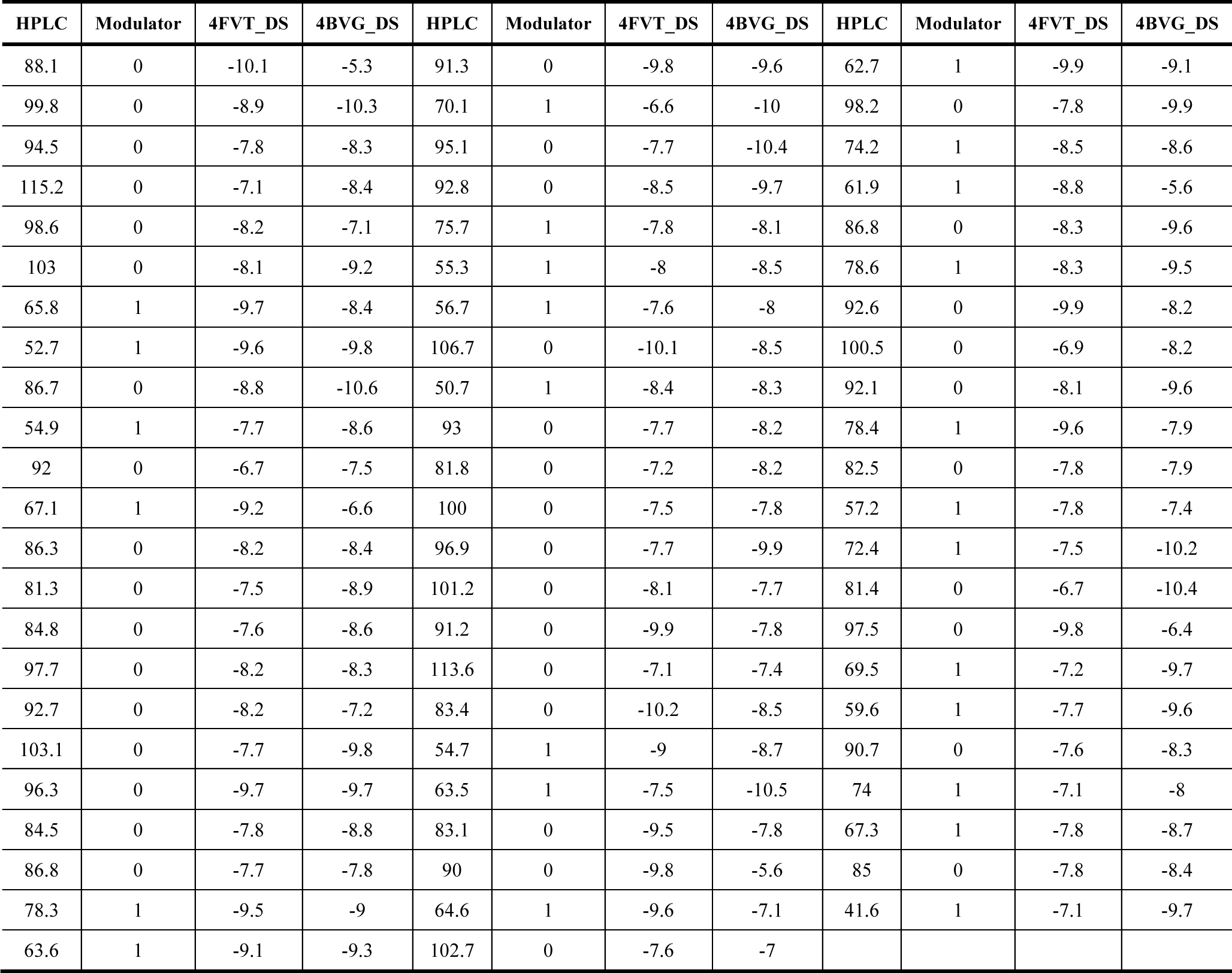
The modulation effect of two sets of docking scores data (4FVT_DS and 4BVG_DS) Modulator index takes the value 1 if the compound is a SIRT3 modulator and 0 otherwise. A SIRT3 modulator is one where the HPLC score is above a threshold in absolute value, i.e. |HPLC-100|>20. The HPLC data are only available amongst the 68 compounds.

**Table S5.**
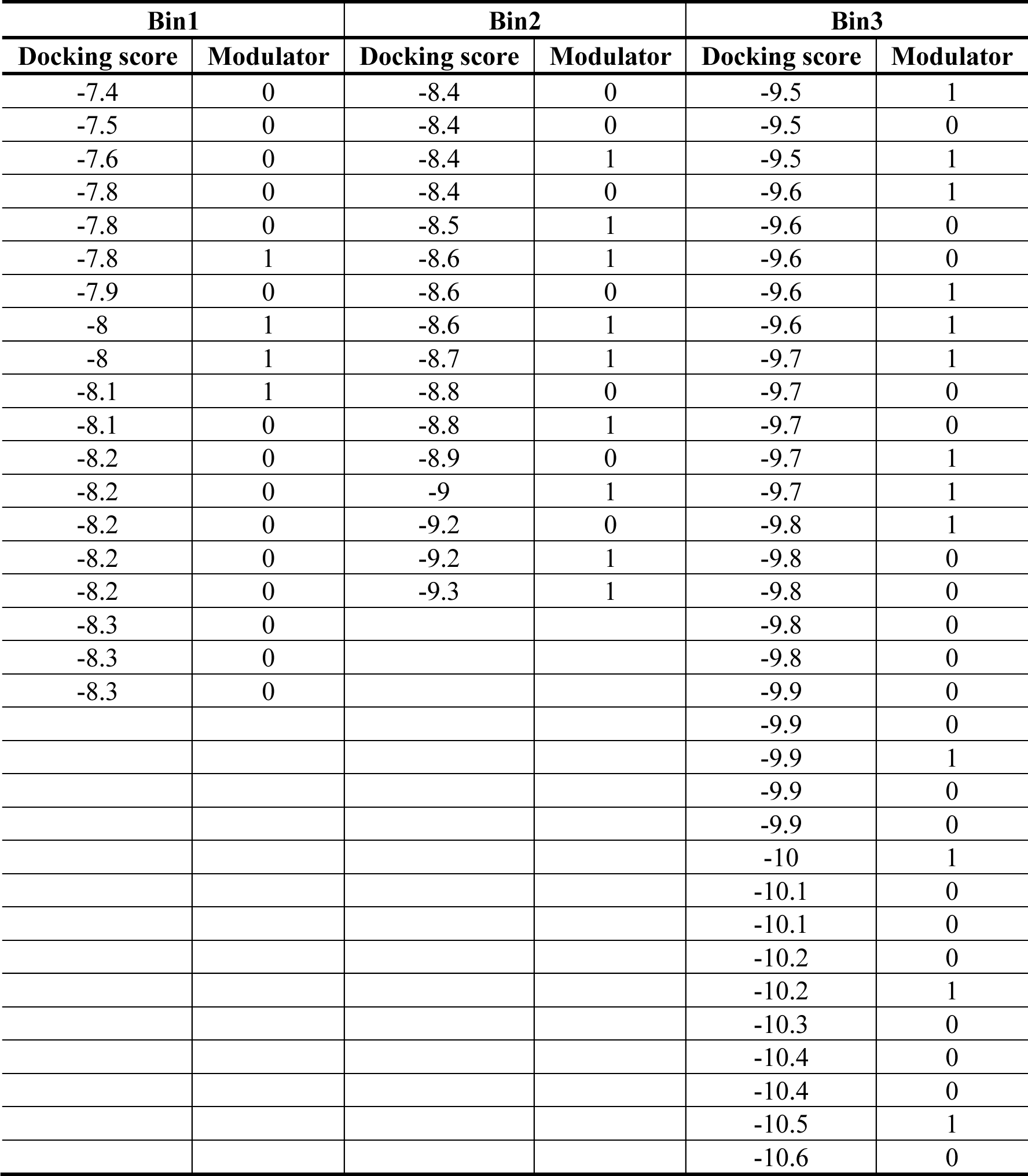
The distribution of bin sampling based on the docking scores.

**Table S6.** Model parameter estimates from global nonlinear fitting of mixed inhibition for SIRT3-FDL in the presence and absence of HKL. [E_0_] = 3.05 µM

|  | **0 $\mu\text{M}$ HKL** | **200 $\mu\text{M}$ HKL** |
| --- | --- | --- |
| $V_{max}$, $\mu\text{M}/\text{min}$ | $0.3923 \pm 0.0150$ | $0.08347 \pm 0.00220$ |
| $\alpha$ | $4.007 \pm 2.111$ | $\sim 3.727\text{e}+107$ |
| $K_i$, $\mu\text{M}$ | $24.52 \pm 4.15$ | $18.14 \pm 3.39$ |
| $K_m$, $\mu\text{M}$ | $1214 \pm 108$ | $846 \pm 62$ |
| **Goodness of Fit** |  |  |
| Degrees of Freedom | 8 | 8 |
| R square | 0.9961 | 0.996 |
| Absolute Sum of Squares | 0.0002103 | 0.00001222 |
| Sy.x | 0.005127 | 0.001236 |

### Supplementary Methods

#### Biophysical characterization of SIRT3 activation by HKL and hit compound - Measurement of binding affinities

In addition to measuring the effect of HKL on NAD^+^ binding affinity, we measured its effect on acetylated peptide binding affinity, NAM binding affinity in the presence of acetylated peptide, deacetylated peptide binding affinity, O-Acetylated ADP Ribose binding affinity in the presence of deacetylated peptide (i.e., the product complex), and NAD binding affinity in the presence of deacetylated peptide (**Figs. S8,** and **S9**).

#### Substrate dependence of SIRT3 modulation by HKL

The p53-AMC substrate was studied in order to explore the substrate dependence of modulation by HKL. It was verified that HKL binds to catalytically active complexes (**Figs. S2C** and **D**) and that there is a slight *k_cat_* efficiency decrease in the presence of modulator (**Table S6**). MST measurements on Carba-NAD^+^ indicate a decrease in NAD^+^ binding affinity (**Fig. S10B**). Non-steady state activation by HKL in the presence of p53-AMC substrate was also studied by measurements of activity under higher enzyme concentration ([E]_0_/[NAD^+^]_0_ = 0.3). The results are reported in **Fig. S10C**.

#### Characterization of HKL modulation of SIRT3 using computational, steady state kinetic, and thermodynamic data

**Fig. S3B** provides the expressions for each of the modulated steady state constants in **Fig. S3A** in the presence of a specified concentration [A] of the hit compound, according to the mechanism-based enzyme activation theory presented in (*1*). The *K*_*di,A*_in **Fig. S3B** denote the dissociation constants of the hit compound with respect to the respective reaction intermediates indexed by *i* in the model schematic **Fig. S11**.

**Fig. S5** depicts the changes to the steady state, Michaelis and dissociation constants in the sirtuin reaction mechanism when such a modulator is bound to the enzyme, as predicted by the model. Within this model, the conditions for activation were described in (*1*). These conditions establish the biophysical properties of the compound that are required to elicit changes in the steady state constants that are conducive to activation. A hit compound may be defined as one that satisfies a subset of the steady state conditions enumerated and also displays activation under non-steady state conditions, which can reduce inhibitory effects of the compound that are observed under steady state conditions, as discussed further below.

### Supplementary Discussion

#### Characterization of SIRT3 modulation by the non-steady state activator Honokiol

According to **Fig. S8B** (also **Table 3**), *K*_*d*,*NAD*^+^_ is roughly unchanged or slightly reduced for this substrate, implying that *K*_*d*1,*A*_ ≈ *K*_*d*2,*A*_ in **Fig. S11**. In the traditional picture, based on this data HKL binding would be characterized as noncompetitive with respect to NAD^+^. According to **Fig. S8D**, the increase in HKL binding affinity in the presence of NAM suggests that *K*_*d*,*NAM*_ decreases for this substrate, implying that *K*_*d*3,*A*_ > *K*_*d*4,*A*_ in **Fig. S11**. In the traditional picture of enzyme activity modulation, HKL binding is uncompetitive with respect to NAM.

However, the effect of HKL on activity can only be understood by application of the full kinetic and thermodynamic model described above, which involves simultaneous effects of the modulator on both the forward and reverse reactions in the context of a steady state model of the reaction. Note that in the presence of HKL, *K*_*m*,*NAD*^+^_ increases and steady state catalytic efficiency decreases by a somewhat larger margin. The parameter estimates in **Table 2** together with the full set of MST binding affinity data can be interpreted within the context of the model above to shed light on the mechanism of HKL modulation and to evaluate its properties as a hit compound.

The aim of steady state characterization of hit compounds for mechanism-based activation is to estimate the parameters in **Eq. (1)** in the absence of modulator and at a saturating concentration of modulator, varying both [NAD^+^] and [NAM] in each case. This provides estimates of both the front and back face steady state parameters in the model schematic **Fig. S11A**. By contrast, a mixed inhibition model with respect to the modulator concentration (as, e.g., applied to certain mechanism-based inhibitors (*2*)) would not have an interpretation for the steady state constants in terms of fundamental rate constants in the sirtuin mechanism, whereas characterizing unsaturating modulator concentrations at each of multiple product (NAM) concentrations, while providing more information, would similarly estimate only apparent steady state parameters depicted in **Fig. S11B**.

First, note from **Table 2** that *K_1_* decreases several fold in the presence of HKL, consistent with the increase in NAM binding affinity suggested by MST measurements. As discussed in (*1*), a decrease in *K_1_* may be inconsequential in terms of a molecule’s propensity to increase the enzyme’s catalytic efficiency as long as the common assumption of fast NAM release (fast *k_-3_* approximation) holds.

Second, *K_3_* decreases several fold in the presence of HKL (**Table 2**), whereas an increase in *K_3_* is desirable. However, *K_1_/K_3_* remains roughly unchanged, as can be observed graphically in **Figs. S6B** and **S6D** where the effect of *K_1_* becomes diminished at high [NAM]. This indicates that the reduction in *K_d,NAM_* may be playing an important role in the observed reduction of *K_3_*. Under the approximations discussed in (*1*), the modulated *K_ex_’* may be relatively close to *K_ex_* (see **Fig. S11** for the relationship between *K_3_* and *K_ex_*). This is a favorable property of a hit compound since it would imply *K*_*d*3,*A*_ ≈ *K*_*d*2,*A*_ ; hit evolution could be carried out to reduce *K_d3, A_*, which is conducive to activation. Note that since *K*_*d*,*NAD*^+^_ remains roughly unchanged by the modulator, if *K_ex_* is also roughly unchanged, any reduction in k_2_ that affects *K_m_* (see **Eq. (2)**) is associated with a similar reduction in *k_-2_*.

Third, α decreases several fold in the presence of HKL (**Table 2** and **Eq. (3)** for α). This shifts the intersection point of the lines in **Fig. S7A** to the left in **Fig. S7B** (see **Fig. S5** for a graphical depiction of *⍺* ∗ *K*_*m*,*NAD*^+^_ in terms of the intersection point). Under the above hypotheses, this is expected according to the expressions for *K_2,app_* and *K_3,app_* in **Fig. S3B**. Applying the approximations described in (*1*), this decrease in α is due to the fact that *K_m_* increases in the presence of HKL while *K*_*d*,*NAD*^+^_ does not increase / decreases slightly (see **Eq. (3)**). MST results on HKL’s effect on *K*_*d*,*NAD*^+^_ (**Fig. S10B**) are consistent with the reduction in *⍺* ∗ *K*_*m*,*NAD*^+^_ obtained from the steady state parameters in **Table 2**. According to **Eq. (3)**, *⍺* ∗ *K*_*m*,*NAD*^+^_ is highly correlated with *K*_*d*,*NAD*^+^_ and the methods are complementary, since MST employs a chemically modified NAD^+^ analog. Moreover, a slight decrease in *K*_*d*,*NAD*^+^_ but increase in *K*_*m*,*NAD*^+^_ can explain the observed non-steady state activation in the case of this substrate. We can also observe by comparing **Figs. S6 (c), (d)** to **(a), (b)** that in the presence of NAM, the extent of steady state inhibition by HKL diminishes, and that due to the reduction in α, this effect is most pronounced at low [NAD^+^], which is the condition under which activation is therapeutically most useful. Under the above hypotheses, through further hit-to-lead evolution of HKL, this property might be exploited to activate the enzyme under steady state conditions at low [NAD^+^] in the presence of physiological [NAM] or, going further, to allow a *K_m_* decrease in the absence of product, as was demonstrated for the novel steady state activators discovered in this work.

We can also understand the properties of conventional steady state inhibition plots with respect to modulator concentration based on the mechanistic information gleaned from our studies. Note from **Fig. S3A**, the intersection point of the lines with and without modulator lies to the left of the y axis and above the x axis. In the traditional mixed inhibition picture of steady state enzyme modulation (which does not distinguish between *K_m_* and *K_d_*), this is due to the modulator both increasing *K_m_* and decreasing *v_max_*, but this traditional picture does not distinguish between *K_m_* and *K_d_*, and also does not account for the relation between *K_m_* and *v_max_* (see **Eq. (2)** and **Fig. S3A**). By contrast, the equations for the lines in the presence and absence of modulator can be understood in terms of the fundamental dissociation constants and rate constants of the reaction based on our mechanistic analysis above.

Note also from **Fig. S6C**, the intersection point of the lines with and without modulator moves closer to the x axis in the presence of NAM. For sufficiently high [NAM], the intersection point of these lines falls below the x axis and then will eventually move to the right of the y axis. The fact that HKL does not increase *K*_*d*,*NAD*^+^_ gives it higher relative activity at low [NAD^+^] in the presence of NAM (this is especially pronounced in the pre-steady state phase, see inset in **Fig. S7 (d)**). If this could be achieved in the absence of NAM, it would signify an activator that increases steady state catalytic efficiency while reducing *v_max_* (as was achieved by some of the steady state activators identified in this work).

Turning to the p53-AMC substrate, in **Table S6**, which compares the initial rates of catalysis with this substrate in the presence and absence of saturating HKL, we observe a change in *v_max_* but not *K_m_* -- which would be characterized as noncompetitive inhibition in the traditional picture. However, as discussed above for the MnSOD substrate, the traditional picture is not sufficiently informative and the mixed inhibition plots with respect to NAM provide the important mechanistic information. In the absence of NAM, there is a 3-4 fold decrease in steady state catalytic efficiency for this substrate. However, whereas α decreased in the presence of HKL for the MnSOD substrate, it increases for the p53-AMC substrate. Since for MnSOD substrate, there was no increase in *K*_*d*,*NAD*^+^_ and a reduction in *⍺* ∗ *K*_*m*,*NAD*^+^_ induced by HKL, the increase in *⍺* ∗ *K*_*m*,*NAD*^+^_ induced by HKL for p53-AMC substrate observed in **Table S6** is consistent with the increase in *K*_*d*,*NAD*^+^_ as measured by MST (**Fig. S10** and **Table 4**). Taken together with the significant pre-steady state activation induced by HKL for this substrate (**Fig. 8C**), this also demonstrates that reduction in *K*_*d*,*NAD*^+^_ is not the only means for activation by HKL, consistent with the theory of mechanism-based activation.

In summary, HKL binds to all complexes in the sirtuin reaction mechanism, not just the product as in the case of mechanism-based sirtuin inhibitors like Ex-527. The tight binding of HKL to the coproduct does not reduce catalytic efficiency. Compounds like Ex-527 do not bind to any other catalytically relevant complexes and because they reduce product dissociation rate significantly, thus extinguishing the reaction under saturating conditions. Comparing the effects of saturating HKL and saturating Ex-527 (*3*) on SIRT3 activity, due to the substantial reduction in coproduct dissociation rate (steady state *k_cat_* close to 0), saturating Ex-527 effectively constitutes “single hit” conditions wherein each enzyme can only turn over products once. Ex-527 does this for many substrates and sirtuins including Ac-ACS2 and p53-AMC (*3*), due to the fact that it protrudes into the C pocket and is hence incapable of binding NAD^+^ in a productive conformation. **Fig. S11C** depicts the reaction diagram corresponding to the front face of **Fig. S11B** that includes separate representations of the product release and final chemistry (deacetylation) steps for modeling of the reaction under non-steady state conditions. (It can be shown that the addition of such irreversible rate constants does not alter the expressions for the steady state constants applied herein.) In the presence of HKL, *k_cat_* in the steady state model does not correspond to the rate determining step under non-steady state conditions; rather, being the rate limiting step of product release and the final chemistry step, it is approximately equal to the rate of product release. Since the rate determining step under non-steady state conditions is the final (deacetylation) chemistry step, the pre-steady state activation (**Fig. 8C**) of SIRT3 by HKL, as reported above, indicates that HKL reduces the rate of product release (by binding tightly to the enzyme·O-AADPR complex depicted in **Fig. S9**), but not necessarily the rate of the final chemistry step. Non-steady state (phases i and iii defined in the main text) measurements do not provide information on the effect of a modulator on product release, while measurements on the steady state system do not distinguish between effects the modulator has on the rate of product release and the rate of the final chemistry step. Thus, the two types of measurements play complementary roles in the characterization of hit compounds for mechanism-based activation.

Importantly, note that in the expression for the steady state catalytic efficiency *k*_*cat*_/*K*_*m*,*NAD*^+^_, which is relevant to steady state sirtuin activation under NAD^+^ depletion conditions associated with aging, the rate constants in *k_cat_* do not explicitly appear. Thus, the reduction in *k_cat_* by such a modulator does not affect catalytic efficiency. Hence the non-steady state activation by HKL is relevant to the goal of increasing catalytic efficiency, but further hit-to-lead evolution is required. These design goals have been achieved in the novel steady state activators discovered in the present work.

#### Structural similarity of top hit compounds

While searching for analogs of the top 9 assay-validated non-steady state activators from our 1.2 million compound docking study, with two SIRT3 receptor structures (4BVG and 4FVT), it became obvious that some hits have common structural features. While these common features are not sufficient for construction of a proper SAR model, they are adequate to influence choice of building blocks for combinatorial libraries. The most obvious recurring motif is a bicyclic ring (aromatic, partially aromatic or with partial delocalization) connected to another solitary aromatic ring via an amide (or urea) linker. This simple criterion can be applied when designing the scaffold of future ligands and combinatorial libraries.

## REFERENCES

1. M. Kaeberlein, M. McVey, L. Guarente, The SIR2/3/4 complex and SIR2 alone promote longevity in Saccharomyces cerevisiae by two different mechanisms. Genes & Development 13, 2570–2580 (1999).

2. A. A. Sauve, Sirtuin chemical mechanisms. Biochimica Et Biophysica Acta-Proteins and Proteomics 1804, 1591–1603 (2010).

3. P. Hu, S. Wang, Y. Zhang, Highly dissociative and concerted mechanism for the nicotinamide cleavage reaction in Sir2Tm enzyme suggested by ab initio QM/MM molecular dynamics simulations. Journal of the American Chemical Society 130, 16721–16728 (2008).

4. Y. Zhou et al., The bicyclic intermediate structure provides insights into the desuccinylation mechanism of human sirtuin 5 (SIRT5). The Journal of biological chemistry 287, 28307–28314 (2012).

5. M. D. Jackson, M. T. Schmidt, N. J. Oppenheimer, J. M. Denu, Mechanism of nicotinamide inhibition and transglycosidation by Sir2 histone/protein deacetylases. The Journal of biological chemistry 278, 50985–50998 (2003).

6. B. C. Smith, J. M. Denu, Sir2 protein deacetylases: evidence for chemical intermediates and functions of a conserved histidine. Biochemistry 45, 272–282 (2006).

7. B. C. Smith, J. M. Denu, Sir2 deacetylases exhibit nucleophilic participation of acetyl-lysine in NAD^+^ cleavage. Journal of the American Chemical Society 129, 5802–5803 (2007).

8. E. M. Mercken et al., SRT2104 extends survival of male mice on a standard diet and preserves bone and muscle mass. Aging Cell 13, 787–796 (2014).

9. S. J. Mitchell et al., The SIRT1 activator SRT1720 extends lifespan and improves health of mice fed a standard diet. Cell reports 6, 836–843 (2014).

10. J. A. Zorn, J. A. Wells, Turning enzymes ON with small molecules. Nature chemical biology 6, 179–188 (2010).

11. D. A. Sinclair, L. Guarente, Small-Molecule Allosteric Activators of Sirtuins. Annual review of pharmacology and toxicology 54, 363–380 (2014).

12. K. G. Tanner, J. Landry, R. Sternglanz, J. M. Denu, Silent information regulator 2 family of NAD-dependent histone/protein deacetylases generates a unique product, 1-O-acetyl-ADP-ribose. Proc Natl Acad Sci U S A 97, 14178–14182 (2000).

13. A. D. Napper et al., Discovery of indoles as potent and selective inhibitors of the deacetylase SIRT1. Journal of Medicinal Chemistry 48, 8045–8054 (2005).

14. B. P. Hubbard et al., Evidence for a Common Mechanism of SIRT1 Regulation by Allosteric Activators. Science 339, 1216–1219 (2013).

15. H. Dai et al., Crystallographic structure of a small molecule SIRT1 activator-enzyme complex. Nature Communications 6, (2015).

16. M. Lakshminarasimhan, D. Rauh, M. Schutkowski, C. Steegborn, Sirt1 activation by resveratrol is substrate sequence-selective. Aging-Us 5, 151–154 (2013).

17. K. E. Dittenhafer-Reed, J. L. Feldman, J. M. Denu, Catalysis and mechanistic insights into sirtuin activation. Chembiochem : a European journal of chemical biology 12, 281–289 (2011).

18. M. T. Borra, B. C. Smith, J. M. Denu, Mechanism of human SIRT1 activation by resveratrol. Journal of Biological Chemistry 280, 17187–17195 (2005).

19. X. A. Cambronne et al., Biosensor reveals multiple sources for mitochondrial NAD⁺. Science 352, 1474–1477 (2016).

20. C. Canto et al., The NAD(+) precursor nicotinamide riboside enhances oxidative metabolism and protects against high-fat diet-induced obesity. Cell Metab 15, 838–847 (2012).

21. T. L. Yang, A. A. Sauve, NAD metabolism and sirtuins: Metabolic regulation of protein deacetylation in stress and toxicity. Aaps Journal 8, E632–E643 (2006).

22. Q. Hu et al., Genetically encoded biosensors for evaluating NAD^+^/NADH ratio in cytosolic and mitochondrial compartments. Cell Reports Methods 1, 100116 (2021).

23. A. Mai et al., Study of 1,4-Dihydropyridine Structural Scaffold: Discovery of Novel Sirtuin Activators and Inhibitors. Journal of Medicinal Chemistry 52, 5496–5504 (2009).

24. V. B. Pillai et al., Honokiol blocks and reverses cardiac hypertrophy in mice by activating mitochondrial Sirt3. Nat Commun 6, 6656 (2015).

25. S. Valente et al., 1,4-Dihydropyridines Active on the SIRT1/AMPK Pathway Ameliorate Skin Repair and Mitochondrial Function and Exhibit Inhibition of Proliferation in Cancer Cells. Journal of Medicinal Chemistry 59, 1471–1491 (2016).

26. W. You et al., Structural Basis of Sirtuin 6 Activation by Synthetic Small Molecules. Angewandte Chemie (International ed. in English) 56, 1007–1011 (2017).

27. J. L. Feldman, J. Baeza, J. M. Denu, Activation of the protein deacetylase SIRT6 by long-chain fatty acids and widespread deacylation by mammalian sirtuins. The Journal of biological chemistry 288, 31350–31356 (2013).

28. J. L. Feldman et al., Kinetic and Structural Basis for Acyl-Group Selectivity and NAD^+^ Dependence in Sirtuin-Catalyzed Deacylation. Biochemistry 54, 3037–3050 (2015).

29. X. Guan, A. Upadhyay, S. Munshi, R. Chakrabarti, Biophysical characterization of hit compounds for mechanism-based enzyme activation. PLOS ONE 13, e0194175 (2018).

30. X. Guan, A. Upadhyay, R. Chakrabarti, Mechanism-based enzyme activating compounds. bioRxiv, 2020.2004.2008.032235 (2020).

31. C. Reverdy et al., Discovery of novel compounds as potent activators of Sirt3. Bioorganic & medicinal chemistry 73, 116999 (2022).

32. A. Ansari et al., Function of the SIRT3 mitochondrial deacetylase in cellular physiology, cancer, and neurodegenerative disease. Aging cell 16, 4–16 (2017).

33. R. A. H. van de Ven, D. Santos, M. C. Haigis, Mitochondrial Sirtuins and Molecular Mechanisms of Aging. Trends in molecular medicine 23, 320–331 (2017).

34. W. You et al., Structural Basis of Sirtuin 6 Activation by Synthetic Small Molecules. Angewandte Chemie International Edition 56, 1007–1011 (2017).

35. J. L. Feldman, J. Baeza, J. M. Denu, Activation of the protein deacetylase SIRT6 by long-chain fatty acids and widespread deacylation by mammalian sirtuins. The Journal of biological chemistry 288, 31350–31356 (2013).

36. B.-H. Ahn et al., A role for the mitochondrial deacetylase Sirt3 in regulating energy homeostasis. Proceedings of the National Academy of Sciences of the United States of America 105, 14447–14452 (2008).

37. W. Anderson, Structural Genomics and Drug Discovery: Methods and Protocols. (2014), vol. 1140.

38. J. Eberhardt, D. Santos-Martins, A. F. Tillack, S. Forli, AutoDock Vina 1.2.0: New Docking Methods, Expanded Force Field, and Python Bindings. Journal of Chemical Information and Modeling 61, 3891–3898 (2021).

39. O. Trott, A. J. Olson, AutoDock Vina: improving the speed and accuracy of docking with a new scoring function, efficient optimization, and multithreading. Journal of computational chemistry 31, 455–461 (2010).

40. S. Vilar, G. Cozza, S. Moro, Medicinal chemistry and the molecular operating environment (MOE): application of QSAR and molecular docking to drug discovery. Current topics in medicinal chemistry 8, 1555–1572 (2008).

41. J. Gallego-Jara, A. Écija Conesa, T. de Diego Puente, G. Lozano Terol, M. Cánovas Díaz, Characterization of CobB kinetics and inhibition by nicotinamide. PLOS ONE 12, e0189689 (2017).

42. W. Dang, The controversial world of sirtuins. Drug discovery today. Technologies 12, e9–e17 (2014).

43. J. L. Avalos, J. D. Boeke, C. Wolberger, Structural basis for the mechanism and regulation of Sir2 enzymes. Molecular Cell 13, 639–648 (2004).

44. X. Guan, P. Lin, E. Knoll, R. Chakrabarti, Mechanism of inhibition of the human sirtuin enzyme SIRT3 by nicotinamide: computational and experimental studies. PLoS One 9, e107729 (2014).

45. N. Xie et al., NAD^+^ metabolism: pathophysiologic mechanisms and therapeutic potential. Signal Transduction and Targeted Therapy 5, 227 (2020).

46. M. R. McReynolds, K. Chellappa, J. A. Baur, Age-related NAD(+) decline. Experimental gerontology 134, 110888 (2020).

47. Y. Cao et al., Quercetin promotes in vitro maturation of oocytes from humans and aged mice. Cell Death & Disease 11, 965 (2020).

48. R. Chakrabarti, A. M. Klibanov, R. A. Friesner, Sequence optimization and designability of enzyme active sites. Proc Natl Acad Sci U S A 102, 12035–12040 (2005).

49. The CCP4 suite: programs for protein crystallography. Acta crystallographica. Section D, Biological crystallography 50, 760–763 (1994).

50. P. Emsley, K. Cowtan, Coot: model-building tools for molecular graphics. Acta crystallographica. Section D, Biological crystallography 60, 2126–2132 (2004).

51. S. M. Pandey, A.; Phale, P.S.; Bhaumik, P., Cloning, Purification, Crystallization and Preliminary X-Ray Diffraction Studies of Periplasmic Glucose Binding Protein of Pseudomonas putida CSV86. Advances in Bioscience and Biotechnology 6, 8 (2015).

52. P. Chys, P. Chacón, Random Coordinate Descent with Spinor-matrices and Geometric Filters for Efficient Loop Closure. J Chem Theory Comput 9, 1821–1829 (2013).

53. J. R. López-Blanco, A. J. Canosa-Valls, Y. Li, P. Chacón, RCD+: Fast loop modeling server. Nucleic Acids Res 44, W395–400 (2016).

54. W. Elhefnawy, L. Chen, Y. Han, Y. Li, ICOSA: A Distance-Dependent, Orientation-Specific Coarse-Grained Contact Potential for Protein Structure Modeling. Journal of molecular biology 427, 2562–2576 (2015).

55. V. Hornak et al., Comparison of multiple Amber force fields and development of improved protein backbone parameters. Proteins 65, 712–725 (2006).

56. J. Wang, P. Cieplak, P. A. Kollman, How well does a restrained electrostatic potential (RESP) model perform in calculating conformational energies of organic and biological molecules? Journal of Computational Chemistry 21, 1049–1074 (2000).

57. Q. Lu, Y.-H. Tan, R. Luo, Molecular dynamics simulations of p53 DNA-binding domain. The journal of physical chemistry. B 111, 11538–11545 (2007).

58. R. C. Walker, M. M. de Souza, I. P. Mercer, I. R. Gould, D. R. Klug, Large and Fast Relaxations inside a Protein: Calculation and Measurement of Reorganization Energies in Alcohol Dehydrogenase. The Journal of Physical Chemistry B 106, 11658–11665 (2002).

59. J. J. Pavelites, J. Gao, P. A. Bash, A. D. Mackerell Jr., A molecular mechanics force field for NAD^+^ NADH, and the pyrophosphate groups of nucleotides. Journal of Computational Chemistry 18, 221–239 (1997).

60. G. V. Papamokos et al., Structural role of RKS motifs in chromatin interactions: a molecular dynamics study of HP1 bound to a variably modified histone tail. Biophysical journal 102, 1926–1933 (2012).

61. T. Darden, D. York, L. Pedersen, Particle mesh Ewald: An N⋅log(N) method for Ewald sums in large systems. The Journal of chemical physics 98, 10089–10092 (1993).

62. J. C. Phillips et al., Scalable molecular dynamics with NAMD. Journal of computational chemistry 26, 1781–1802 (2005).

63. J. P. Ryckaert, G. Ciccotti, H. J. C. Berendsen, Numerical integration of the Cartesian Equations of Motion of a System with Constraints: Molecular Dynamics of n-Alkanes. Journal of Computational Physics 23, 327–341 (1977).

64. B. G. Szczepankiewicz et al., Synthesis of carba-NAD and the structures of its ternary complexes with SIRT3 and SIRT5. The Journal of organic chemistry 77, 7319–7329 (2012).

65. M. Gertz et al., Ex-527 inhibits Sirtuins by exploiting their unique NAD(+)-dependent deacetylation mechanism. Proceedings of the National Academy of Sciences of the United States of America 110, E2772–E2781 (2013).

66. M. Feher, E. Deretey, S. Roy, BHB: A Simple Knowledge-Based Scoring Function to Improve the Efficiency of Database Screening. Journal of Chemical Information and Computer Sciences 43, 1316–1327 (2003).

67. P. Emsley, B. Lohkamp, W. G. Scott, K. Cowtan, Features and development of Coot. Acta crystallographica. Section D, Biological crystallography 66, 486–501 (2010).

68. A. T. Brünger, Free R value: a novel statistical quantity for assessing the accuracy of crystal structures. Nature 355, 472–475 (1992).

69. K. Moffat, in Methods in enzymology. (Academic Press, 1997), vol. 277, pp. 433–447.

70. I. J. Tickle, Statistical quality indicators for electron-density maps. Acta crystallographica. Section D, Biological crystallography 68, 454–467 (2012).

71. M. D. Winn et al., Overview of the CCP4 suite and current developments. Acta crystallographica. Section D, Biological crystallography 67, 235–242 (2011).

## Supplementary References

1. X. Guan, A. Upadhyay, S. Munshi, R. Chakrabarti, Biophysical Characterization of Hit Compounds for Mechanism-based Enzyme Activation. PLOS One 13, e0194175 (2018).

2. T. Rumpf et al., Selective Sirt2 inhibition by ligand-induced rearrangement of the active Textte. Nat Commun 6, 6263 (2015).

3. M. Gertz et al., Ex-527 inhibits Sirtuins by exploiting their unique NAD(+)-dependent deacetylation mechanism. Proceedings of the National Academy of Sciences of the United States of America 110, E2772–E2781 (2013).

